# Novel regulators of growth identified in the evolution of fin proportion in flying fish

**DOI:** 10.1101/2021.03.05.434157

**Authors:** Jacob M. Daane, Nicola Blum, Jennifer Lanni, Helena Boldt, M. Kathryn Iovine, Charles W. Higdon, Stephen L. Johnson, Nathan R. Lovejoy, Matthew P. Harris

**Author notes:** authors for communication (Jacob M. Daane, Matthew P. Harris). posthumous work (Stephen L. Johnson).

## Abstract

Identifying the genetic foundations of trait variation and evolution is challenging as it is often difficult to parse meaningful signals from confounding signatures such as drift and epistasis. However, identification of the genetic loci underlying morphological and physiological traits can be honed through the use of comparative and complementary genetic approaches, whereby shared sets of genes that are repeatedly implicated across large evolutionary time periods as under selection can illuminate important pathways and epistatic relationships that function as novel regulators of trait development. Here we intersect comparative genomic analyses with unbiased mutagenesis screens in distantly related species to define the control of proportional growth, as changes in the size and relative proportions of tissues underlie a large degree of the variant forms seen in nature. Through a phylogenomic analysis of genome-wide variation in 35 species of flying fishes and relatives, we identify genetic signatures in both coding and regulatory regions underlying the convergent evolution of increased paired fin size and aerial gliding behaviors, key innovations for flying fishes and flying halfbeaks. To refine our analysis, we intersected convergent phylogenomic signatures with mutants identified in distantly related zebrafish with altered fin size. Through these paired approaches, we identify a surprising role for an L-type amino acid transporter, *lat4a*, and the potassium channel, *kcnh2a*, in the regulation of fin proportion. We show that specific epistatic interaction between these genetic loci in zebrafish closely phenocopies the observed fin proportions of flying fishes. The congruence of experimental and phylogenomic findings point to a conserved, non-canonical signaling interaction that integrates bioelectric cues and amino acid transport in the establishment of relative size in development and evolution.

## Introduction

Changes to allometry, or the relative proportions of organs and tissues within organisms, is a common means for adaptive character change in evolution. However, little is understood about how relative size is specified during development and shaped during evolution. Actinopterygian (ray-finned) fishes provide rich opportunities to study variation in proportion, as they are the most species-rich and diverse class of vertebrates with over 30,000 species [1]. The actinopterygian fin exhibits dramatic changes in size and shape that not only impacts movement and balance, but has been modified into more elaborate innovations, including lures, defensive spines and sucking disks. Not surprisingly, in many actinopterygian species, modification of the size and shape of the fin is a defining and ecologically important trait.

Several actinopterygian lineages have independently evolved elongated, wing-like fins that enable gliding through the air [2]. Among the most accomplished aerial gliders are members of the family Exocoetidae (Beloniformes), or the “flying fishes”. All flying fish species have greatly enlarged pectoral fins as adults (**Fig 1A**) that they use as airfoils to glide through the air to avoid the aquatic predators of their open water epipelagic habitats that are largely devoid of shelter [2]. A subset of flying fishes, the four-winged gliders, also have elongated pelvic fins that provide additional aerodynamic support. These fishes have the most refined gliding characteristics of the group and can glide for up to 400 meters with controlled turns and altitude changes [2]. In combination with elongated paired fins, flying fishes have a hypocercal caudal fin with a stiffened and elongated ventral lobe (**Fig 1A**). The hypocercal caudal fins not only facilitate the initial emergence and take-off from the water, but can also provide thrust when the rest of the body is above the water level, allowing some species to generate additional momentum without fully re-entering the water [3].

**Fig 1.**
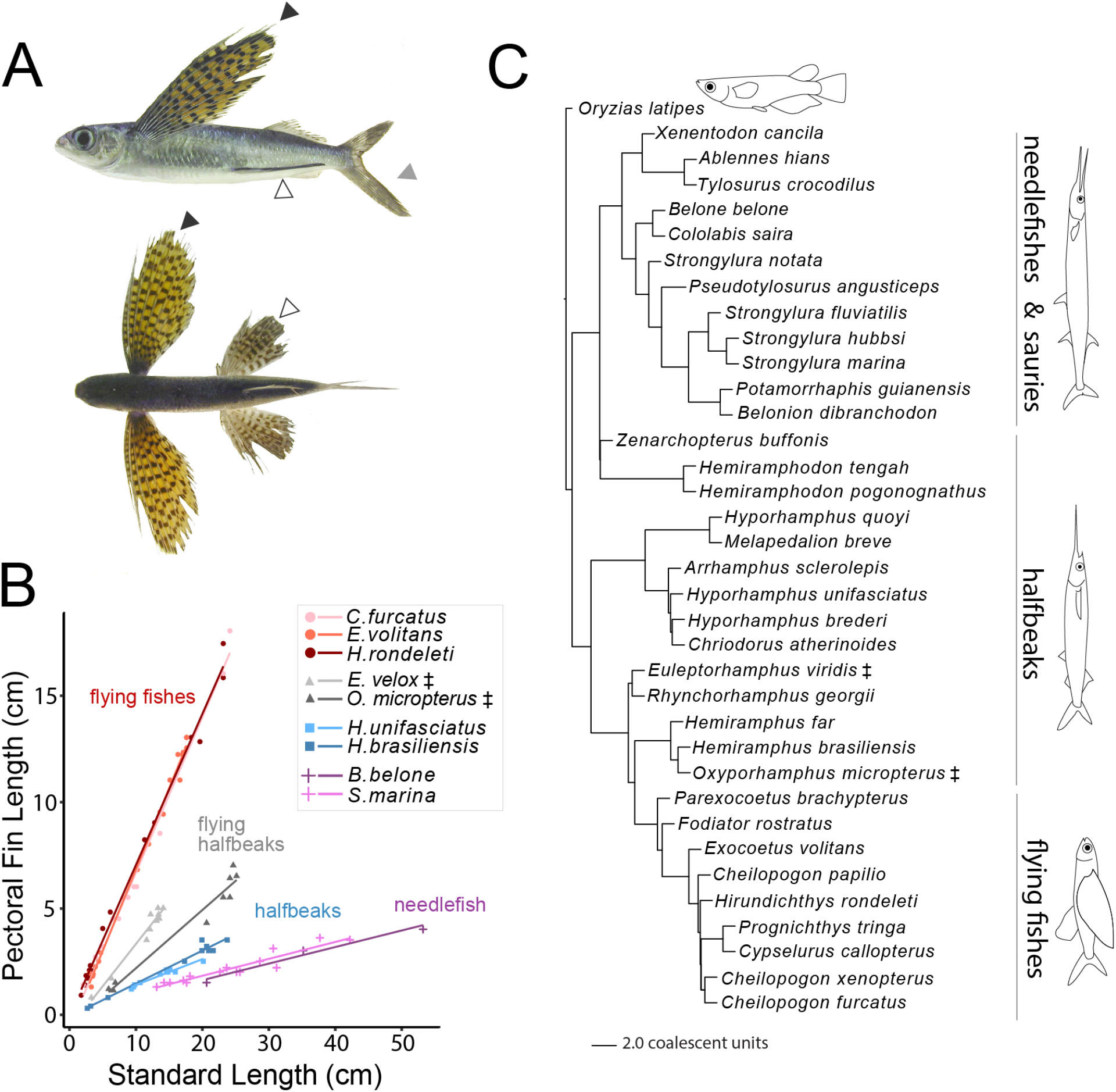
Evolution of fin allometry and gliding behavior in Beloniformes. (**A**) Lateral and dorsal views of a flying fish, *Cypselurus callopterus*, highlighting elongated pectoral (black arrowhead) and pelvic fins (white arrowhead) that act as an airfoil to enable aerial gliding behavior. The asymmetry of the caudal fin (hypocercal) aids in above-water propulsion (gray arrowhead). (**B**) Static allometry of pectoral fins in flying fishes (Exocoetidae; *C. furcatus, E. volitans, H. rondeleti*) compared to sauries and needlefish (*B. belone, S. marina*) and halfbeaks (*H. unifasciatus, H. brasiliensis*). The ‘flying halfbeaks’ (*E. velox, O. micropterus*) also exhibit aerial gliding behavior and have an intermediate pectoral fin length relative to the fishes of Exocoetidae. (**C**) Phylogeny of the beloniforms sequenced in this study. ‡ indicates lineages of flying halfbeak.

Flying fishes are not the only beloniforms with aerial gliding behavior, as several partial-gliding species occur within the closely related halfbeak family (Hemiramphidae, **Fig 1B,C**). “Flying halfbeaks” of the genera *Oxyporhamphus* and *Euleptorhamphus* have elongated pectoral fins and exhibit gliding behaviors, though these fishes fail to attain the degree of controlled, prolonged flight associated with Exocoetidae. Species of the flying halfbeak genus *Oxyporhamphus* have also lost the halfbeak jaw morphology, similarly to flying fishes [3–5].

Here, we used integrated genetic and comparative genomic approaches to investigate the shift in fin allometry in Beloniformes as an evolutionary model to identify regulators of growth and proportion. Though high-density taxonomic sampling of genetic variation across Beloniformes, we define unique genomic patterns associated with constellation of traits related to gliding flight. Through this ‘phylomapping’ approach, we define species-specific changes across the coding and non-coding genomic complement across the clade (**Fig 1C**, **Fig 2**) that highlight pathways and genes tied to the proportional growth and size of appendages. In parallel, we identify genetic causes underlying mutants found in experimental forward genetic screens in laboratory zebrafish. Through these methods, we demonstrate overlap in the genes and processes underlying overgrowth phenotypes even among these distantly related fishes and identify a novel function for amino acid transport in regulating size control. Through these paired approaches, we demonstrate sufficiency of the additive function of two loci to phenocopy patterns of fin proportionality of flying fish. These findings extend current understanding of growth regulation pathways and support a model of modulation of ionic or bioelectric signaling that potentiates fin allometry that is a key innovation for the group.

**Fig 2.**
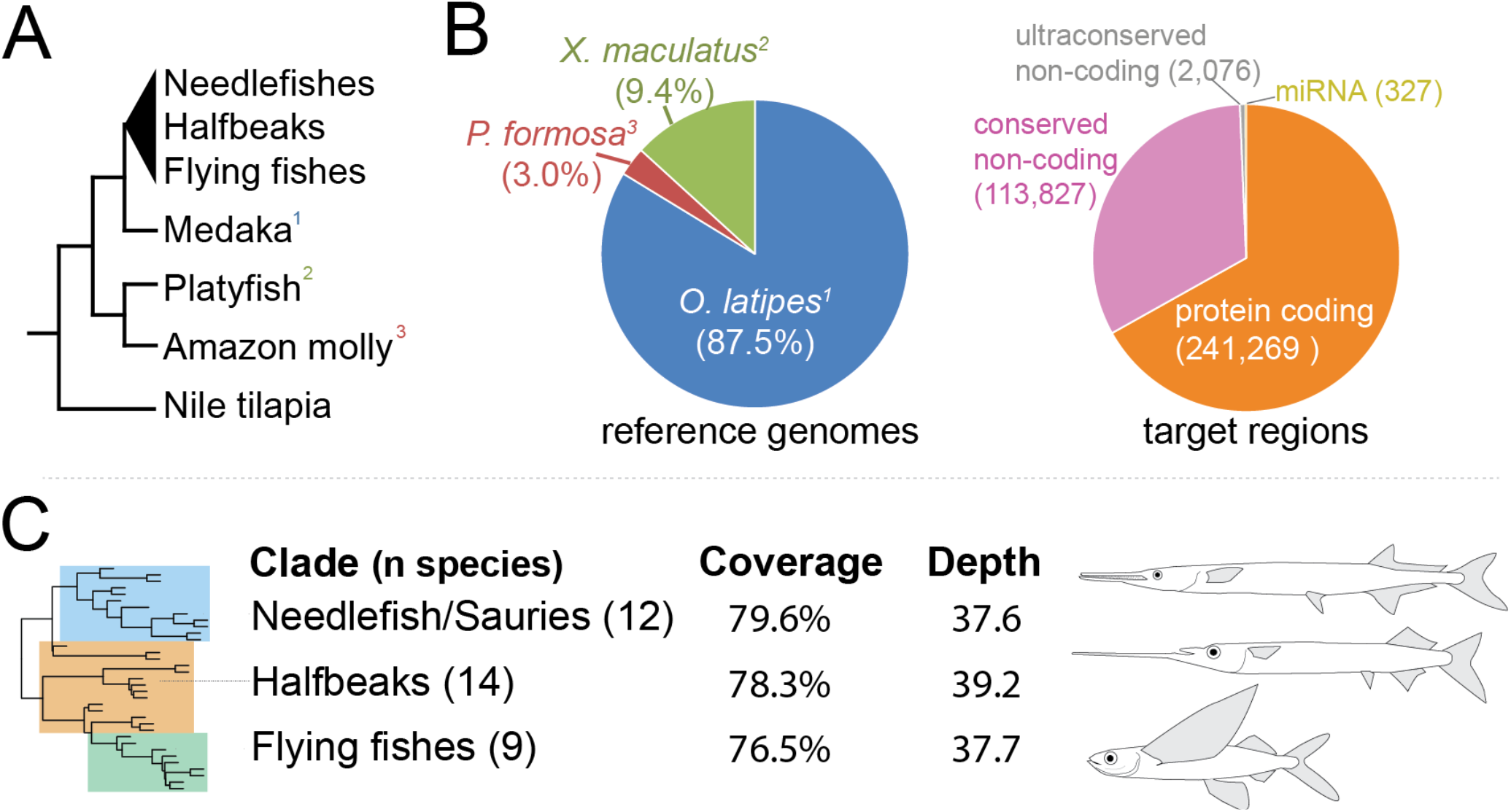
Beloniformes targeted sequence enrichment design and coverage. **(A)** Phylogenetic relationship of Beloniformes with the outgroup genomes used to design targeted sequencing baits. Numbers in phylogeny indicates the species in (**B). (B-C)** ‘Phylochip’ capture array design. **B)** The medaka genome (*O. latipes* HdrR, MEDAKA1) was used as foundation for the design of capture baits. To ensure broad coverage and avoid medaka-specific artifacts, regions from the platyfish (*X. maculatus*, Xipmac4.4.2) and Amazon molly (*P. formosa*, Poecilia_formosa-5.1.2) genomes that could not be identified within the *O. latipes* assembly were included in the capture design. (**C**) Number and breakdown of targeted genomic elements, including protein coding exons, conserved non-coding elements, ultraconservative elements and miRNA hairpins. (**D**) Sequencing read coverage and depth data from the different beloniform clades. Detailed coverage statistics and analysis are provided in **S1 Fig** and **S4-S6 Tables**.

## Results

### Evolutionary shaping of fin allometry in Beloniformes

The clear evolutionary and ecological significance of fin allometry in flying fish make this lineage an attractive case study to understand the regulation of scaling in evolution. To understand the changes in allometry of pectoral fins during beloniform evolution, we measured fin length as a function of body size (standard length) in archived samples from representative species across the clade (**Fig 1B**, **S1 Table**). While the halfbeaks and needlefishes show only subtle increases in pectoral fin length as the body grows, the length of flying fish pectoral fins increases rapidly (**Fig 1B**, **S1 Table**). The allometric relationship between body size and pectoral fin length was identical across the two and four-wing flying fishes we analyzed (**Fig 1B**) indicating that the regulation of fin proportion may be fixed across all flying fishes. Intriguingly, the two genera of gliding halfbeaks (*Euleptorhamphus* and *Oxyporhamphus*) that exhibit an increase in fin size that is convergent with flying fishes show a fin-body length allometric scaling relationship that is intermediate between halfbeaks and flying fishes (**Fig 1B,C** and **S1 Table**)[2].

### Identification of genetic variation across Beloniformes

As genetic analysis in Beloniformes is limited by the absence of sequenced genomes and the inability to assess linkage or association through hybrid analyses of natural populations, we mined the genomic variability within the clade through targeted sequencing of 35 species to look for genetic trends shared among flying fishes and gliding halfbeaks but not seen in related lineages. We sampled over 300,000 distinct genetic loci representing conserved non-coding sequences and protein coding genes. The closest relative species with a sequenced and annotated genome to the flying fishes is the ricefish, the Japanese medaka (*Oryzias latipes*). To facilitate genetic analysis of Beloniformes, we designed targeted capture baits based on predicted coding and non-coding sequence (microRNA and conserved non-coding elements (CNEs)) from the medaka as well as genomic data compiled from the platyfish (*Xiphophorus maculatus*) and Amazon molly (*Poecilia formosa*), members of the cyprinodontiform sister group to Beloniformes (**Fig 2A** and **2B**). This taxonomically diverse ‘phylochip’ enables recovery of targeted genomic regions that may be missing or highly divergent across the genomes of different species.

We extracted and pooled genomic DNA from several individuals from each of 35 different beloniform species, including species from the two-and four-winged flying fishes, halfbeaks, sauries, and needlefishes (**S2** and **S3 Tables**). Using relaxed hybridization of DNA fragments to our phylochip capture array, we were successful in enriching and sequencing genomic DNA covering an average of 78.3% of the targeted genomic regions at an average 38.3-fold depth (**Fig 2C, S1 Fig, S4** and **S5 Tables**). This recovery was impressive given, and despite, the large evolutionary distance between the species sequenced and the reference genomes used to design baits (>70 million years (My) [6–8]). We found comparable recovery among CNE, microRNA and coding sequence categories, providing a broad sampling of functionally annotated and conserved genetic regions across Beloniformes (**S4 Table**). This phylogenomic approach, termed ‘phylomapping’ [9], is empowered through the usage of dense sampling of taxa across the phylogeny, which permits the pinpointing of species-specific signals and high-resolution reconstruction of key ancestral sequences.

As with other cross-species targeted sequence capture experiments based on hybridization [9,10], there is enrichment within the set of poorly covered regions for genes that are rapidly evolving, particularly in gene clusters involving the immune system (**S6 Table**). Importantly, even though these gene classes are enriched for those genes with missing coverage compared to the rest of the genome, the majority of the captured genes (77.9%) within the GO-term categories that are enriched for missing coverage do not have low coverage (**S5 Table**). By sequencing pooled DNA samples from several individuals for each species (**S2 Table**), we were able to categorize fixed and heterozygous nucleotide variation for each species sample in the phylogeny (**S7 Table**). Among reconstructed exons, we found no evidence for genome-wide differences in copy number variation among species (**S8 Table**).

We estimated a phylogeny for the clade from 4,683 reconstructed gene trees in our dataset (**Fig 1D**). This tree is largely concordant with previously published phylogenies [3,5,11]. Notably, the resulting phylogeny supports the paraphyly of halfbeaks, with *Hermiramphus* and *Oxyporhamphus* the sister group to the monophyletic flying fishes [3]. In concordance with other recent molecular phylogenies of Beloniformes [3,5], we place *Oxyporhamphus*, a genus with elongated pectoral fins that also lacks the halfbeak jaw morphology as an adult, within the halfbeaks and not as a sister group to flying fishes as has been previously proposed [4].

### Genomic imprint of the evolution of aerial gliding in fishes

To assess genetic changes associated with gliding behavior and morphology, we analyzed patterns of evolutionary rate across genomes of gliding beloniform compared to non-gliding beloniform branches (**Fig 3**, **S9 Table**). Randomly-sampled controls representing species groupings with the same tip and ancestral node distributions as in the gliding lineages do not show statistical evidence for accelerated or constrained evolution across any gene groups (**S2 Fig**, **S9 Table**). However, in gliding branches we observed an elevated evolutionary rate in gene classes associated with the key morphological adaptations previously associated with gliding (**Fig 3**, **S9 Table**). Notably, accelerated gene groupings include fin development and morphogenesis, pectoral fin morphogenesis, embryonic digit morphogenesis, and fore-and hindlimb morphogenesis (**Fig 3**, **S9 Table**). The above-water taxiing and aerial gliding observed in gliding fishes is made possible by adaptations that affect balance and musculature as well as behavior. These morphological changes include enlarged semicircular canals [12], increased size of pectoral fin musculature and neuromuscular adaptations to enable propulsive tail oscillations at speeds up to 50 beats per second [2,13]. Accordingly, we observed an accelerated sequence evolution across a number of gene groupings relating to balance, including vestibular receptor cell development, semicircular canal formation and morphogenesis, adult locomotory behavior and directional locomotion, hindbrain development and cerebellar granule cell migration (**Fig 3**, **S9 Table**). Additionally, the genes across several muscle and heart-related terms are accelerated in gliders, including skeletal muscle hypertrophy, heart development and cardiac tissue morphogenesis (**Fig 3**, **S9 Table**). Gliding behaviors are thought to represent an anti-predator response [2]. Intriguingly, we find accelerated sequence evolution in the genes involved in locus cereleus development, a key region of the brain involved in alertness, anxiety and panic response in humans [14], as well as epinephrine secretion and transport and adrenergic receptor activity (**Fig 3**, **S9 Table**). Finally, flying fish have unique pyramidal corneas adapted to provide clear image resolution both above and below the water surface [15], and we observed accelerated evolutionary rate across a number of visual pathways, including camera-type eye morphogenesis, retinal development, lens development and morphogenesis, and lens fiber cell differentiation (**Fig 3**, **S9 Table**). Thus, a number of genes associated with key adaptations for gliding are under accelerated sequence evolution in gliding beloniforms.

**Fig 3.**
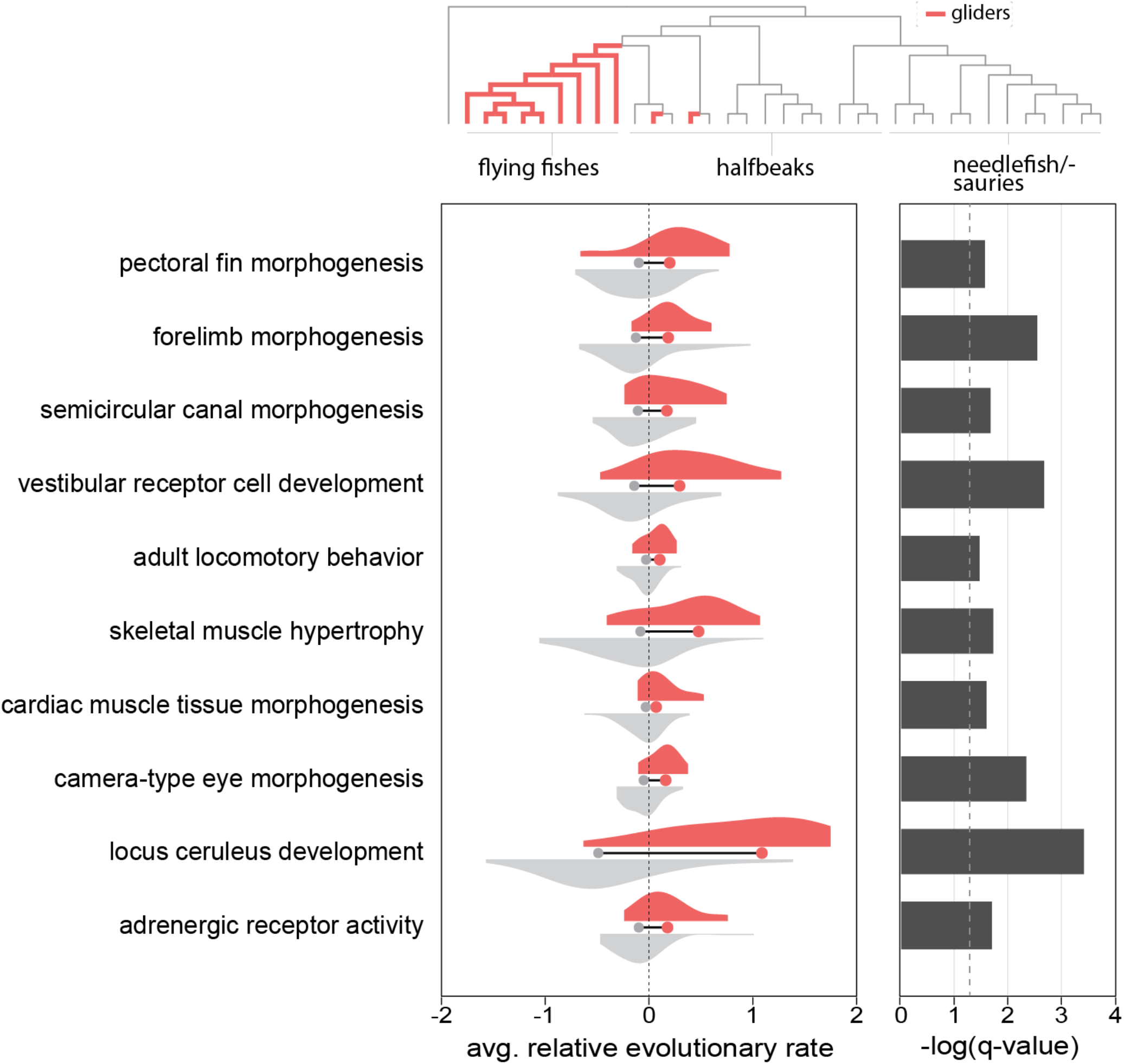
Accelerated sequence evolution of key genomic regions in gliding beloniforms. Comparison of average relative evolutionary rate between the gliding and non-gliding beloniform nodes across specific gene ontology terms across the beloniform phylogeny (gliding fish lineages highlighted in red). Histograms represent the distribution across each node in the phylogeny of the average relative evolutionary rate for all genes within a given gene ontology term. The mean of the average relative evolutionary rates in gliding and non-gliding beloniform nodes is denoted at the base of each histogram. Significance (-log(q-value)) based on FDR-corrected p-values from a Wilcoxon signed-rank test of differences between the mean relative evolutionary rate of gliding beloniforms compared to non-gliding beloniform branches. For full enrichment data, see **S9 Table**.

### Genetic basis of fin allometry

To explore the genetic basis of elongated paired fins in flying fishes, we intersected forward genetic and comparative genomic approaches. The genetic basis of fin proportion is actively investigated through genetic and developmental analyses in laboratory fish populations [16], and there is a growing body of information regarding genes and regulatory mechanisms controlling fin size. We leveraged an existing mutant collection in zebrafish to ask if mutant phenotype analysis could refine the analysis of natural genomic variation towards isolating new, potentially core, conserved developmental mechanisms. In this effort, we detail mapping of two mutants that affect genes that were previously not known to regulate fin growth. We then expand on these defined mechanisms in the pattern of genome evolution identified in lineages of gliding beloniforms.

#### The zebrafish longfin mutant and the regulation of fin proportion

The dominant *longfin^t2^* mutant (*lof^dt2^*) was first isolated in an aquarium population, and is one of the initial strains identified in zebrafish (*Tüpfel long fin*) [17]. The *lof* mutant has a coordinated overgrowth of all fins that occurs during post-embryonic development. The effect of the mutant allele on growth is dose sensitive such that homozygous fish have larger fins (**Fig 4A**). Importantly, *lof* mutants maintain elongated but regular segmentation of lepidotrichia (**S3 Fig A,B**), in contrast to the *kcnk5b^alf^* fin overgrowth mutant which has elongated and irregular segmentation patterning (**S3 Fig A,B**; ref [18]). As in the *lof* mutant, the segmentation pattern of flying fish lepidotrichia are regular (**S3 Fig C**).

**Fig 4.**
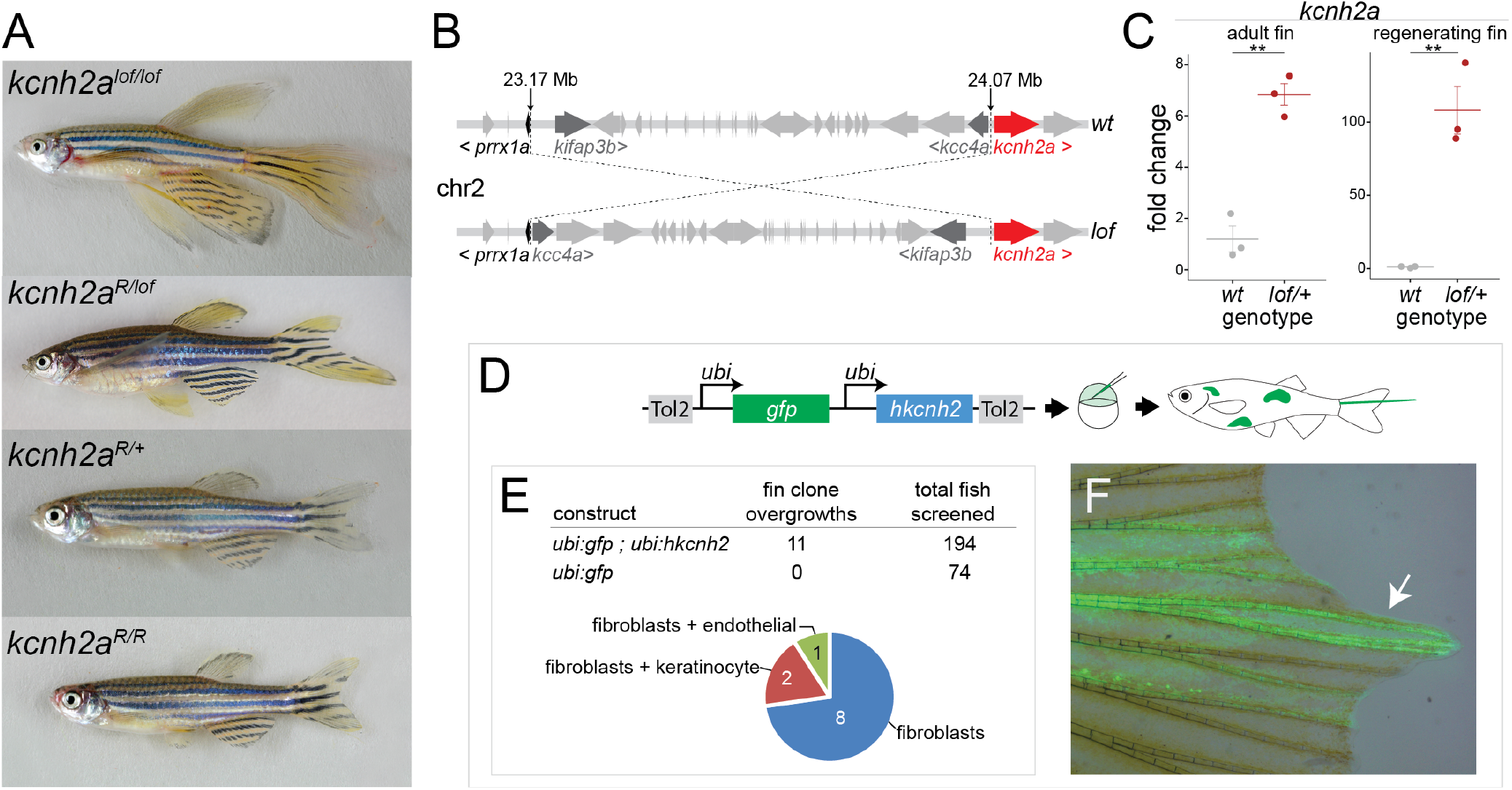
Identification of potassium channel *kcnh2a* underlying the *longfin^dt2^* zebrafish phenotype. **(A)** Revertant mutants of the zebrafish *longfin^dt2^ (lof*) mutant obtained from a mutagenesis screen. Reversion alleles mapped to loss-of-function mutations in the voltage-gated potassium channel *kcnh2a*. **(B)** Identified chromosomal inversion in *lof*. Note this inversion juxtaposes the regulatory region of *prrx1a* upstream of the *kcnh2a* transcription start site. For detailed positional mapping information see **S4 Fig** and **S11 Fig**. **(C)** qRT-PCR showing upregulation of *kcnh2a* in adult and regenerating caudal fins of *lof* compared to wild-type. **(D)** Mosaic overexpression assay to assess effect of hKCNH2 overexpression on fin growth. The plasmid is injected into single cell zebrafish embryos with Tol2 transposase mRNA and is incorporated randomly into the genome, resulting in a mosaic patchwork of gene overexpression in the adult. **(E)** Number of overgrowths observed in injected zebrafish and marked cells underlying the overgrowth. **(F)** Example fin overgrowth showing GFP+ fibroblast clones in within the overgrown fin rays (arrow).

We investigated the genetic causes of the *lof* phenotype. While it has been known for decades that the *lof* phenotype is caused by mutations on chromosome 2 [19], the specific molecular changes causing fin growth have eluded detection. Positional mapping showed limited ability to refine the mapping interval of *lof*. Using both ENU and γ-ray mutagenesis, we performed two independent reversion screens in mutagenized homozygous *lof* founders. In these independent analyses, we screened F1 progeny from wild-type outcrosses for reversion of the *lof* phenotype. Several mutant revertant lines were recovered from the ENU screen, each with independent nonsynonymous mutations within the gene encoding the potassium channel Kcnh2a (*lof*^*l*fr1^ Y669N, lof^lfr2^ L739Q, *lof*^WL4^, *lof*^WL7^ Y418X; **Fig 4A**, **S4 Fig**). Further, a single revertant was isolated from the γ-ray screen (*lof*^j6g1^) which disrupted much of the mapping interval, including removal of the sequence upstream of the *kcnh2a* transcription start site (**S4 Fig**) [19]. Homozygous mutants of *kcnh2a* revertants are viable and have normal proportion of wild-type fins, revealing this gene to be dispensable for normal development (**Fig 4A**, **S4 Fig**). The ability to revert the mutant phenotype by loss-of-function alleles of *kcnh2a in cis* to the *lof* allele indicates the mutant is due to a gain-of-function of the channel.

Analysis of long sequencing reads (PacBio), identified an inversion on chromosome 2 in *lof* with break points between 5’ *prrx-1a* and 5’ *knch2a* without affecting the coding region of either gene (**Fig 4B**, **S4 Fig**). This new position of Kcnh2a in *lof* suggests that the effect of the channel on growth may be due to enhanced or new expression activity during development. Consistent with this mechanistic argument, expression analysis from resting and regenerating fins reveal that the *lof* mutation results in an upregulation of *kcnh2a* in fins (**Fig 4C**). We further assessed if misexpression of *kcnh2* was sufficient to phenocopy the *lof* mutant. Upregulation of human *kcnh2* during development of the wild-type zebrafish fin (*ubi:hKCNH2;ubi:GFP*) was sufficient to cause localized overgrowth of rays autonomous to the fin clone (**Fig 4D-F**). Activating clones contained mostly mesenchymal expression and regular normal segmental lengths similar to the *lof* phenotype (**Fig 4B**, **S3 Fig**). The segmentation pattern in *kcnh2a/lof* fin clones is distinct from comparable experiments with another altered potassium channel, *kcnk5b/alf*, that also is sufficient to induce localized overgrowth [18]. These findings are supported by recent preprint detailing comparable analyses of *lof* and the role of *kcnh2a* in mediating size of the fin, though without the genetic lesion underlying the *lof* phenotype [20].

#### Identification of leucine transporter lat4a as a novel regulator of fin size

We furthered our analysis through cloning of a second mutant that we identified in an ENU mutagenesis screen for genes affecting adult phenotypes of the zebrafish. *nr21* is a dominant mutant that was identified by having small fins as an adult compared to wild-type siblings (**Fig 5A**). In addition to overall size, the length of lepidotrichia segments in *nr21* are smaller than in wild-type fishes (**S3 Fig**). Though the fins are smaller in *nr21* individuals, the *nr21* mutation does not have a statistically significant effect on overall body size (**S5 Fig**). We used a heterogeneity-based mapping approach to identify the *nr21* mutation [21]. Importantly, as *nr21* is dominant and has only a subtle homozygous phenotype (**S5 Fig**), we mapped *nr21* through exome sequencing of several families of wild-type siblings, which represent the recessive, homozygous chromosome at the *nr21* locus (**S6 Fig**). We mapped *nr21* to chromosome 15, where, in sequencing reads from paired *nr21* carriers, we identified a non-synonymous mutation (T200K) in a conserved domain of the amino acid transporter *lat4a* (**Fig 5B-C**, **S6 Fig**). Mammalian Lat4 is an L-type amino acid transporter that shuttles isoleucine, leucine, methionine and phenylalanine into the cell in an ion-and pH-independent manner [22]. Lat4 has been previously linked to overall growth in mice [23], but its role in size regulation or patterning is not known. We produced frameshift mutations *in cis* to the *nr21/lat4a* allele though CRISPR/Cas9-mediated gene editing. These induced mutations of the *nr21* allele resulted in reversion to wild-type fin phenotype, indicating that the *nr21* mutation was gain-of-function and the phenotype was not due to haploinsufficiency (**Fig 5A,D**). Zebrafish with homozygous loss-of-function mutations in *lat4a* are phenotypically normal and viable (**S7 Fig**).

**Fig 5.**
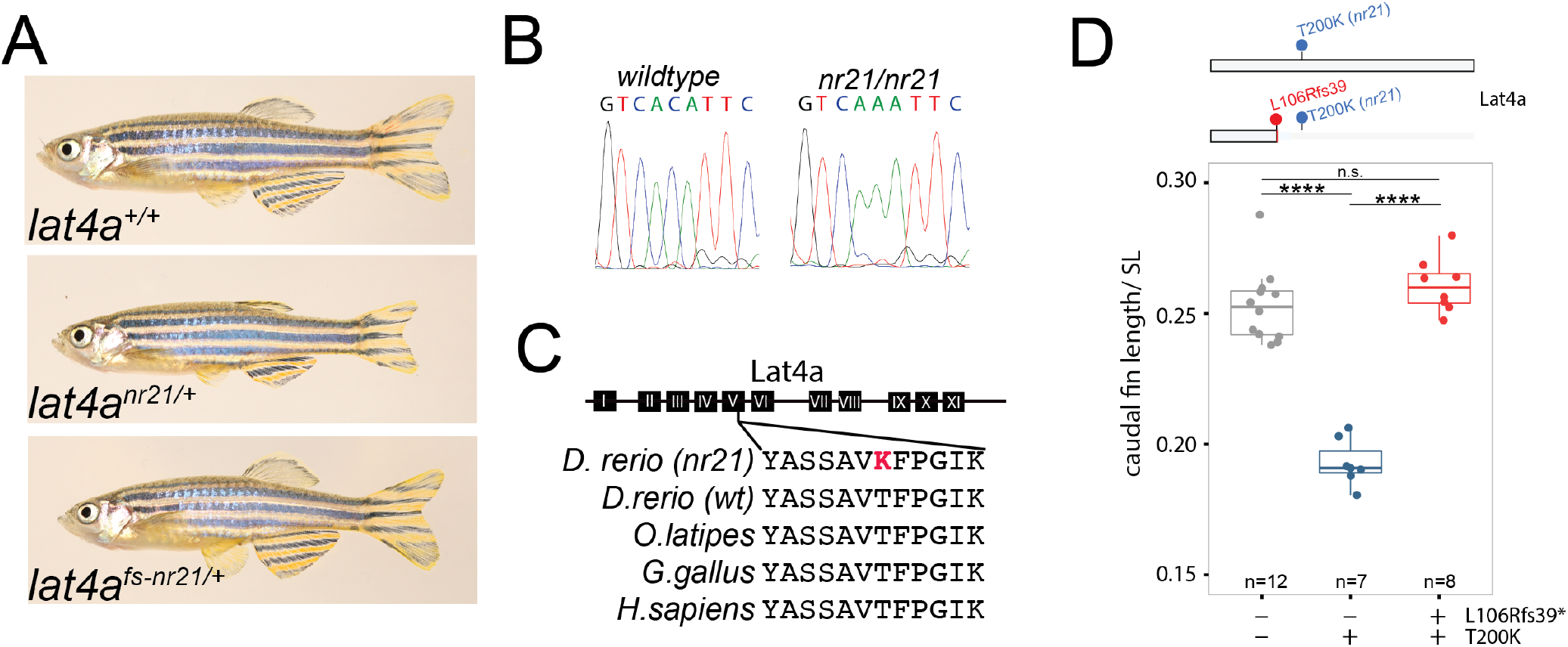
Identification of *lat4a* underlying the *nr21* mutant phenotype in zebrafish. **(A)** Images of wildtype, *nr21* short fin mutant and *nr21 in cis* frameshift revertant (L106Rfs39, *nr21*) zebrafish. **(B)** Sanger trace of causative *nr21* nonsynonymous substitution (C/A). For mapping details see **S6 Fig. (C)** Multiple sequence alignment of Lat4a showing the predicted T200K substitution of *nr21* within a highly conserved transmembrane domain. **(D)** Caudal fin length normalized to fish standard length (SL) in wild-type, *nr21* and revertant mutants. Schematic indicates location of mutations on Lat4a. **** indicates Tukey’s HSD adjusted p-value ≤ 0.0001. n.s. indicates not significant.

#### Convergent genomic signatures of fin allometry

With a broadened complement of genes known to affect fin proportion stemming from mutagenesis work defined here and previously, we compared and contrasted this work in model organisms with patterns of genomic variation in gliding beloniforms.

The independent origins of gliding behavior and elongated pectoral fin morphology in flying fishes and flying halfbeaks provide an opportunity to define genomic patterns that may be associated with gliding evolution. We observed an increase in convergent substitutions across the three lineages of gliding beloniforms compared to topologically similar control distributions, which contained between 14-28 such amino acid substitutions (**S8 Fig**). We identified 44 amino acid substitutions that are present only within all three lineages of gliding beloniforms (**Fig 6**, **S8 Fig**, **S10 Table**).

**Fig 6.**
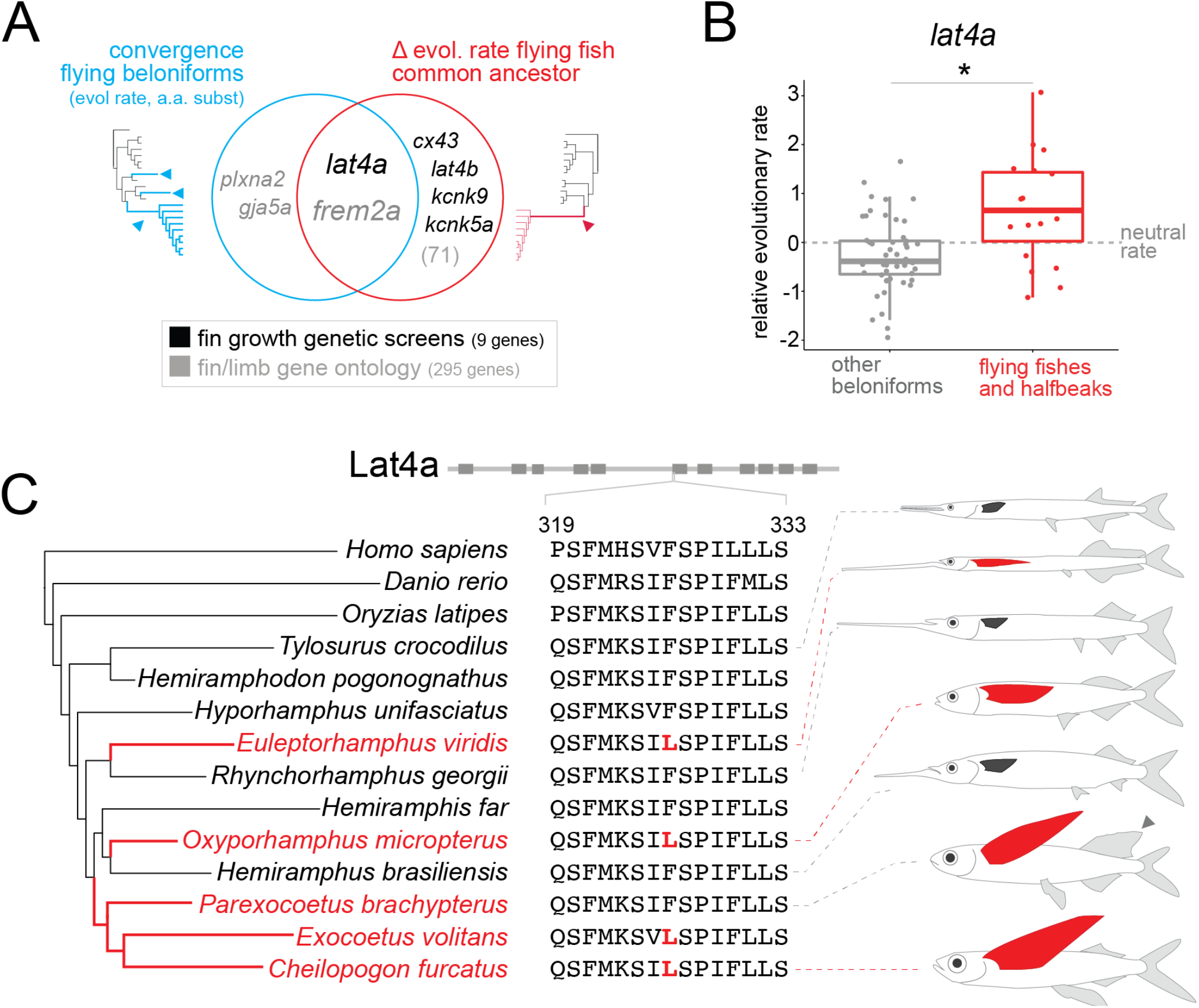
Convergent and ancestral genetic signatures in fin and limb genes in the evolution of gliding beloniforms. (**A**) Intersection of molecular convergence and changes to evolutionary rate in key fin-associated genes, as defined from genetic screens and gene ontology sets, at key nodes in the Beloniformes phylogeny associated with flight. See **S11 Table** for full gene list. We detected parallel amino acid substitutions (**S10 Table**) and convergent changes to relative evolutionary rate (**S12 Table**) in gliding beloniforms. We further identified sequence elements under accelerated or constrained evolution in the ancestral flying fish lineage (Exocoetidae; **S13 Table**). (**B**) Elevated relative evolutionary rate in *lat4a* in gliding beloniforms. Each dot represents a node in the beloniform phylogeny. * indicates p-value <0.05 for two-tailed test. Lat4a also has an phyloP acceleration p-value of 0.008 on the common ancestral node of flying fishes. (**C**) Convergent amino acid substitution in a highly conserved region of Lat4a. This substitution is found in both lineages of flying halfbeak and all flying fishes (Exocoetidae) with the exception of the sailfin flying fish, *P. brachypterus*, which has an elongated dorsal fin (gray arrowhead).

We intersected the identified convergent amino acid substitutions with known genes defined from forward genetic screens in zebrafish and from studies of fin and limb development (**Fig 6A**, **S11 Table**). Intriguingly, we detected a convergent amino acid substitution in *lat4a* that is found across multiple lineages having enlarged pectoral fins (**Fig 6A**, **S10 Table**). The *lat4a* gene tree suggests the convergent amino acid substitutions are not a result of incomplete lineage sorting in beloniforms, but likely evolved independently in all three lineages (**S9 Fig**). The *lat4a* locus not only harbors convergent amino acid substitutions, but has an accelerated rate of evolution in the gliding beloniforms (**Fig 6B**). Intriguingly, only one flying fish species in our dataset lacks the convergent *lat4a* allele, the sailfin flying fish *Parexocoetus brachypterus* (**Fig 6C**). This species is unique amongst the sequenced flying fishes in having an elongated dorsal fin. The evolutionary rate of *P. brachypterus lat4a* is highly accelerated and is distinct from that of other flying fishes (**Fig 6B**), revealing unique evolution of this gene in this species.

In addition to analysis of convergent amino acid substitutions, we further explored convergent shifts in evolutionary rate in flying fishes. We identify 48 genes (out of 17,937 analyzed) as having convergent changes to evolutionary rate after taking into account false discovery rate correction, though few of these genes are known to be involved in fin and limb development (**Fig 6**, **S11** and **S12 Table**). Of note, the potassium channel *kncj13* shows a weak signature of convergent evolutionary rate, though is not significant after multiple hypothesis correction. Overexpression of *kcnj13* in zebrafish, like *kcnk5b* and *kcnh2a*, is sufficient to induce fin overgrowth [24].

#### Ancestral genomic footprints of flying fish evolution

In addition to analyses of molecular convergence in gliding beloniforms, we explored patterns of accelerated or constrained evolution along the ancestral branch leading to the common ancestor of all flying fishes, where the shared elongated pectoral fin allometry in Exocoetidae is likely to have first emerged. While we observed no statistically significant enrichment for accelerated or constrained evolution in any specific gene ontology grouping across the coding regions along this ancestral branch, several genes involved in fin and limb development are under varied selection regimes. Intriguingly, among the accelerated genes in the flying fish common ancestor is *lat4a*, which also harbors convergent amino acid substitutions in flying halfbeaks (**Fig 6, S13 Table**). There is also accelerated sequence evolution of *kcnk9*, overexpression of which is sufficient to drive fin overgrowth in zebrafish [24], and in *kcnk5a*, the paralog of the long-fin zebrafish mutant *kcnk5b^alf^* [18] (**Fig 6, S13 Table**). Additionally, the transcription factor *evx1*, which regulates joint patterning in the fin lepidotrichia [25], is also under accelerated sequence evolution in ancestral flying fishes (**S13 Table**), and shows weak convergence in relative evolutionary rate (**S12 Table**). In contrast, the gap junction protein *cx43*, the absence of which causes short fins in zebrafish [26,27], is under constrained sequence evolution along the ancestral branch of flying fishes (**Fig 6**, **S13 Table**). Retinoic acid signaling is involved in outgrowth and patterning of limbs and is active in a proximal-distal gradient [28]. Interestingly, the retinoic acid-inactivating enzymes Cyp26b1 and Cyp27c1 are both under accelerated sequence evolution in our dataset (**S13 Table**). Further supporting a potential role for retinoic acid signaling in beloniform fin growth, we also observed constraint in the evolution of the retinoic acid synthesis gene *aldh1a2* in gliding beloniforms (**S12 Table**), though this signature is not significant after false discovery rate correction.

#### Regulatory changes near fin and limb developmental genes

Using a nearest-neighbor algorithm, GREAT [29], we assigned CNEs to neighboring genes to assess whether there is selection in putative gene-regulatory regions in the common ancestor of flying fishes. Similar to analysis of coding regions (**Fig 3**), we see a significant enrichment for accelerated sequence evolution in CNEs near multiple gene classes associated with gliding behavior and morphology (**Fig 7**, **S14-17 Table**), specifically those genes associated with pectoral fin and appendage development (**Fig 7**, **S14 Table**). A particularly intriguing locus is *sall1a. Sall1a* is required for pectoral fin development in zebrafish [30], limb development in mouse [31], and has been implicated in the reduced forelimbs of emus [32]. There is widespread accelerated sequence evolution at the *sall1a* locus along the ancestral flying fish branch, both in coding and non-coding regions (**S10 Fig**, **S13-14 Tables**). Intriguingly, the *sall1a* locus also exhibits a convergent shift in evolutionary rate in the gliding beloniforms, though is not significant after false discovery rate correction (**S11 Table**).

**Fig 7.**
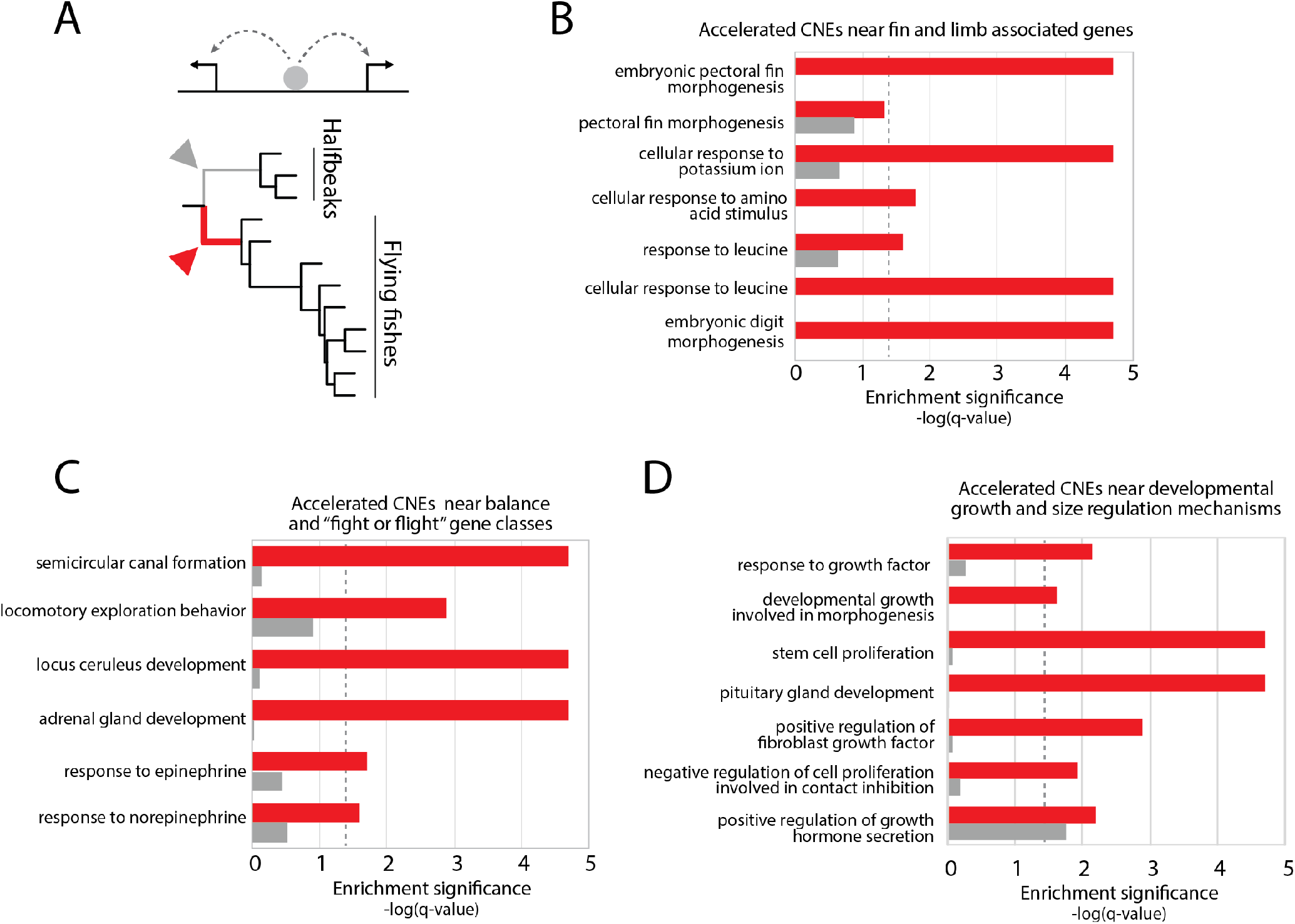
Accelerated sequence evolution in predicted gene regulatory regions that are associated with morphological and behavioral traits in flying fishes. **(A)** Analysis of accelerated sequence evolution using the program phyloP, with a focus on the ancestral common ancestor of flying fishes (Exocoetidae; red) compared to the sister lineage of halfbeak fishes (gray). CNEs were assigned to neighboring genes through the GREAT approach [29] for gene ontology enrichment analysis. (**B-D**) Significant enrichment of accelerated sequence evolution in CNEs near genes associated with fin and limb development (**B**), balance and fear response (**C**), and general organ and tissue growth and size regulation (**D**). For full list of significantly enriched terms under acceleration and constraint, see **S14-17 Tables**.

Interestingly, our CNE data also showed congruence with mutational analyses of the zebrafish. Specifically, in the common ancestor of flying fishes we see enrichment for accelerated sequence evolution in CNEs near genes involved in the cellular response to potassium ion, and near genes involved in cellular response to amino acids, specifically leucine, one of the amino acids transported by Lat4 (**Fig 7**, **S14 Table**) [22].

### Sufficiency of simple genetic changes to phenocopy flying fish morphology in zebrafish

Given the observed comparative genomic signatures in Beloniformes in genes and in CNEs that are involved in potassium regulation and amino acid transport, we asked how these genetic mechanisms may interact to regulate fin size and patterning.

Double transheterozygous mutants between *kcnh2a^lof/+^* and *lat4a^nr21/+^* are viable and show specific variation in proportion among fins. While all fins of *kcnh2a^lof/+^* are overgrown and all fins of *lat4a^nr21/+^* are shortened, the *lat4a^nr21/+^; kcnh2a^lof/+^* transheterozygous fish showed wild-type sized medial fins while the paired fins were overgrown (**Fig 8**). Thus, *lat4a^nr21/+^* is a medial fin suppressor of *kcnh2a^lof/+^*, exposing fin-type specific regulation of size. Surprisingly, the caudal fin of *kcnh2a^lof/+^; lat4a^nr21/+^*, while generally wild-type in size, is hypocercal, with an elongated ventral lobe. The overall fin morphology pattern, and general bauplan, of the transheterozygous fish is thus remarkably similar to the fin phenotype of flying fishes (**Fig 8**). The interaction between these two genes, and the changes in developmental regulation in the zebrafish is sufficient to phenocopy this phylogenetic key innovation.

**Figure 8.**
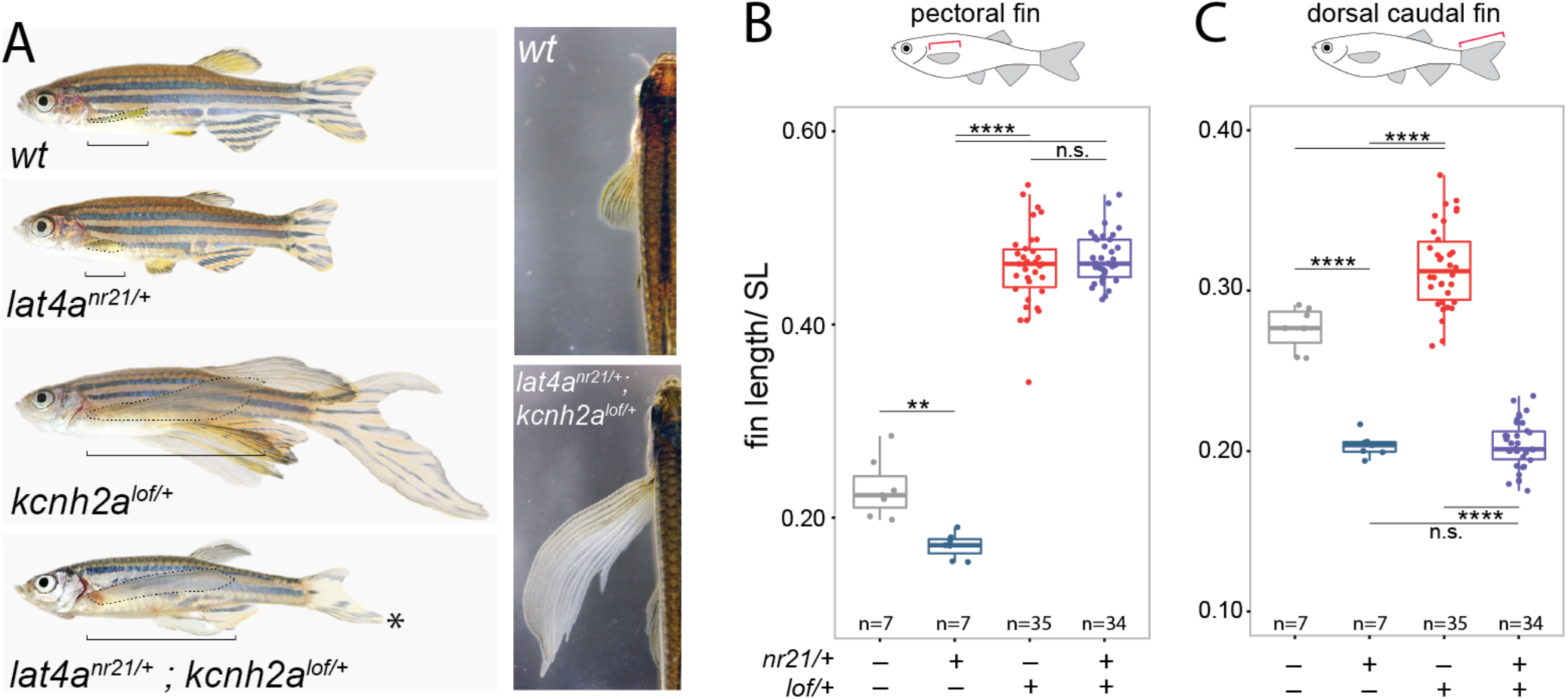
Intersection of bioelectric and Lat4a-mediated growth pathways is sufficient to phenocopy the evolved flying fish fin allometry. (**A**) Images from representative fish from a *lat4a^nr21/+^* x *kcnh2a^lof/+^* cross indicate *lat4a* is a fin suppressor of overgrowth caused by increased potassium channel function. Brackets indicate length of pectoral fin. Pectoral fins outlined with dashed line. As in flying fishes, the *nr21/lof* transheterozygotes exhibit elongated paired fins and a hypocercal caudal fin. Asterisk (*) highlights hypocercal caudal fin (**B-C**) Pectoral fin length (**B**) and caudal fin length (**C**) suppression of overgrowth in medial fin by lat4a. In (**B-C**), length normalized by standard length (SL); p-values generated through Tukey’s HSD. **** adjusted p-value ≤ 0.0001, ** adjusted p-value ≤0.01. n.s. indicates not significant.

## Discussion

### Discovery of novel regulators of growth and form

Here, we map the mutation in the zebrafish *lof* line and further identify an inversion on chromosome 2 responsible for this mutant. The inversion results in ectopic overexpression of the voltage-gated potassium channel *kcnh2a* in the fin (**Fig 4**) which we show is sufficient to drive coordinated overgrowth of the fin. This finding adds to a growing body of evidence that potassium channels and bioelectric signaling are instrumental in regulating fin size [16,24,33]. It is unknown whether there is a specific role for Kcnh2a protein in fin growth, or if *kcnh2a* is just one of many potassium channels where overexpression is sufficient to drive fin overgrowth [18,24]. Importantly, there are subtle phenotypic differences in patterning and growth between *kcnh2a^lof^* and other potassium-related overgrowth mutants, namely *kcc4a^schleier^* and *kcnk5a^alf^*, which have irregular segmentation and blood vessel patterning [18,34]. In contrast, *kcnh2a^lof^* fins appear to have normal segmentation and vessel patterning. Thus, there is at least some distinction between the growth mechanisms across different classes of potassium channel mutants.

We further identify Lat4a as a novel regulator of growth and proportion during development. The function of the amino acid transporter *lat4a* in development is not well defined. We show that dominant mutations in *lat4a* suppress relative fin size in the zebrafish. Zebrafish with *lat4a* loss-of-function mutations are viable (**S7 Fig**); this could potentially be due to compensation by the *lat4b* paralog as has been shown for other developmental genes with close paralogues. Transporters such as *lat4a* regulate amino acid levels within the cell, and thus can have complex and pleiotropic effects on cell nutrient sensing and growth pathways [35]. Reflecting these potential roles in regulating growth, Lat4 knockout mice are smaller than littermates [36], and many of the L-type amino acid transporter (LAT) family members are upregulated in human cancer, including *LAT4* [37,38]. However, the mechanism by which these amino acid transporters regulate growth is not well understood, particularly in the broader context of bioelectric signaling. Lat family members transport methionine, which can stimulate cell proliferation [39]. Additionally, Lat4 family members are a primary means of leucine uptake, which is an activator of TOR signaling [38]. The link of amino acid transport to cellular growth mechanisms and TOR signaling provides an intriguing mechanistic link to bioelectric signaling cascades.

### Bioelectric signaling and fin size regulation

With the notable exception of the zebrafish *rapunzel* mutant [40] caused by a mutation in a teleost-specific gene of unknown function [41], all zebrafish fin overgrowth mutants currently mapped are caused by an alteration in potassium channel function [16]. This pattern across a class of mutants that cause coordinated fin overgrowth indicates an important role for bioelectric signaling in regulation of size and proportion. However, the action of potassium channels in the regulation of growth and proportion remains an outstanding question.

Fin overgrowth can be caused through both mutation as well as overexpression of potassium channels from multiple channel classes, including two-pore domain (K2P) [18], voltage gated (Kv) (**Fig 4**), inwardly-rectifying (Kir) channels [24], and potassium-chloride co-transporters [34]. In the case of misexpression of the potassium channels *kcnk5b* and *kcnj13*, conductance is necessary for the overexpression phenotype [18,24]. Fin clone analysis here (**Fig 4**) and in Perathoner *et al*. [18] both demonstrate a role for potassium channel-expressing fibroblasts in regulating fin growth. Additionally, at least one zebrafish potassium channel mutant, *kcc4a/schleier* [34], as well as the chemical calcineurin inhibitor FK506 [42] both require the presence of *kcnk5b* to stimulate fin overgrowth, indicating that there may be some common molecular regulation of bioelectricity in fin growth and/or a specific molecular connection of bioelectricity to a core cellular and developmental growth pathway.

### Convergence, developmental constraint and evolution of a flying fish bauplan

While it is difficult to disentangle mutations of adaptive value from those resulting from neutral drift and bottleneck along a single lineage, parallel substitutions occur with low probability in models of neutral evolution [43]. This low probability makes parallel genetic changes strong candidates for association with specific phenotypes when they occur in lineages with independently-derived character changes [44]. Importantly, while the number of convergent substitutions across species with convergently evolved traits is frequently similar to control comparisons lacking known trait convergence [45,46], a recent analysis of multiple instances of molecular convergences in mammals indicates that convergent amino acid substitutions are not randomly distributed in the genome and are likely associated with genes related to key phenotypic traits [47]. Regardless of frequency, parallel amino acid changes are thus compelling candidates for functional association with convergent evolutionary traits. Here we show specific convergent changes have occurred in flying fishes and flying halfbeaks. One of these genes, *lat4a*, we identify as important in regulating fin size in the zebrafish, a distantly related species of cypriniform fish having a shared common ancestor with Beloniformes >200 My ago [6]. We demonstrate sufficiency of *lat4a* to modify bioelectric growth cues in the zebrafish, leading to a flying fish-like fin pattern with specific elongation of paired fins and shortened medial fins (**Fig 8**).

Aerial gliding in fishes has evolved several times independently in teleosts [2], and multiple examples of the ‘flying fish’ morphotype can be seen in the fossil record [48,49]. That such a bauplan has evolved multiple times may represent a developmental bias in shaping morphology available for selection. There appear to be numerous, adult-viable paths through which bioelectric signaling can impact fin size, ranging from point mutations to regulation of gene expression. That simple genetic changes can lead to coordinated alteration in fin patterning may underlie the general evolvability and diversity of fin allometry in fishes. Future studies of other lineages of ‘flying fishes’, such as gasteropelicid flying hatchet fishes or osteoglossid butterfly fishes, as well as experimental investigations the other convergent genetic signatures we detect in gliding beloniforms (e.g. **S10** and **S12 Table**), may provide further insight into the morphology and physiology of this remarkable behavior.

### Targeted sequencing and comparative genomics

In parallel to advances in whole genome sequencing, targeted sequencing studies have effectively and efficiently probed both patterns of genome evolution and have identified developmental regulation underlying trait evolution [9,10,50]. The majority of comparative genomic studies asking questions over large evolutionary distances of many millions of years tend to focus on highly conserved protein coding sequences and non-coding elements due to the lack of clear homology and/or obvious functional roles for large swaths of the genome. Targeted sequencing results in high sequencing read depth over functionally annotated and conserved regions, enabling efficient population analysis on top of comparative genomics and candidate gene analysis. By sequencing population samples, a degree of population variation can be assessed and the pitfalls of identifying of species-defining SNPs based on a single individual can be avoided. Genetic analysis of large numbers of species is also made more economic by targeted sequencing, and as in whole genome sequencing, analysis of large numbers of species can clarify and refine comparative genomic signatures [51].

In addition to gliding behaviors mentioned here, Beloniformes exhibit numerous independently evolved traits that are useful in answering a variety of fundamental questions in biology, from physiological ecology (*e.g*. marine to freshwater transitions), to development (*e.g*. craniofacial morphology), and reproduction (*e.g*. egg buoyancy) [3,5,52]. Further, the complex jaw morphologies of halfbeaks and needlefish represent a classic example in the study of ontogeny and developmental timing (heterochrony) in evolution [5,11,53,54]. The phylogenetic sequencing data for Beloniformes will be a powerful resource for unraveling the genetic basis of these and other morphological and behavioral traits within this clade.

### Summary

The detection of comparative genomic signatures associated with bioelectric signaling and amino acid transport in flying fishes paired with functional association points to a role for these developmental regulators in the development and evolution of allometry. This combination of forward genetic approaches, both experimental and evolutionary, is a powerful means to parse out regulatory pathways of development and physiology.

## Materials and methods

### Zebrafish husbandry and lines

Zebrafish were housed and maintained as described in [55] and performed in accordance with Institutional Animal Care and Use Committee (IACUC) guidelines at Boston Children’s Hospital and Washington University in Saint Louis. A description of the husbandry and environmental conditions in housing for the fish used in these experiments at Boston Children’s Hospital is available as a collection in protocols.io (https://doi.org/10.17504/protocols.io.mrjc54n). Similar conditions were present at Washington University Medical School (SLJ lab). For all experiments, adult stages were defined by reproductively mature fish ≥ months old. Males and females were used together in analyses as both sexes showed comparable overgrowth and growth regulation. Longfin alleles used in this work are *lof^dt2^*, the revertants *lof* (*lfr1*(Y669N), *lfr2*(L739Q), and *WL7* (Y418X). These lines are no longer maintained nor frozen, thus are used for confirmation of mapping only. The deletion line *lof^j6e1^* [19] (lof^Df(Chr02:csnk1g2a,rnf2,kifap3b)j6g1/j6g1^) is present at ZIRC repository (ZL1494). Lat4a alleles identified and used in this work are *lat4^nr21^* (*dmh26*), and *lat4a^del^ (mh152*).

### Chemical and gamma-ray mutagenesis

For identification of *lof^dt2^* revertants, homozygous *lof^dt2^* mutant males were mutagenized with 3.3μM N-ethyl-N-nitrosourea (ENU) for four repeated doses following Rohner *et al*. [56]. Founder mutagenized males were crossed to Tübingen wild-type females. F1 progeny were scored for a reduction or increase in the fin length compared to heterozygous *lof* fish. Radiation induced mutations were generated by exposing sperm from *lof^dt2^* homozygous males to 250 rads of gamma-irradiation [19,57]. Irradiated sperm was used to fertilize eggs from C32 females. Progeny exhibiting wild-type phenotypes were recovered as potential *lof^dt2^* revertants. Identification and description of *lof^j6e1^* is described in Iovine and Johnson [19].

### Mapping and mutational analysis of *lof* (*dt2*)

The *lof^dt2^* mutant was independently mapped in both the MPH and SLJ laboratories using bulk segregant analysis and analysis of both simple sequence length polymorphisms (SSLP) and single nucleotide polymorphisms (SNPs) [19]. As both datasets were consistent, the mapping data were combined to refine the identified mapping interval on chromosome 2 (**S4 Fig**). As a complementary approach, both MPH and SLJ laboratories performed revertant screens, as described above, and identified independent nonsynonymous mutations in *kcnh2a in cis* to the *lof* allele (*lfr1, lfr2* and *WL7*) that were essential for expression of the long-finned phenotype. In a similar approach, γ-ray irradiation of *lof^dt2^* founders were used to generate deletions within the *lof* chromosome to map region of mutation.

In the absence of coding variants within *kcnh2a*, we sought to identify the causative genetic lesions underlying the *lof* phenotype. Given the suppression of recombination over a large mapping interval (~1 Mb), and to account for the possibility of structural variants in *lof*, we reconstructed the chromosome 2 locus using long sequencing reads from a *lof^dt2^* incross (NCBI Sequence Read Archive ERX1428166). As the revertant screens identified *kcnh2a* as necessary for the *lof* phenotype, we searched for single molecule PacBio sequencing reads from a *lof^dt2^* incross that contained a portion of *kcnh2a* using BLASTN (ncbi-blast-2.2.30+; parameters ‘-max_target_seqs 1 - outfmt 6’). Candidate reads were then assembled into contigs using the Canu assembler [58] (parameters ‘genomeSize=0.01m - pacbio --minInputCoverage=2 stopOnLowCoverage=2’). Both individual reads, and the assembled contig identified a breakpoint upstream of the transcription start site of *prrx1a* (~23.174 Mb in Zv11) and one between *kcc4a* and *kcnh2a* (~24.075 Mb) (**S4 Fig**). We further refined these contigs by identifying reads that contained a high-quality match (BLASTN) to each side of the identified breakpoint. This merged assembly resulted in two contigs, one for each breakpoint (**S11 Fig**). Notably, we did not generate contigs representing the wildtype (Zv11) chromosome organization at these breakpoints. The breakpoint between *kcc4a* and *kcnh2a* is within a DNA transposon that is not present between these genes in the zebrafish reference genome (Zv11) and is likely part of a transposable element expansion in this region in *lof* relative to the Tübingen strain (**S4 Fig C**, **S11 Fig**).

### Overexpression constructs

pmTol2-ubi:hKCNH2, ubi:eGFP was constructed by inserting a ubi:hKCNH2 and a ubi:eGFP cassette into the multiple cloning site of pminiTol2 (Addgene #31829).

The ubi:hKCNH2 and ubi:eGFP cassettes were obtained by inserting either the full length coding sequence of hKCNH2 with the SV40 late polyadenylation signal (SV40pA) or eGFP from pME-eGFP (Tol2kit #383) downstream of the ubi promoter of pENTR5’_ubi (Addgene #27320). Plasmids (20 ng/μl) and Tol2 transposase mRNA (25 ng/μl) were injected into single cell zebrafish embryos. Adults were screened for GFP+ cell clones containing the integrated plasmid and the impact of GFP+ fin clones on fin growth was assessed.

### Isolation and mapping of *nr21* mutant line

*nr21* was isolated in a small-scale dominant screen. In this screen, wild-type male founders were mutagenized with ENU as above and first-generation outcross progeny screened for fin phenotypes. A single *nr21* founder was isolated as having a dominant, shortened length of all fins compared to wild-type. This mutant was outcrossed to establish the mutant line. As *nr21* is dominant and without an obvious homozygous phenotype, we could not easily isolate homozygous *nr21* individuals for mutation mapping by heterozygosity. Instead, we used homozygosity by descent to identify the recessive, wild-type chromosome [59]. However, given normal variability in zebrafish, more than one wild-type chromosome can be present in the F2 generation (e.g. AB^1^, AB^2^). To reduce the likelihood of mixing parental wild-type chromosomes (AB^1^,AB^2^), in which polymorphisms show up as heterozygosity, three wild-type F2 families were mapped (**S6 Fig**). Next generation sequencing libraries were made from pools of 10 wild-type siblings from each of three separate F2 crosses. Additionally, in order to identify candidate mutations, we also sequenced a pool of 10 short-finned *nr21* individuals from a mixture of all three F2 families. DNA was extracted from fin clips using Qiagen DNeasy Blood & Tissue Kit. Barcoded sequencing libraries were prepared as in Bowen *et al*. [60], and hybridized to a custom Agilent SureSelect 1M Capture Array (cat# G3358A) targeting the zebrafish exome (Zv9). The zebrafish design encompassed 974,016 probes of 60 bp length that are tiled on average every 40bp (20bp overlap) over the 41 Mb exome. Sequencing libraries were sequenced as 100bp single end reads on an Illumina HiSeq 4000. Sequencing reads were aligned to the zebrafish genome with Novoalign (http://novocraft.com/), and patterns of heterogeneity used to map the *nr21* chromosome as in Bowen *et al*. [59].

### CRISPR-Cas9 gRNA design and injection

CRISPR guide RNAs (gRNAs) were designed against the third coding exon of *lat4a* using the online ChopChop tool to limit predicted off-target gRNA cutting [61,62]. Given the variable success rates of individual gRNAs, a “blanket” of 5 gRNAs targeting the same exon were synthesized and injected as a pool. gRNAs were assembled according to Gangon *et al*. [63]. Briefly, oligos containing the gRNA target were annealing to a universal oligo containing the tracrRNA and SP6 promoter. The annealed oligo ends were then filled in with T4 polymerase for 20 minutes at 12°C. The gRNA was synthesized from this oligo template using Ambion MEGAscript SP6 Kit. For transcription efficiency, the first two bases of each gRNA were changed to ‘GG’ as there is evidence that these bases have less effect on Cas9 cutting efficiency or off-target binding than mutations closer to the PAM site [64–66]. gRNAs were injected into single cell zebrafish embryos at a concentration of 50 ng/μl gRNA pool and 600 ng/μl Cas9. To screen for deletion efficiency, the target exon was amplified from pools of three 24-hour embryos and the resulting amplicons were heated to 95°C and cooled at - 0.1°C per second to form heteroduplexes. Following heteroduplex PCR, a T7 endonuclease digestion for 30 minutes at 37°C in NEB Buffer 2 was used to generate deletions in the presence of Cas9-induced indels. PCR primers for T7 endonuclease assay: 5’-CACTGAAAACTGACCACAAGTCA-3’, 5’CAGCACTTCCCAGAGTGTCA-3’.

Full gRNA oligo (target sequence underlined, 5’-3’):

1. ATTTAGGTGACACTATA<u>GGCCCTGTACCGTTACCTGG</u>GTTTTAGAGCTAGAAATAGCAAG
2. ATTTAGGTGACACTATA<u>GGTGAATGCCACAAGACTTG</u>GTTTTAGAGCTAGAAATAGCAAG
3. ATTTAGGTGACACTATA<u>GGATGCCACAAGACTTGAGG</u>GTTTTAGAGCTAGAAATAGCAAG
4. ATTTAGGTGACACTATA<u>GGCCACAAGACTTGAGGAGG</u>GTTTTAGAGCTAGAAATAGCAAG
5. ATTTAGGTGACACTATA<u>GGAGTGTGATGGCGCTCAGG</u>GTTTTAGAGCTAGAAATAGCAAG

Universal constant oligo:

5’AAAAGCACCGACTCGGTGCCACTTTTTCAAGTTGATAACGGACTAGCCTTATTTTAACTTGCTATTTCTAGCTC TAAAAC-3’

### qRT-PCR

Uncut and regenerating caudal fins were homogenized in TRIzol Reagent (Invitrogen) for RNA extraction. cDNA synthesis was performed SuperScript III Reverse Transcription Kit (Invitrogen) and oligo dT primers. We performed qRT-PCR using the Power SYBR Green Master Mix (Applied Biosystems) on the Applied Biosystems ViiA 7 Real Time PCR System. Cycling conditions: 10 minutes at 95 °C; 40 cycles of 15 seconds at 95 °C followed by 1 minute at 58 °C; melting curve analysis with 15 seconds at 95 °C, 1 minute at 58 °C and 15 seconds at 95 °C. Temperature was varied at 1.6 °C/s. Expression levels of *kcnh2* were normalized relative to *gapdh*. Fold expression of *kcnh2a* in wild-type and *lof* was calculated using 2^-(△△Ct)^. *kcnh2a* primers: 5’-GCTGGCAGATCAGCAGGACA-3’, 5’-CGTTCAGGTAGGGAGAGCAG-3’*. gapdh* primers: 5’-GGACACAACCAAATCAGGCATA - 3’, 5’-CGCCTTCTGCCTTAACCTCA-3’.

### Targeted Sequence Capture Design

We based the sequence capture design on primary on the medaka reference genome (*Oryzias latipes*, MEDAKA1), which was the only genome from Beloniformes available when this study was initiated. However, to account for the possibility that specific genetic regions may be absent or in drift in the medaka genome but conserved in the suborder Belonoidei, we included regions from the outgroup genomes of *Poecilla formosa* (Poecilia_formosa-5.1.2) and *Xiphophorus maculatus* (Xipmac4.4.2) (**Fig 2**). To identify annotations in *P. formosa* and *X. maculatus* that were absent or not well-conserved in *O. latipes*, we used BLASTN (ncbi-blast-2.2.30+; parameters ‘-max_target_seqs 1 - outfmt 6’). Any element from *P. formosa* or *X. maculatus* with an E-value >0.001 and/or covered <70% in the *O. latipes* genome were included in the capture. If the element missing in medaka was found in both *P. formosa* and *X. maculatus*, we included the *X. maculatus* version. To avoid adding DNA regions that are specific to Poeciliidae, DNA sequences from *P. formosa* and *X. maculatus* were identified by BLASTN (E-value <0.001) in the genome of an additional outgroup, the Nile tilapia (*Oreochromis niloticus;* Cichlidae; Orenil1.0), prior to inclusion in the capture. SeqCap EZ Developer (cat #06471684001) capture oligos were designed in collaboration with the Nimblegen design team to reduce probe redundancy, standardize oligo annealing temperature and remove repetitive regions.

We generated two sequence capture designs for targeted enrichment of beloniform sequencing libraries, one for protein-coding exons and another for conserved-non-coding elements (CNEs, miRNA, UCNEs). For each genome, protein coding exons were extracted from Ensembl BioMart [67]. CNEs were defined from the constrained regions in the Ensembl compara 11-way teleost alignment [68]. We removed CNEs < 75bp in length to facilitate space in the capture design. In addition to constrained non-coding elements, miRNA hairpins were extracted from miRbase and ultraconservative elements (UCNEs) from UCNEbase and included in the capture design [69,70]. We padded miRNA hairpins to be > 100bp, and removed CNEs, miRNAs and UCNEs that overlapped coding exons using Bedtools (v2.26.0) intersectBed [71]. We prioritized miRNA and UCNE annotations where these overlapped with CNE annotations.

The final capture design for protein coding exons encompassed a total of 225,182 regions and targeting 38,295,260 total bases. The CNE design encompassed 119,080 regions and targeted 17,983,636 total bases. Importantly, we recovered an average of 61.4% read coverage of targeted regions in the genome of *X. maculatus* and 41.5% coverage in *P. formosa*. Recovery of these targeted regions from outgroup genomes that lack clear homology to the *O. latipes* reference genome indicates conservation in Belonoidei with specific loss in *O. latipes*. That we recovered reads from these genomes highlights the importance of a taxonomically diverse capture probe set in minimizing genome bias in cross-species sequence enrichment (**S4 Table**).

### Specimen tissue collection and sequencing library preparation

Tissues were acquired from archived samples in the NRL laboratory and the Royal Ontario Museum (**S3 Table**). DNA was extracted from multiple individuals of each species using the Qiagen DNeasy Blood & Tissue Kit (**S2, S3 Table**). Equal quantities of DNA from each individual were pooled for each species prior to library preparation and sequencing. The DNA pools were diluted in a shearing buffer (10 mM Tris, 0.1 mM EDTA, pH 8.3) and were mechanically sheared to an average size of 200bp in a Covaris E220 ultrasonicator (duty cycle, 10%; intensity, 5; cycles/burst, 200; time, 400 seconds; temperature, 8°C). Barcoded sequencing libraries were generated using a KAPA HyperPrep Kit (Roche, No. 07137923001) following the Nimblegen SeqCap EZ Library protocol (Version 4.3) and using dual-SPRI (solid phase reversible immobilization) size selection to generate libraries of 200-450 bp. The libraries were hybridized to the capture baits according to the SeqCap EZ Library protocol. In place of Human CotI DNA, the SeqCap EZ Developer Reagent was used (cat #06684335001) during hybridization. Additionally, libraries were hybridized to the capture baits and washed at a reduced stringency of 45°C relative to the manufacturer recommended temperature of 47°C in order to allow extra mismatches for crossspecies hybridization. Multiple barcoded libraries were then pooled for 100 bp single-end sequencing via Illumina HiSeq 2500.

### Reference contig assembly

To generate reference contigs for each species, we followed the Phylomapping *de novo* contig assembly pipeline as described in Daane *et al*. [10]. Briefly, low-quality bases in sequencing reads were masked and sequencing reads de-duplicated with the FASTX toolkit (http://hannonlab.cshl.edu/fastx_toolkit). Sequencing adaptor sequences were removed using Trimmomatic v.0.36 [72]. Sequencing reads were binned by orthology to target regions in the *O. latipes*, *X. maculatus* and *P. formosa* reference genomes using BLASTN and dc-megaBLAST. Binned reads were then assembled into contigs *de novo* using CAP3 and UCLUST [73,74]. Sequencing reads were aligned back to the assembled contigs using NextGenMap to recruit previously unmapped reads to the binned assembly [75]. All reads were re-assembled using CAP3 in a second round of *de novo* assembly. If multiple contigs are assembled for any given element (for example, CNE, exon), the multiple contigs were then merged if they overlapped and had >95% identity. This results in consensus contig sequence for each target region. See Daane *et al*. [10] for details.

### Identification of orthologs

As in Daane *et al*. [10], we paired orthologous sequences between species using a gene tree-species tree reconciliation approach. Contigs were automatically annotated according to the orthologous element within the reference genome that was identified using BLAST and subsequently used to scaffold and refine contig assembly. In the event that multiple contigs are assembled for a given targeted element, the multiple contigs for all species were aligned using MAFFT v7.313 (parameters ‘-op 10 - ep 10’)[76], and a maximum likelihood tree generated with IQTree v1.7beta2 (parameters ‘-alrt 1000’) [77]. To infer patterns of duplication and loss, gene tree reconciliation was performed using Notung-2.9 (parameters ‘-reconcile-rearrange-silent-threshold 90%-treeoutput nhx’). The total number of duplication and loss events across the phylogeny as estimated by Notung was then compared to a scenario where all copies are local duplicates. The most parsimonious scenario where the fewest gain/loss events occurred was then selected. Simulations of this ortholog pairing approach found accurate segregation of orthologs provided there ≥4-6% variation between paralogous sequences, which coincides with thresholds of variation necessary to distinguish copy number variation during contig assembly [10].

### Read coverage and depth of targeted regions

The targeted sequence captures were designed from existing reference genomes, which were then used to scaffold contig assembly. In order to assess capture efficiency and estimate the coverage of targeted regions, we need to identify where the coordinates on each assembled contig correspond to on the original reference genomes. We performed a pairwise alignment between each assembled contig and the targeted region of the genome using Biopython v.1.70 (parameters ‘pairwise2; match = 5, mismatch = - 4, gap_open = - 15, gap_extend = - 1’). Raw sequencing reads were re-aligned to the assembled contigs using NextGenMap v0.5.5 [75], and the data from the pairwise alignment, including indels relative to the reference genome, were used to lift the read alignment data from the contig to the reference genome. Alignments were converted from SAM to BAM format and indexed using SAMtools (v1.9) [78] and were visually inspected for accuracy in the Integrative Genome Viewer [79]. We defined coverage as the number of targeted bases overlapping at least one sequencing read. Coverage was calculated with BEDTools v2.23.0 [71], with read depth per base calculated with using the depth flag of coverageBed (parameters ‘-d’). Coverage was estimated at multiple levels of read depth (**S5 Table**).

The majority of the targeted regions had either 0 or 100% coverage (**S1 Fig**), with a wide distribution of depths in across species (**S1 Fig**). Among the genes with low coverage, we detected enrichment for gene classes associated with the cell adhesion, extracellular matrix proteins and the immune system (<25% coverage in ≥75% of exons; **S6 Table**). Enrichment was calculated with a Fisher’s exact test (SciPy v.0.18.1; fisher_exact), and multiple hypothesis test corrections using Benjamini-Hochberg FDR (python module statsmodels v.0.6.1; fdrcorrection0). The missing genes are comparable to similar cross-species targeted sequence enrichment experiments for other clades [9,10], indicating that these fast-evolving gene classes are less likely to be highly represented in these datasets.

### Nucleotide diversity and recovery of population variation

Where possible, we performed pooled sequencing of multiple individuals for each species (**S2,S3 Table**). We calculated genome-wide nucleotide diversity (π) across all sequenced regions using the program PoPoolation v1.2.2 (parameters ‘--min-count 2 --min-coverage 4’) [80]. We also distinguished population variation from polymorphisms that are fixed within the species (**S7 Table**). Mutations in the dataset for each species were considered heterozygous with a minimum allele depth of 2 sequencing reads and an allele frequency between 20-80% in the sequencing reads.

### Multiple sequence alignment

We used the program MAFFT v 7.313 to align orthologous sequences across our dataset (parameters ‘–maxiterate 1000-localpair-op 10-ep’) [76]. In the event the MAFFT alignment had an indel that disrupted a codon, we re-aligned these exons as codon alignments using the frameshift-aware multiple sequence alignment program MACSE v2.03 (parameters ‘prog alignSequences - seq - seq_lr - fs_lr 10 - stop_lr 15’) [81]. We masked the multiple sequence alignment using the program Spruceup (v 2020.2.19, parameters ‘data_type:nt, distance_method:uncorrected, window_size:6, overlap:5, fraction:1, criterion:lognorm, cutoffs:0.97’) [82]. Spruceup uses a lognormal distribution of genetic distances across the multiple sequence alignment to detect local outliers in the alignment. For more accurate estimation of genetic distances, we ran Spruceup on a concatenated set of all contigs in our multiple sequence alignment instead of on the much smaller individual exons or CNEs that make up the contigs.

### Beloniformes phylogeny

We used IQTree v1.7beta2 (parameters ‘-bb 1000 - st CODON - m MFP’) to calculate a maximum likelihood tree for each gene ≥400 bp in our dataset in which there was ≥70% coverage across all species (4,683 genes) [77]. For each gene we used ModelFinder to find the optimum codon model [83], and assessed support for phylogenetic relationships using 1,000 ultrafast bootstrap replicates [84]. We then estimated a species tree from the distribution of gene trees using ASTRAL v5.6.1 [85], with local posterior probabilities to support each quadpartition in the tree [86].

### Reconstruction of gene sequences

We reconstructed full gene sequences from our individual exon contig data. As in Daane *et al*.[10], we concatenated single copy exons in gene order as found in the reference genomes that were used to guide contig assembly. We used the *Oryzias latipes* HdrR (Japenese medaka; MEDAKA1), *Poecilla formosa* (Amazon molly, Poecilia_formosa-5.1.2), or *Xiphophorus maculatus* (platyfish, Xipmac4.4.2) reference genomes to guide exon concatenation. A total of 23,162 gene sequences were reconstructed for each species.

### CNE association with genes

Many CNEs function as tissue specific enhancers and promoters. As in Daane *et al*. [50], we used the Genomic Regions Enrichment of Annotations Tool (‘GREAT’) approach to assign CNEs to neighboring genes as putative regulatory targets [29]. This approach defines a basal regulatory window that is 5kb downstream and 1kb upstream of each gene’s transcription start site (TSS). This window is then extended up to 1Mb upstream and downstream of the TSS or until overlap the basal regulatory window of another gene. As opposed to simple nearest neighbor approaches, this approach allows CNEs to fall within the window of one or multiple genes, enhancing statistical power for gene ontology enrichment studies [29]. We based the regulatory windows in our analysis on the *Oryzias latipes* reference genome.

### Ontology annotations

To supplement the gene functional annotations for the Japense medaka, Amazon molly and platyfish orthologs we generated a merged gene ontology dataset from multiple species. We mined gene ontology annotations from human (*Homo sapiens*), mouse (*Mus musculus*), rat (*Rattus norvegicus*), chicken (*Gallus gallus*), three-spined stickleback (*Gasterosteus aculeatus*), medaka (*Oryzias latipes HdrR*), and zebrafish (*Danio rerio*) in Ensembl BioMart (downloaded January 2020) [67]. All Ensembl gene identifiers were then converted to a non-redundant set of medaka orthologs.

### Evolutionary sequence rate analysis

We calculated branch lengths for each gene along a fixed species tree topology using IQTree v1.7beta2 (parameters ‘–te-keep-ident-st DNA-m MFP’) [77]. We then used the program RERConverge v0.1.0 to estimate the relative evolutionary rates for each gene at all nodes in the phylogeny (parameters ‘transform = “sqrt”, weighted = T, scale = T, cutoff=0’)[87].

To track genome-wide patterns in relative evolutionary rate, we averaged the relative evolutionary rate across all genes in a given gene ontology term to generate a gene ontology-wide average relative evolutionary rate for each node in the phylogeny. We then compared the mean relative evolutionary rate between the gliding beloniform nodes and the other beloniform species using a Wilcoxon signed-rank test. P-values were corrected using FDR (Python module statsmodels v0.6.1; fdrcorrection0). To assess potential sampling bias and noise within the dataset, we calculated differences in relative evolutionary rate for all GO-terms across 500 random samplings of similar tip and ancestral node distribution as the gliding beloniforms (**S2 Fig**). Only 8 out of 500 samples showed any statistically significant difference in relative evolutionary rate for a GO-term, and all 8 samples had ≤4 significant terms (**S2 Fig**).

We also assessed accelerated or constrained sequence evolution along the ancestral branches of flying fishes and halfbeaks using the program phyloP (PHAST v1.4, parameters ‘--method LRT --noprune --features --mode ACC’)[88,89]. The tree model for phyloP was derived using phyloFit and the species tree (**Fig 1C**). For CNEs, the tree models were based on 6,874 elements of at least 200 bp and with ≥75% coverage across all species. For CDS, the phyloP tree models were derived from 2,718 reconstructed gene sequences of at least 750 bp and with ≥75% coverage across all species. We calculated gene ontology enrichment for accelerated or constrained evolution using a SUMSTAT approach as in Daub *et al*. [90]. Briefly, we normalized the log-likelihood ratio test score output from phyloP (ΔlnL) by taking a fourth root (ΔlnL4), and then summed these normalized ΔlnL4 scores across all genes in a given ontology term. P-values are estimated by boot-strap resampling (50,000 replicates) and corrected for multiple hypothesis testing using a Benjamini Hochberg false discovery rate procedure (FDR; python module statsmodels v.0.6.1; fdrcorrection0).

### Analysis of molecular convergence

To search for convergent amino acid substitutions, we looked for amino acid substitutions that were present within both lineages of flying halfbeaks (*Euleptorhamphus viridis* and *Oxyporhamphus micropterus*) and within ≥70% of flying fishes (Exocoetidae) but that were not observed in the other Beloniformes. We required coverage of the amino acid position in all immediate sister lineages to the gliding beloniforms (*Rhynchorhamphus georgii, Hemiramphus brasiliensis* and *Hemiramphus far*) and in at least 50% of the other beloniforms in the dataset. To assess background levels of convergent amino acid substitutions in the phylogeny, we calculated the number of identical amino acid substitutions observed in three topologically similar species groupings (**S8 Fig**). To account for coverage differences among species, we further normalized amino acid counts by the number of bases analyzed and by the total number of SNPs (convergent or not) that are unique to one or more of the foreground species and not found in the outgroups (**S6 Fig**). This SNP normalization helps control for differences in topology, whereby the relatedness of species may impact the likelihood of having unique SNPs not seen in outgroup lineages.

We assessed convergent evolutionary rates in gliding beloniforms using RERConverge v 0.1.0 with topology weighted correlations to treat each clade as a single observation during p-value calculation (parameters for getAllResiduals ‘transform = “sqrt”, weighted = T, scale = T, cutoff=0’; parameters for foreground2Tree ‘clade=“all”, weighted = TRUE’; parameters for correlateWithBinaryPhenotype ‘min.sp=10, min.pos=5, weighted=“auto”‘) [87]. P-values were corrected for multiple comparisons within RERConverge using the Benjamini-Hochberg procedure.

## Data Availability

The assembled contigs and annotations for Beloniformes will be made publicly available upon acceptance in the Zenodo repository (https://zenodo.org/). The raw sequencing data will be deposited in the NCBI database as a Bioproject. PacBio sequencing reads from a *lof* incross are available on the NCBI Sequence Read Archive (SRA; ERX1428166).

## Acknowledgements

This work was supported by a John Simon Guggenheim Fellowship, Milton Foundation Fund, and partially supported by NIH grant R01HD084985 to MPH, a National Science Foundation Doctoral Dissertation Improvement Grant (DDIG) to JMD (DEB-1407092), a German Research Foundation fellowship to NB (BL 1614/1-1), and an NSERC Discovery Grant to NRL. Specimen and tissue collection and curation was made possible by generous assistance from R. Pitman, L. Ballance, B. Collette, H.H. Tan, E. Lewallen, D. Bloom, M. Sabaj, J.A. Dorman, NOAA, the Scripps Institution of Oceanography, and the Royal Ontario Museum. The authors would like to acknowledge assistance by the Central Caribbean Marine Institute (CCMI) and their generous support of this research program. This work is dedicated to the insight and career of Dr. Steven Johnson who helped shape and identify the novel role of *kcnh2a* in regulating the longfin phenotype.

## Notes

### Competing Interest Statement

The authors have declared no competing interest.

## References

1. Nelson JS, Grande TC, Wilson MVH. Fishes of the World. Fifth Edit. Hoboken, NJ: John Wiley & Sons, Inc.; 2016.

2. Davenport J. How and why do flying fish fly? Rev Fish Biol Fish. 1994;4: 184–214.

3. Lewallen E a., Pitman RL, Kjartanson SL, Lovejoy NR. Molecular systematics of flyingfishes (Teleostei: Exocoetidae): evolution in the epipelagic zone. Biol J Linn Soc. 2011;102: 161–174. doi:10.1111/j.1095-8312.2010.01550.x

4. Dasilao JC, Sasaki K, Okamura O. The hemiramphid, Oxyporhamphus, is a flyingfish (Exocoetidae). Ichthyol Res. 1997;44: 101–107. doi:10.1007/BF02678688

5. Lovejoy NR, Iranpour M, Collette BB. Phylogeny and jaw ontogeny of beloniform fishes. Integr Comp Biol. 2004;44: 366–77. doi:10.1093/icb/44.5.366

6. Hughes LC, Ortí G, Huang Y, Sun Y, Baldwin CC, Thompson AW, et al. Comprehensive phylogeny of ray-finned fishes (Actinopterygii) based on transcriptomic and genomic data. Proc Natl Acad Sci U S A. 2018;115: 6249–6254. doi:10.1073/pnas.1719358115

7. Rabosky DL, Chang J, Title PO, Cowman PF, Sallan L, Friedman M, et al. An inverse latitudinal gradient in speciation rate for marine fishes. Nature. 2018;559: 392–395. doi:10.1038/s41586-018-0273-1

8. Betancur-R R, Wiley EO, Arratia G, Acero A, Bailly N, Miya M, et al. Phylogenetic classification of bony fishes. BMC Evol Biol. 2017;17: 162. doi:10.1186/s12862-017-0958-3

9. Daane JM, Rohner N, Konstantinidis P, Djuranovic S, Harris MP. Parallelism and Epistasis in Skeletal Evolution Identified through Use of Phylogenomic Mapping Strategies. Mol Biol Evol. 2015;33: 162–173. doi:10.1093/molbev/msv208

10. Daane JM, Dornburg A, Smits P, MacGuigan DJ, Brent Hawkins M, Near TJ, et al. Historical contingency shapes adaptive radiation in Antarctic fishes. Nat Ecol Evol. 2019;3: 1102–1109. doi:10.1038/s41559-019-0914-2

11. Lovejoy NR. REINTERPRETING RECAPITULATION: SYSTEMATICS OF NEEDLEFISHES AND THEIR ALLIES (TELEOSTEI: BELONIFORMES). Evolution (N Y). 2000;54: 1349–1362.

12. Kasumyan AO. The Vestibular System and Sense of Equilibrium in Fish. J Ichthyol. 2004;44: 224–268.

13. Dasilao J, Sasaki K. Phylogeny of the flyingfish family Exocoetidae. Ichthyol Res. 1998;45: 347–353.

14. Morris LS, McCall JG, Charney DS, Murrough JW. The role of the locus coeruleus in the generation of pathological anxiety. Brain Neurosci Adv. 2020;4: 239821282093032. doi:10.1177/2398212820930321

15. Baylor ER. Air and Water Vision of the Atlantic Flying Fish, Cypselurus heterurus. Nature. 1967;214: 307–309.

16. Harris MP, Daane JM, Lanni J. Through veiled mirrors: Fish fins giving insight into size regulation. WIREs Dev Biol. 2020; 1–16. doi:10.1002/wdev.381

17. van Eeden FJ, Granato M, Schach U, Brand M, Furutani-Seiki M, Haffter P, et al. Genetic analysis of fin formation in the zebrafish, Danio rerio. Development. 1996;123: 255–62. Available: http://www.ncbi.nlm.nih.gov/pubmed/9007245

18. Perathoner S, Daane JM, Henrion U, Seebohm G, Higdon CW, Johnson SL, et al. Bioelectric signaling regulates size in zebrafish fins. PLoS Genet. 2014;10: e1004080. doi:10.1371/journal.pgen.1004080

19. Iovine MK, Johnson SL. A genetic, deletion, physical, and human homology map of the long fin region on zebrafish linkage group 2. Genomics. 2002;79: 756–759. doi:10.1006/geno.2002.6769

20. Stewart S, Bleu HK Le, Yette GA, Henner AL, Robbins AE, Braunstein JA, et al. longfin causes cis-ectopic expression of the kcnh2a ether-a-go-go K+ channel to autonomously prolong fin outgrowth. bioRxiv. 2020; 1–46. doi:10.1101/790329

21. Bowen M, Henke K, Siegfried K, Warman ML, Harris MP. Efficient Mapping and Cloning of Mutations in Zebrafish by Low Coverage Whole Genome Sequencing. Genetics. 2011.

22. Bodoy S, Martín L, Zorzano A, Palacín M, Estévez R, Bertran J. Identification of LAT4, a novel amino acid transporter with system L activity. J Biol Chem. 2005;280: 12002–12011. doi:10.1074/jbc.M408638200

23. Guetg A, Mariotta L, Bock L, Herzog B, Fingerhut R, Camargo SMR, et al. Essential amino acid transporter Lat4 (Slc43a2) is required for mouse development. J Physiol. 2015;593: 1273–1289. doi:10.1113/jphysiol.2014.283960

24. Silic MR, Wu Q, Kim BH, Golling G, Chen KH, Freitas R, et al. Potassium channel-associated bioelectricity of the dermomyotome determines fin patterning in Zebrafish. Genetics. 2020;215: 1067–1084. doi:10.1534/genetics.120.303390

25. Schulte CJ, Allen C, England SJ, Juárez-Morales JL, Lewis KE. Evx1 is required for joint formation in zebrafish fin dermoskeleton. Dev Dyn. 2011;240: 1240–1248. doi:10.1002/dvdy.22534

26. Iovine MK, Higgins EP, Hindes A, Coblitz B, Johnson SL. Mutations in connexin43 (GJA1) perturb bone growth in zebrafish fins. Dev Biol. 2005;278: 208–19. doi:10.1016/j.ydbio.2004.11.005

27. Sims K, Eble DM, Iovine MK. Connexin43 regulates joint location in zebrafish fins. Dev Biol. 2009;327: 410–418. doi:10.1016/j.ydbio.2008.12.027

28. Maden M. Vitamin A and pattern formation in the regenerating limb. Nature. 1982;295: 672–675. doi:10.1038/295672a0

29. McLean CY, Bristor D, Hiller M, Clarke SL, Schaar BT, Lowe CB, et al. GREAT improves functional interpretation of cis-regulatory regions. Nat Biotechnol. 2010;28: 495–501. doi:10.1038/nbt.1630

30. Harvey SA, Logan MPO. Sall4 acts downstream of tbx5 and is required for pectoral fin outgrowth. Development. 2006;133: 1165–1173. doi:10.1242/dev.02259

31. Kawakami Y, Uchiyama Y, Esteban CR, Inenaga T, Koyano-Nakagawa N, Kawakami H, et al. Sall genes regulate region-specific morphogenesis in the mouse limb by modulating Hox activities. Development. 2009;136: 585–594. doi:10.1242/dev.027748

32. Young JJ, Grayson P, Edwards S V., Tabin CJ. Attenuated Fgf Signaling Underlies the Forelimb Heterochrony in the Emu Dromaius novaehollandiae. Curr Biol. 2019;29: 3681–3691.e5. doi:10.1016/j.cub.2019.09.014

33. Schartl M, Kneitz S, Ormanns J, Schmidt C, Anderson JL, Amores A, et al. The Developmental and Genetic Architecture of the Sexually Selected Male Ornament of Swordtails. Curr Biol. 2021;31: 1–12. doi:10.1016/j.cub.2020.11.028

34. Lanni JS, Peal D, Ekstrom L, Chen H, Stanclift C, Bowen ME, et al. Integrated K+ channel and K+Cl-cotransporter functions are required for the coordination of size and proportion during development. Dev Biol. 2019;456: 164–178. doi:10.1016/j.ydbio.2019.08.016

35. Nicklin P, Bergman P, Zhang B, Triantafellow E, Wang H, Nyfeler B, et al. Bidirectional Transport of Amino Acids Regulates mTOR and Autophagy. Cell. 2009;136: 521–534. doi:10.1016/j.cell.2008.11.044

36. Guetg A, Mariotta L, Bock L, Herzog B, Fingerhut R, Camargo SMR, et al. Essential amino acid transporter Lat4 (Slc43a2) is required for mouse development. J Physiol. 2015;5: 1273–1289. doi:10.1113/jphysiol.2014.283960

37. Haase C, Bergmann R, Fuechtner F, Hoepping A, Pietzsch J. L-type amino acid transporters LAT1 and LAT4 in cancer: uptake of 3-O-methyl-6-18F-fluoro-L-dopa in human adenocarcinoma and squamous cell carcinoma in vitro and in vivo. J Nucl Med. 2007;48: 2063–2071. doi:10.2967/jnumed.107.043620

38. Wang Q, Holst J. L-type amino acid transport and cancer: targeting the mTORC1 pathway to inhibit neoplasia. Am J Cancer Res. 2015;5: 1281–94. Available: http://www.pubmedcentral.nih.gov/articlerender.fcgi?artid=4473310&tool=pmcentrez&rendertype=abstract

39. Sutter B, Wu X, Laxman S, BP T. Methionine Inhibits Autophagy and Promotes Growth by Inducing the SAM-Responsive Methylation of PP2A. Cell. 2013;154: 403–415. doi:10.1016/j.cell.2013.06.041.Methionine

40. Goldsmith MI, Fisher S, Waterman R, Johnson SL. Saltatory control of isometric growth in the zebrafish caudal fin is disrupted in long fin and rapunzel mutants. Dev Biol. 2003;259: 303–317. doi:10.1016/S0012-1606(03)00186-0

41. Green J, Taylor JJ, Hindes A, Johnson SL, Goldsmith MI. A gain of function mutation causing skeletal overgrowth in the rapunzel mutant. Dev Biol. 2009;334: 224–34. doi:10.1016/j.ydbio.2009.07.025

42. Daane JM, Lanni J, Rothenberg I, Seebohm G, Higdon CW, Johnson SL, et al. Bioelectric-calcineurin signaling module regulates allometric growth and size of the zebrafish fin. Sci Rep. 2018;8: 10391. doi:10.1038/s41598-018-28450-6

43. Zou Z, Zhang J. Are Convergent and Parallel Amino Acid Substitutions in Protein Evolution More Prevalent Than Neutral Expectations? Mol Biol Evol. 2015;32: 2085–2096. doi:10.1093/molbev/msv091

44. Stern DL. The genetic causes of convergent evolution. Nat Rev Genet. 2013;14: 751–64. doi:10.1038/nrg3483

45. Zou Z, Zhang J. No genome-wide protein sequence convergence for echolocation. Mol Biol Evol. 2015;32: 1237–1241. doi:10.1093/molbev/msv014

46. Thomas GWC, Hahn MW. Determining the null model for detecting adaptive convergence from genomic data: A case study using echolocating mammals. Mol Biol Evol. 2015;32: 1232–1236. doi:10.1093/molbev/msv013

47. Marcovitz A, Turakhia Y, Chen HI, Gloudemans M, Braun BA, Wang H, et al. A functional enrichment test for molecular convergent evolution finds a clear protein-coding signal in echolocating bats and whales. Proc Natl Acad Sci. 2019;116: 21094–21103. doi:10.1073/pnas.1818532116

48. Xu G, Zhao L, Gao K, Wu F. A new stem-neopterygian fish from the Middle Triassic of China shows the earliest over-water gliding strategy of the A new stem-neopterygian fish from the Middle Triassic of China shows the earliest over-water gliding strategy of the vertebrates. Proc R Soc B Biol Sci. 2013;280: 20122261.

49. Tintori A, Sassi D. Thoracopterus Bronn (Osteichthyes: Actinopterygii): A gliding fish from the Upper Triassic of Europe. J Vertebr Paleontol. 1992;12: 265–283. doi:10.1080/02724634.1992.10011459

50. Daane JM, Auvinet J, Stoebenau A, Yergeau D, Harris MP, Detrich HW. Developmental constraint shaped genome evolution and erythrocyte loss in Antarctic fishes following paleoclimate change. PLoS Genet. 2020;16: 1–22. doi:10.1371/journal.pgen.1009173

51. Feng S, Stiller J, Deng Y, Armstrong J, Fang Q, Reeve AH, et al. Dense sampling of bird diversity increases power of comparative genomics. Nature. 2020;587: 252–257. doi:10.1038/s41586-020-2873-9

52. Lovejoy NR, Collette BB. Phylogenetic Relationships of New World Needlefishes (Teleostei: Belonidae) and the Biogeography of Transitions between Marine and Freshwater Habitats. Copeia. 2001;2: 324–338.

53. de Beer G. Embryos and Ancestors. Oxford University Press; 1940. doi:10.1038/148545a0

54. Gould SJ. Ontogeny and phylogeny. Harvard University Press; 1977.

55. Nüsslein-Volhard C, Dahm R. Zebrafish: a practical approach. Oxford University Press; 2002. doi:10.1017/S0016672303216384

56. Rohner N, Perathoner S, Frohnhöfer HG, Harris MP. Enhancing the Efficiency of N-Ethyl-N-Nitrosourea-Induced Mutagenesis in the Zebrafish. Zebrafish. 2011;8: 10–15. doi:10.1089/zeb.2011.0703

57. Fritz A, Rozowski M, Walker C, Westerfield M. Identification of selected gamma-ray induced deficiencies in zebrafish using multiplex polymerase chain reaction. Genetics. 1996;144: 1735–1745.

58. Koren S, Walenz BP, Berlin K, Miller JR, Bergman NH, Phillippy AM. Canu: Scalable and accurate long-read assembly via adaptive κ-mer weighting and repeat separation. Genome Res. 2017;27: 722–736. doi:10.1101/gr.215087.116

59. Bowen ME, Henke K, Siegfried KR, Warman ML, Harris MP. Efficient mapping and cloning of mutations in zebrafish by low-coverage whole-genome sequencing. Genetics. 2012;190: 1017–24. doi:10.1534/genetics.111.136069

60. Bowen ME, Boyden ED, Holm IA, Campos-Xavier B, Bonafé L, Superti-Furga A, et al. Loss-of-function mutations in PTPN11 cause metachondromatosis, but not Ollier disease or Maffucci syndrome. PLoS Genet. 2011;7: e1002050. doi:10.1371/journal.pgen.1002050

61. Montague TG, Cruz JM, Gagnon JA, Church GM, Valen E. CHOPCHOP: A CRISPR/Cas9 and TALEN web tool for genome editing. Nucleic Acids Res. 2014;42: 401–407. doi:10.1093/nar/gku410

62. Labun K, Montague TG, Gagnon JA, Thyme SB, Valen E. CHOPCHOP v2: a web tool for the next generation of CRISPR genome engineering. Nucleic Acids Res. 2016;44: gkw398. doi:10.1093/nar/gkw398

63. Gagnon J a, Valen E, Thyme SB, Huang P, Ahkmetova L, Pauli A, et al. Efficient mutagenesis by Cas9 protein-mediated oligonucleotide insertion and large-scale assessment of single-guide RNAs. PLoS One. 2014;9: e98186. doi:10.1371/journal.pone.0098186

64. Hwang WY, Fu Y, Reyon D, Maeder ML, Tsai SQ, Sander JD, et al. Efficient genome editing in zebrafish using a CRISPR-Cas system. Nat Biotechnol. 2013;31: 227–229. doi:10.1038/nbt.2501

65. Hwang WY, Fu Y, Reyon D, Maeder ML, Kaini P, Sander JD, et al. Heritable and precise zebrafish genome editing using a CRISPR-Cas system. PLoS One. 2013;8: e68708. doi:10.1371/journal.pone.0068708

66. Fu Y, Foden J a, Khayter C, Maeder ML, Reyon D, Joung JK, et al. High-frequency off-target mutagenesis induced by CRISPR-Cas nucleases in human cells. Nat Biotechnol. 2013;31: 822–6. doi:10.1038/nbt.2623

67. Kinsella RJ, Kähäri A, Haider S, Zamora J, Proctor G, Spudich G, et al. Ensembl BioMarts: A hub for data retrieval across taxonomic space. Database. 2011;2011: 1–9. doi:10.1093/database/bar030

68. Herrero J, Muffato M, Beal K, Fitzgerald S, Gordon L, Pignatelli M, et al. Ensembl comparative genomics resources. Database. 2016;2016: 1–17. doi:10.1093/database/bav096

69. Kozomara A, Griffiths-Jones S. MiRBase: Integrating microRNA annotation and deepsequencing data. Nucleic Acids Res. 2011;39: 1–6. doi:10.1093/nar/gkq1027

70. Dimitrieva S, Bucher P. UCNEbase - A database of ultraconserved non-coding elements and genomic regulatory blocks. Nucleic Acids Res. 2013;41: 101–109. doi:10.1093/nar/gks1092

71. Quinlan AR, Hall IM. BEDTools: a flexible suite of utilities for comparing genomic features. Bioinformatics. 2010;26: 841–2. doi:10.1093/bioinformatics/btq033

72. Bolger AM, Lohse M, Usadel B. Trimmomatic: A flexible trimmer for Illumina sequence data. Bioinformatics. 2014;30: 2114–2120. doi:10.1093/bioinformatics/btu170

73. Huang X. CAP3: A DNA Sequence Assembly Program. Genome Res. 1999;9: 868–877. doi:10.1101/gr.9.9.868

74. Edgar RC. Search and clustering orders of magnitude faster than BLAST. Bioinformatics. 2010;26: 2460–2461. doi:10.1093/bioinformatics/btq461

75. Sedlazeck FJ, Rescheneder P, Von Haeseler A. NextGenMap: Fast and accurate read mapping in highly polymorphic genomes. Bioinformatics. 2013;29: 2790–2791. doi:10.1093/bioinformatics/btt468

76. Katoh K, Standley DM. MAFFT multiple sequence alignment software version 7: improvements in performance and usability. Mol Biol Evol. 2013;30: 772–80. doi:10.1093/molbev/mst010

77. Nguyen LT, Schmidt HA, Von Haeseler A, Minh BQ. IQ-TREE: A fast and effective stochastic algorithm for estimating maximum-likelihood phylogenies. Mol Biol Evol. 2015;32: 268–274. doi:10.1093/molbev/msu300

78. Li H, Handsaker B, Wysoker A, Fennell T, Ruan J, Homer N, et al. The Sequence Alignment/Map format and SAMtools. Bioinformatics. 2009;25: 2078–9. doi:10.1093/bioinformatics/btp352

79. Robinson JT, Thorvaldsdóttir H, Winckler W, Guttman M, Lander ES, Getz G, et al. Integrative Genome Viewer. Nat Biotechnol. 2011;29: 24–6. doi:10.1038/nbt.1754.Integrative

80. Kofler R, Orozco-terWengel P, De Maio N, Pandey RV, Nolte V, Futschik A, et al. PoPoolation: a toolbox for population genetic analysis of next generation sequencing data from pooled individuals. PLoS One. 2011;6: e15925. doi:10.1371/journal.pone.0015925

81. Ranwez V, Harispe S, Delsuc F, Douzery EJP. MACSE: Multiple Alignment of Coding SEquences Accounting for Frameshifts and Stop Codons. Murphy WJ, editor. PLoS One. 2011;6: e22594. doi:10.1371/journal.pone.0022594

82. Borowiec M. Spruceup: fast and flexible identification, visualization, and removal of outliers from large multiple sequence alignments. J Open Source Softw. 2019;4: 1635. doi:10.21105/joss.01635

83. Kalyaanamoorthy S, Minh BQ, Wong TKF, Von Haeseler A, Jermiin LS. ModelFinder: Fast model selection for accurate phylogenetic estimates. Nat Methods. 2017;14: 587–589. doi:10.1038/nmeth.4285

84. Hoang DT, Chernomor O, Von Haeseler A, Minh BQ, Vinh LS. UFBoot2: Improving the ultrafast bootstrap approximation. Mol Biol Evol. 2018;35: 518–522. doi:10.1093/molbev/msx281

85. Zhang C, Rabiee M, Sayyari E, Mirarab S. ASTRAL-III: Polynomial time species tree reconstruction from partially resolved gene trees. BMC Bioinformatics. 2018;19: 15–30. doi:10.1186/s12859-018-2129-y

86. Sayyari E, Mirarab S. Fast Coalescent-Based Computation of Local Branch Support from Quartet Frequencies. Mol Biol Evol. 2016;33: 1654–1668. doi:10.1093/molbev/msw079

87. Kowalczyk A, Meyer WK, Partha R, Mao W, Clark NL, Chikina M. RERconverge: an R package for associating evolutionary rates with convergent traits. Valencia A, editor. Bioinformatics. 2019;35: 4815–4817. doi:10.1093/bioinformatics/btz468

88. Pollard KS, Hubisz MJ, Rosenbloom KR, Siepel A. Detection of nonneutral substitution rates on mammalian phylogenies. Genome Res. 2010;20: 110–21. doi:10.1101/gr.097857.109

89. Hubisz MJ, Pollard KS, Siepel A. PHAST and RPHAST: phylogenetic analysis with space/time models. Brief Bioinform. 2011;12: 41–51. doi:10.1093/bib/bbq072

90. Daub JT, Moretti S, Davydov II, Excoffier L. Detection of Pathways Affected by Positive Selection in Primate Lineages Ancestral to Humans. Mol Biol Evol. 2017;34: 1391–1402. doi:10.1093/molbev/msx083

