## Supplementary Information for "Novel regulators of growth identified in the evolution of fin proportion in flying fish"

**This PDF file includes:**

Supplementary Figures 1 to 11

Supplementary Tables 1 to 5, 7 and 8, 10 to 13

### Table of Contents

**Figures**

S1 Fig 3
S2 Fig 4
S3 Fig 5
S4 Fig 6
S5 Fig 7
S6 Fig 8
S7 Fig 9
S8 Fig 10
S9 Fig 11
S10 Fig 12
S11 Fig 13

**Tables**

S1 Table 19
S2 Table 22
S3 Table 23
S4 Table 28
S5 Table 29
S7 Table 30
S8 Table 31
S10 Table 32
S11 Table 33
S12 Table 41
S13 Table 42

#### S1 Figure

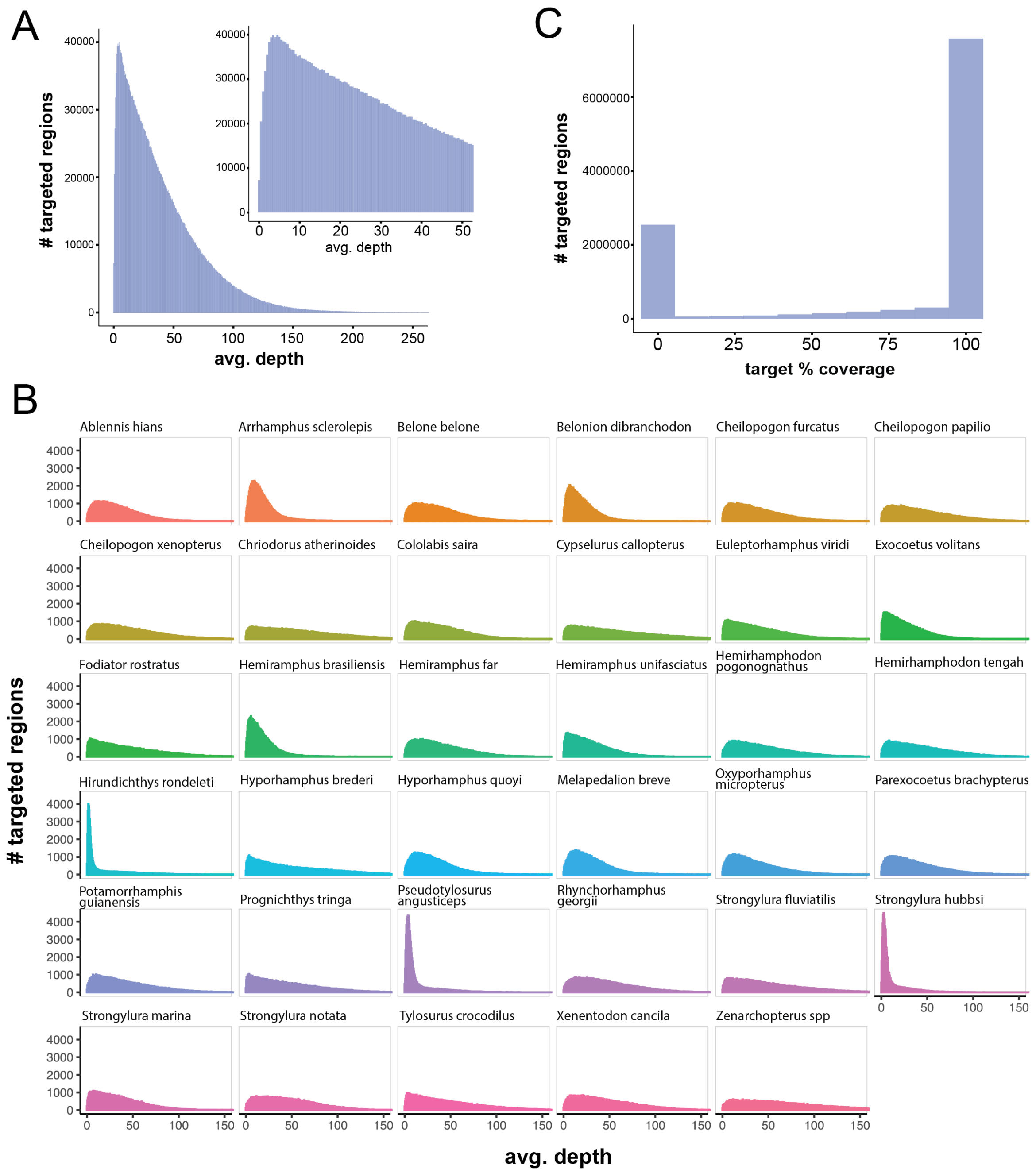


**S1 Fig. Sequencing read coverage and depth across the beloniform dataset**. (**A**) Histogram of the average sequencing depth from targeted sequence enrichment for every targeted element (for example, exon, CNE) and across all sequenced species. Each species has coverage in an average of 270,000 targeted elements. Inset, magnification of lower values (<50x depth). (**B**) Distribution of sequencing depth per targeted sequence element for each species. (**C**) Distribution of read coverage across targeted regions. In general, most targeted elements had full coverage or were absent from the analysis.

#### S2 Figure

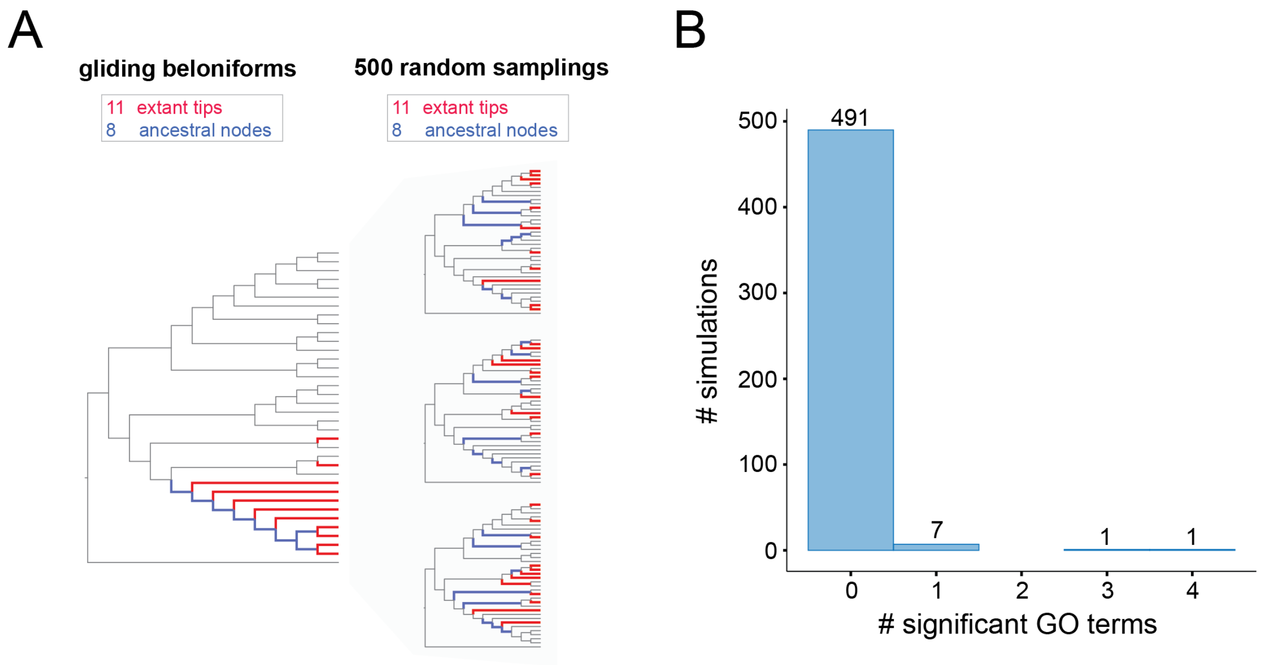


**S2 Fig**. **Simulating branch topology to assess background levels of divergence in evolutionary rate across gene ontology groupings**. (**A**) Beloniformes species tree with extant gliding lineages (red) and ancestral gliding lineages (blue) highlighted. 500 random species samplings of similar tip and ancestral node distribution were generated. For each species sampling, the difference between the average relative evolutionary rate across each GO-term were compared between the selected branches and the background branches. (**B**) Number of significant differences between selected branches and background branches across all GO-terms and all simulations. Significance was determined by Wilcoxon signed-rank test and p-values corrected by Benjamini-Hochberg procedure. As opposed to the analysis of gliding beloniforms (**Fig 3**), random samplings of branches rarely generated significantly divergent relative evolutionary rates between GO-terms.

#### S3 Figure

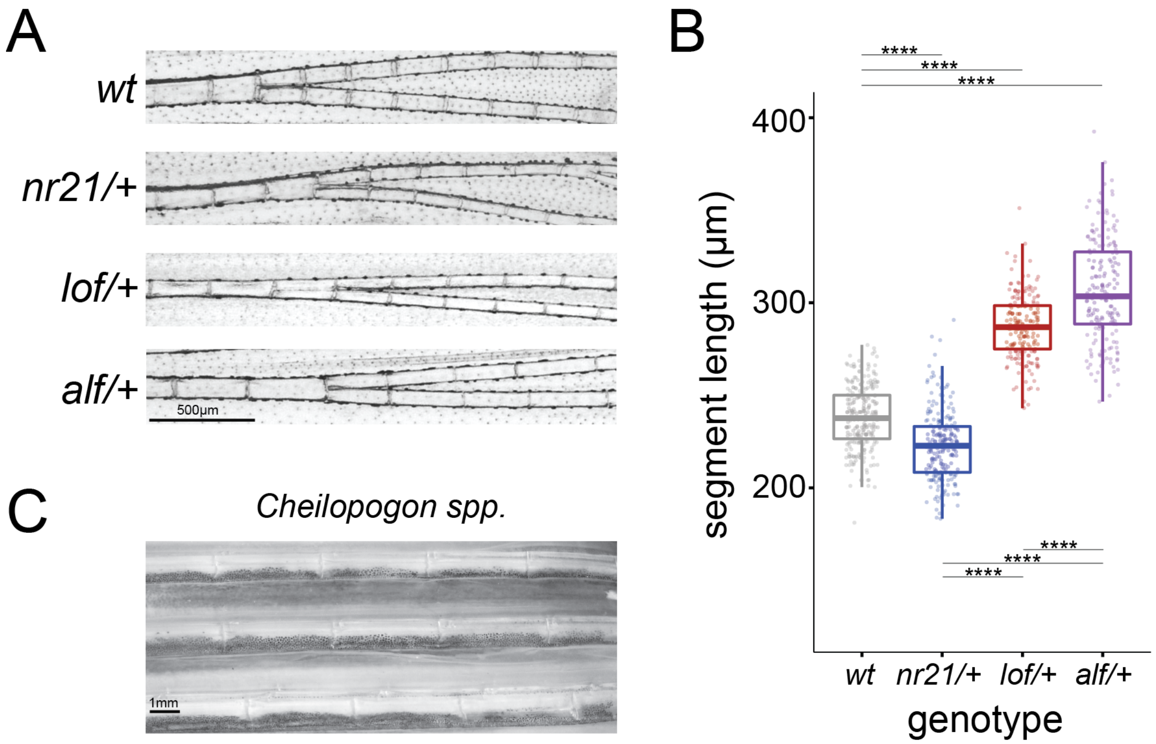


**S3 Fig. Lepidotrichia segment patterning in zebrafish fin mutants**. (**A**) Images of pectoral fin lepidotrichia segments in wildtype (*wt*), short finned (*nr21*) and long finned (*lof*, *alf*) zebrafish mutants. (**B**) Segment length across pectoral fins in zebrafish mutants. Roughly 30 segments were measured per fin across n=8 individuals for *wt* and *nr21* and from n=6 individuals for *lof* and *alf*. Note variable segment length in *alf* fins. **** indicates Tukey HSD adjusted p-value <0.0001. (**C**) Pectoral fin lepidotrichia from a flying fish, *Cheilopogon spp*.

### S4 Figure

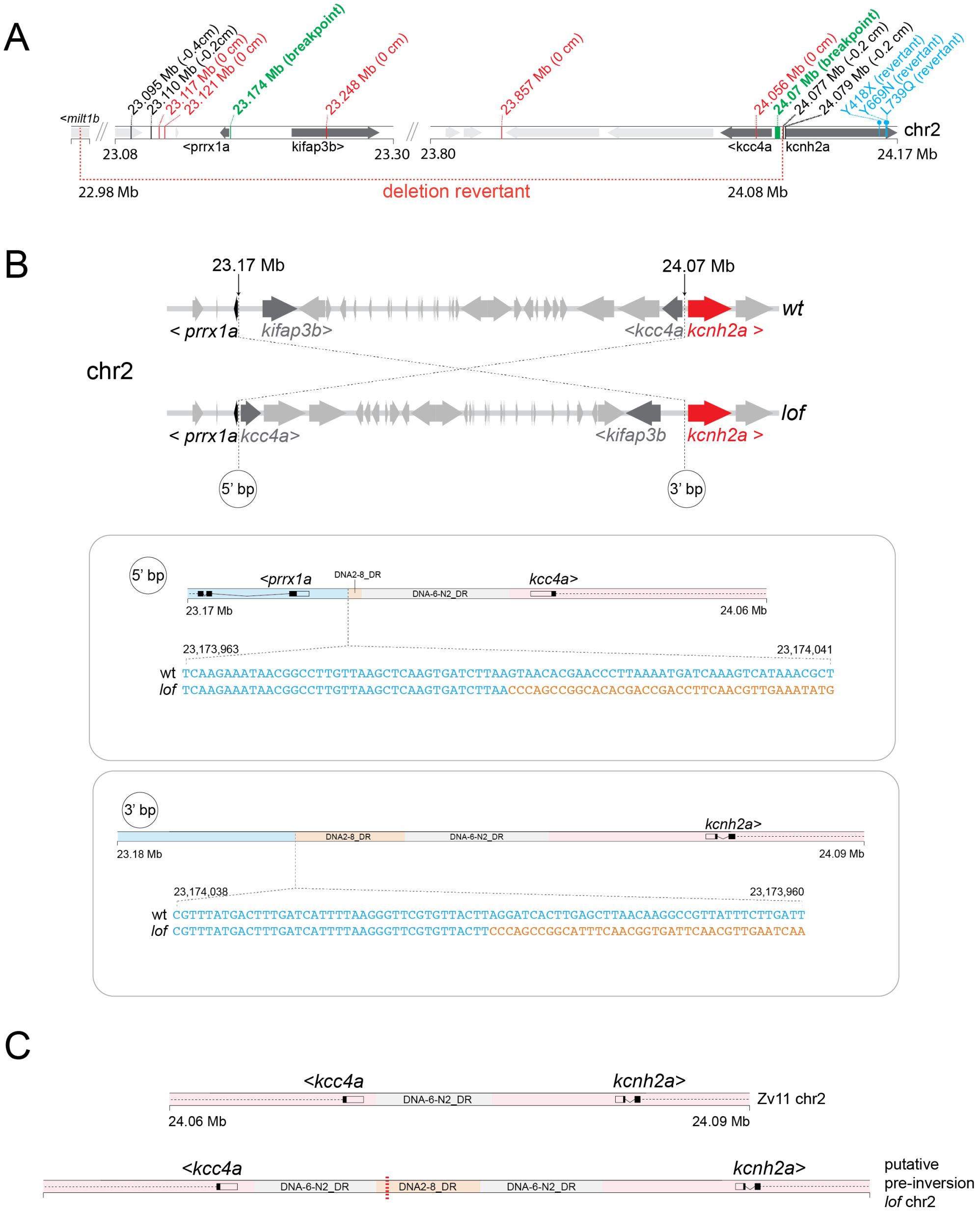


**S4 Fig. Mapping of the *longfin* zebrafish ($lof^{dt2}$).** (**A**) Combined recombinant and revertant data for $lof^{dt2}$. Linkage mapping identified a large region (~1 Mb) on chromosome 2, without recovery of additional recombinants. Four total reversion alleles were isolated, three point mutations in *kcnh2a* (Y418X, Y669N, L739Q) and a large deletion between 22.98 Mb and 24.08 Mb, just upstream of *kcnh2a* ($lof^{j6e1}$). (**B**) Local assembly of PacBio sequencing reads identifies inversion breakpoints (bp) upstream of *prrx1a* and upstream of *kcnh2a*. (**C**) Reconstructed ancestral $lof^{dt2}$ sequence prior to inversion, showing hypothetical transposon expansion relative to the zebrafish reference genome from the Tübingen strain (Zv11). Coordinates represent Zv11 positions.

#### S5 Figure

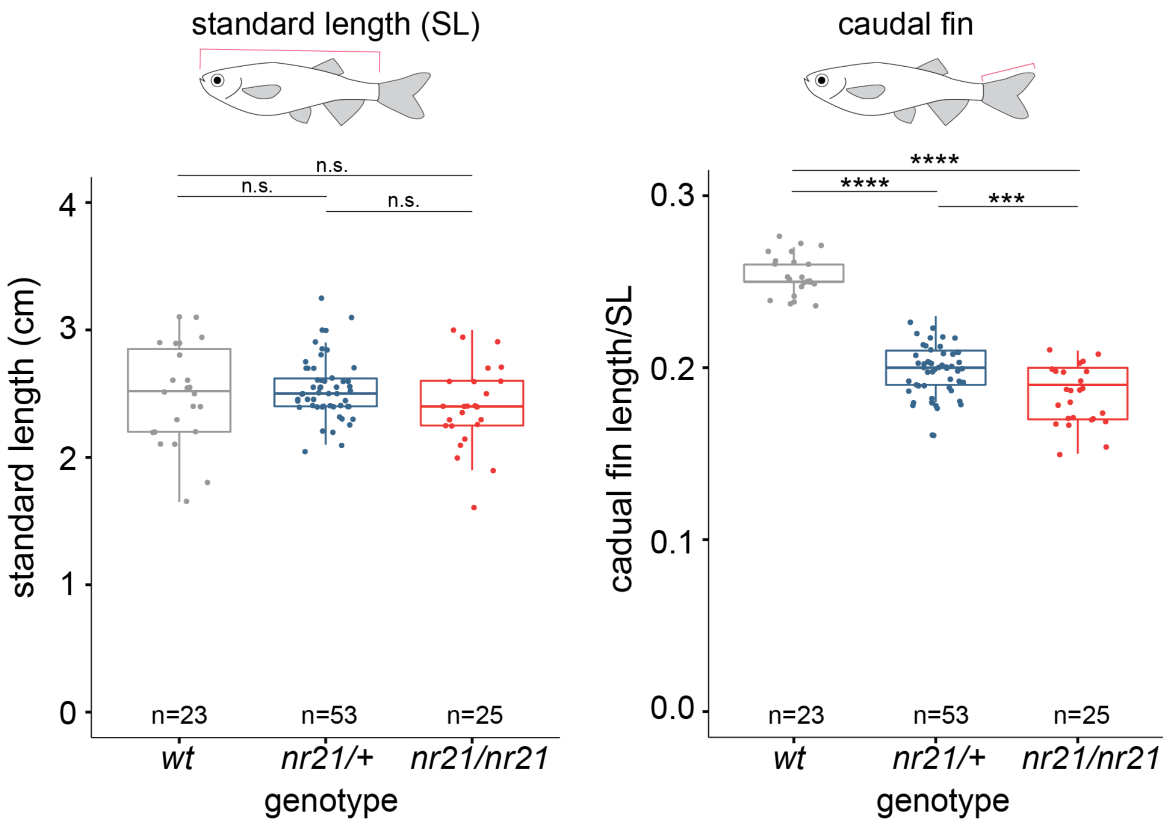


**S5 Fig. Impact of *nr21* mutation on body size and fin growth.** Comparison of fish standard length (SL) and caudal fin length in wildtype, heterozygous (*nr21/+*) and homozygous (*nr21/nr21*) individuals. p-values generated through Tukey's HSD. **** adjusted p-value ≤ 0.0001, *** adjusted p-value ≤0.001. n.s. indicates not significant.

#### S6 Figure

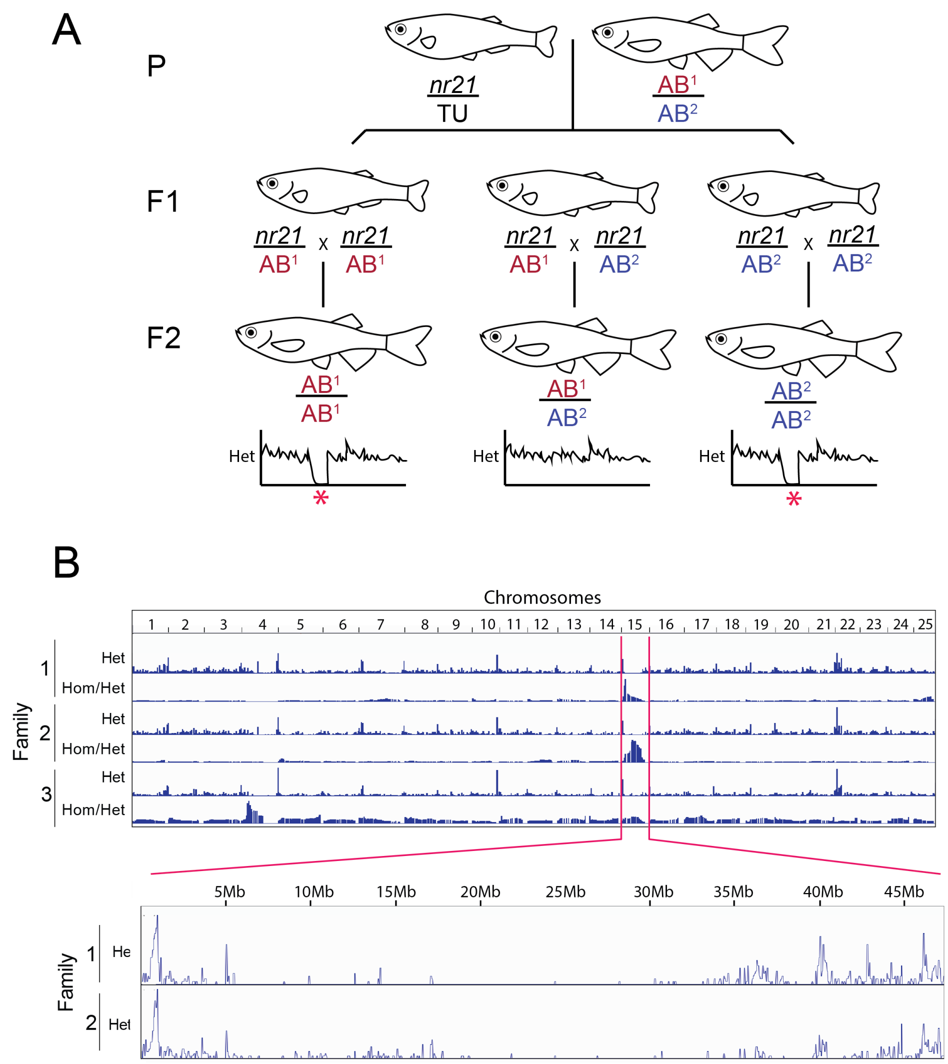


**S6 Fig. Mapping strategy for the shortfin zebrafish mutant, *nr21*.** (**A**) Mapping strategy for identifying *nr21* mutation. Mapping-by-heterozygosity was used to identify the mutant chromosome [59]. As *nr21* is dominant, mapping was performed on F2 sibling pools for the wildtype (recessive) phenotype. However, given SNP diversity in wildtype zebrafish strains, three separate options were possible depending on the F1 parental cross. Some of the F1 crosses would be homogeneous for particular wildtype haplotype (AB1/AB1 or AB2/AB2), but others may be heterozygous at the locus (AB1/AB2), obfuscating the mapping signal. (**B**) Genome-wide patterns of heterozygosity and the ratio of heterozygous to homozygous SNPs along a 15 cenitmorgan (cM) sliding window. Of the three wildtype families, two families showed a strong ratio homozygosity/heterozygosity on chromosome 15 between 17-33Mb. Within this interval, three non-synonymous SNPs (*med13b*: Q1531H, *lat4a*: T200K, *si:dkey-285b23.3*: E354G) were identified within sequence from nr21 carriers. There were 0 recombinants among 145 chromosomes for the *lat4a* T200K SNP.

**S7 Figure**

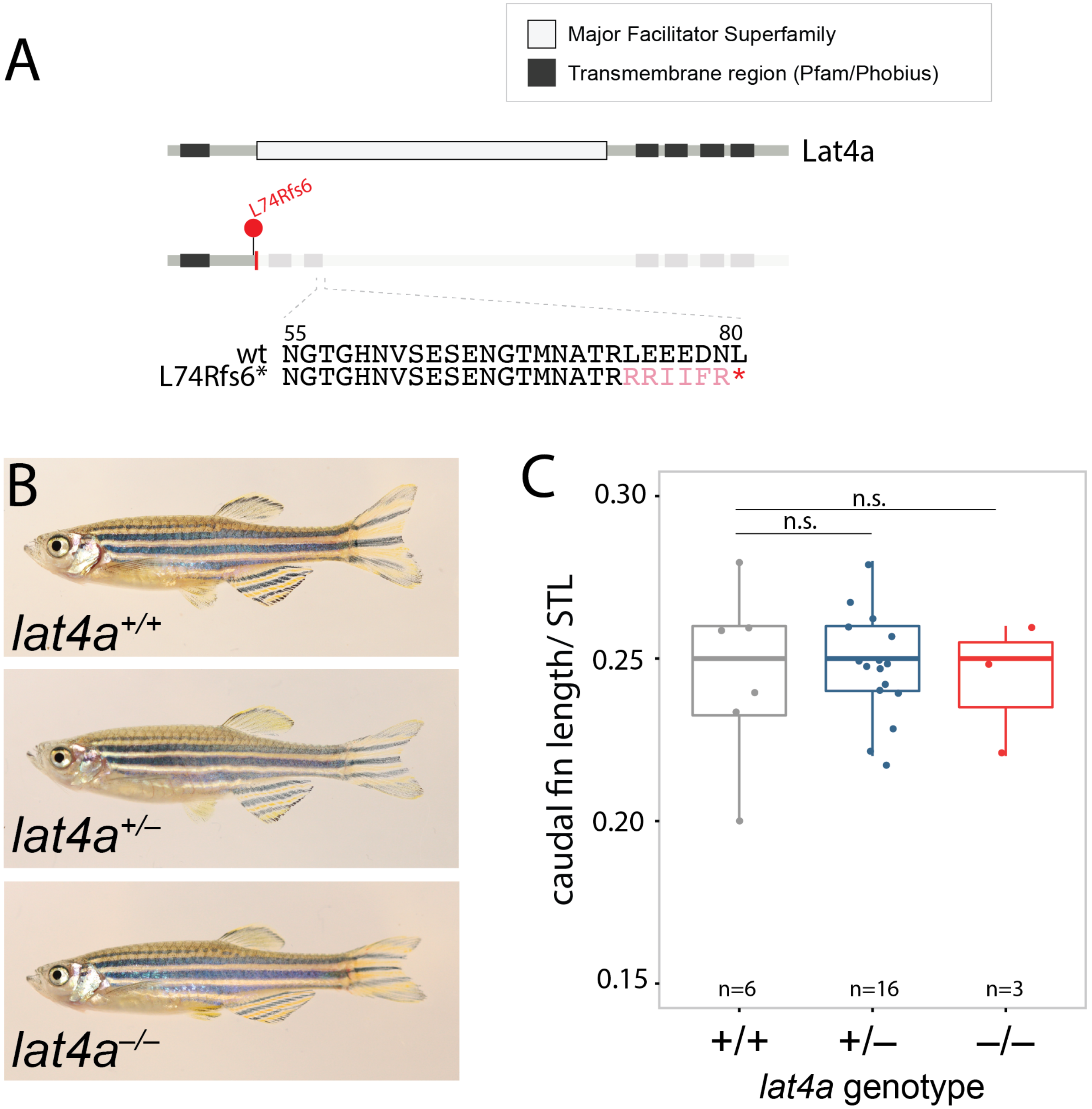


**S7 Fig. *lat4a* is not required for normal fin growth.** (**A**) Truncating frameshift allele (L74Rfs6) in *lat4a* generated through CRISPR/Cas9. (**B**) Images of wildtype, heterozygous and homozygous *lat4a* knockouts with wildtype fin patterning and no obvious phenotype. (**C**) Measurements of caudal fin length normalized to fish standard length (STL) showing no significant difference in fin size in the absence of *lat4a* (Tukey's HSD)

#### S8 Figure

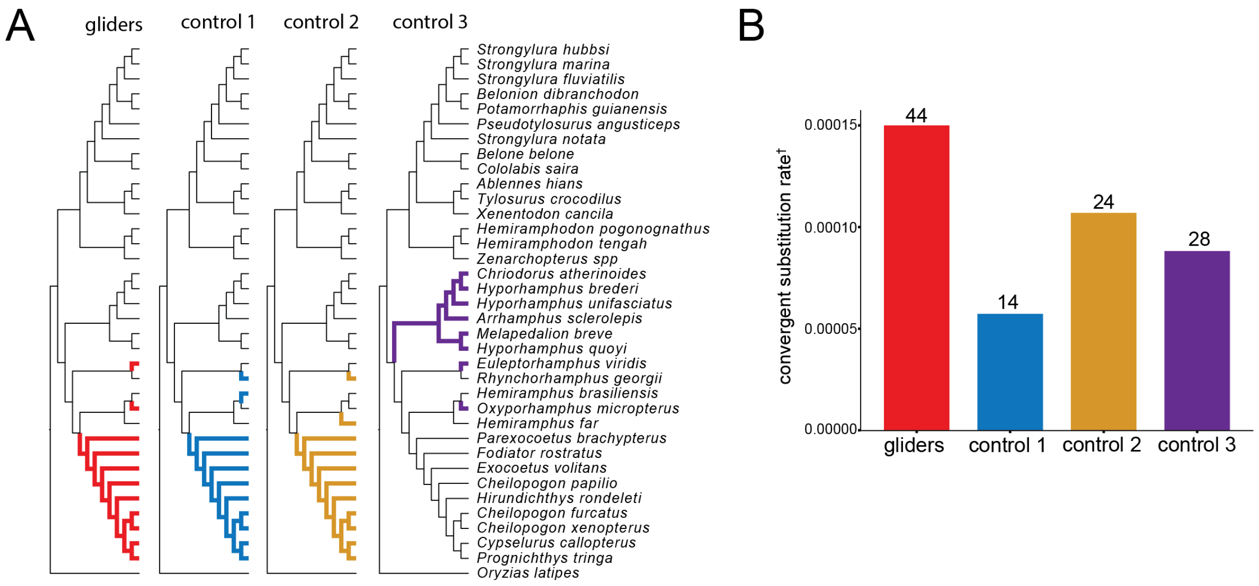


**S8 Fig. Comparison of convergent amino acid substitutions across gliding beloniforms and control species samplings**. (**A**) Highlighted branches for the gliding beloniforms (red), and three topologically similar control groupings of flying fishes and halfbeaks (blue, yellow, purple). (**B**) The number of detected identical amino acid substitutions found in >70% of the species in each of the three clade selections. Amino acid substitutions were ignored if also observed in background branches. Convergent substitution rate (†) refers to the number of identical amino acid substitutions found in all three highlighted clades normalized by the number of total unique substitutions found within at least one of the three clade selections and not within background branches. This normalizes the data to the number of bases analyzed in the multiple sequence alignment and the topological distances between species, which may increase or decrease the likelihood of having unique SNPs not seen in background branches. The number above each bar indicates total number of convergent amino acids. Convergent substitutions in gliders can be found in **S10 Table**.

#### S9 Figure

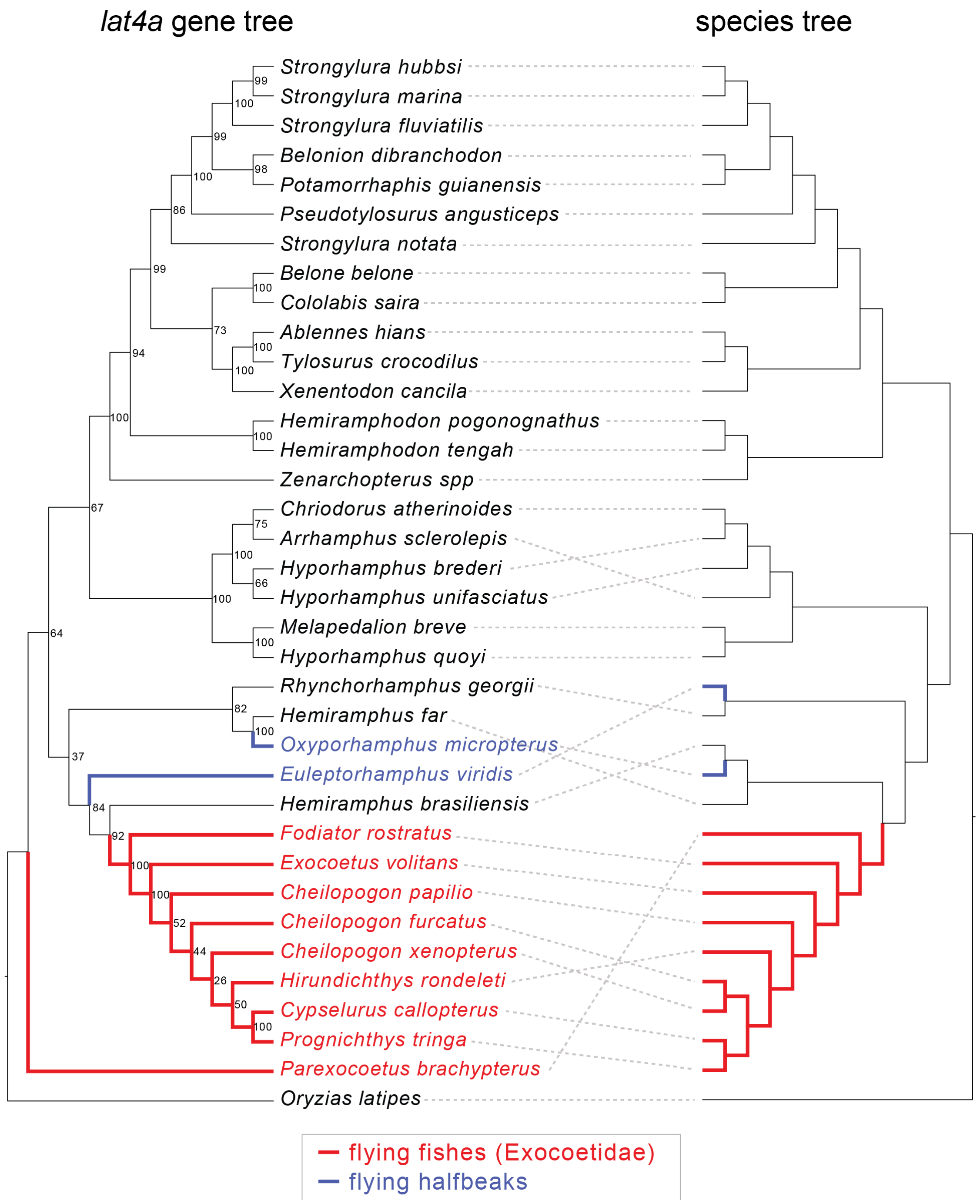


**S9 Fig. Comparison of *lat4a* gene tree and species tree**. The gliding beloniforms are specifically highlighted, with the flying fish branches (Exocoetidae) in red, and flying halfbeaks in blue. Maximum likelihood gene tree was estimated by IQTree. Node labels indicate ultrafast bootstrap support values. Species tree generated from 4,683 gene trees using ASTRAL (**Fig 1**).

#### S10 Figure

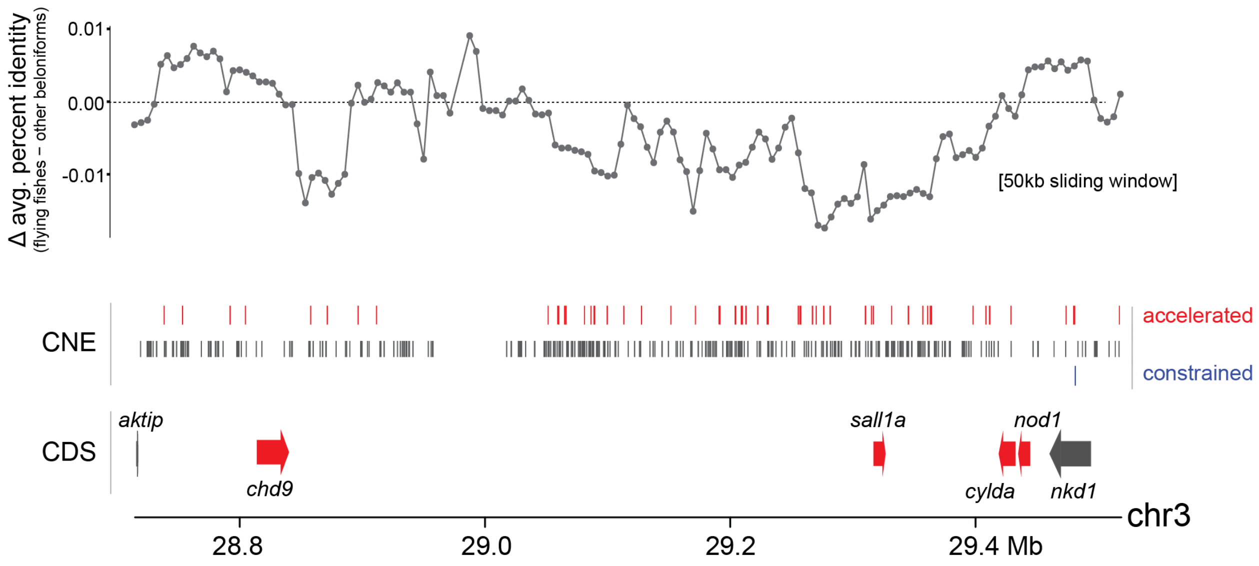


**S10 Fig**. **Accelerated sequence evolution at the sall1a locus in the common ancestor to flying fishes**. Top panel, 50kb genomic sliding window of average percent identity to the Japanese medaka genome (*O. latipes*) across targeted elements (CDS and CNE) in flying fishes compared to the other beloniforms. Low values indicate an increase in mutations in flying fishes relative to outgroups. CNEs and CDS considered by the program phyloP to be under accelerated sequence evolution (red), neutral (gray) or under constraint (blue) along the ancestral branch of flying fishes.

#### S11 Figure – Breakpoint contigs from PacBio assembly

##### --PacBio contig1--

**-Prrx1a side (forward strand) – 5’ breakpoint ~ 2:23,174,000 (Zv11)**
-Middle piece of DNA2-8_DR – 3’ breakpoint inside of this transposon
**-Kcc4a side (reverse strand) (transposon underlined – DNA-6-N2_DR)**

```
CAGTTATTATATATATACATTTTTATTTCCAACCTTTGACAAGTTAATTTGTGTCATAAAATACTTTGCACATTTTTCGTTATGACATGATATGTTATAA
ACATTTATTCACAAGTACATTGCATTCACATTTTAGTTAATATGCTAAAGTTGCTGTCTAATATCATACATTTATGAATAACCTATACAATATCCAGCAC
TACAAATATTCCCTCTGATGAAATAATGGTGAATTTAGGCAGAGTCATATAAAAAGTGTTGGATTTCCTACCTGTATGACGGTGAGCTTCCCCAGGACAG
GTAGTCATTGGGTCTGGGTGCGGGTCGTGGACAATGGGCTGCTCTACGGCTGTTACTTCTCCTGAGAAAGACTTGAGCAATGAAGCATTTTTACTTGCCA
GCATCGCACGTTTCATTTCGTCGAAACTTAGCCCGTCGATTTTGAAACCATACCTGGAAATTAAAGAAAGGTGTTGAGCTTGTCATGCATAGATGGAGAT
CTATCATGATGGAGACCATTAGAAAGGAGAAGGTCAAACACAGGCTAATACAAATCTCTAACCTGAACTCGTGCTTCAGTGAGGTTGACCCGTCGTGCCA
AGTCCTCCCTGACGAAAGCATCGGGGTAATGCGTCCTCTCAAACACTCTCTCTAAGGCCTGCAGCTGACTGCTGTTGAACGTGGTTCTGTTTCGCCTCTG
TTTACGCTTCTTCTTCTCCTCTGCGTTGAGCTGCTCGTCTGGAGGCGCACAAATAGTCAAATATTAATATGACGTGAATATCCAAATTATAATGTGGGTC
AGATTCTTGCACTGTAAAAAGGCTGGCTTCCACAGCCTGGATTTGTGTTGAGACTACATGAAGGATTTAAGTTAAGTTAGTGGATTGAACCTAAAACAAT
TAAGTCCCCCCATCCCCCCCCCTCCCAAAAAAAAAACCAAGAATTTTGTTGCAGCTCATTTTAAATAAGTATTTATTTTTACAATTCAGTAAATCCTAGT
GAGACAGTGTTTGTGTAGGAAATTCAACCAAAAGAAACAGACATTTCTTAAAATGTCTCTGGGAGTGTTTAATAATTGTTTAATAGGCGATCAGAATTTG
ATTAAGGCAATTTAAATGTGTATATACATGTATATGCTGGGTTTCAACCAGAATTTGGGTAAATTATGAACAAACCCTACCACTGGGTTAAACATTTACA
CACACTACATTTATCCTTCTTTAAAATAAATGATACATAAAAATACTAAAGAAGGAAAAATTAAACAACACACAAAACAGCTTAGCTAATTATGCATACT
ATCATTTTCTAATCTATTTTGGGTCAGGCCATAATCATTTATAAGCATATCAACACAATTATAATTATAAGTAATTATCTATGTTCTATCATGATGTTTC
ATGATTTTTATACAGAATTTTAGAGAGATTTTAACTTCGTGTTTATGATGTTTGCTGCTTTGGTCTGCAAAACAGATTTAATTTTAAAGATTTTTTTTAC
ATGTAGACTACAGAATAATAAATTATTAATGTGTAAATATTTTAGGTTCTCCAGTCTTATTTTTCTCTTGCATAAATTAAATTGCACCAAAAGAAATGGA
ACATAAAAGGACGAAATTATACAGTTATTTTTAATTTACAGATTTTGTTAAATTGCCTGAAAAAAAGTTTAACAGTCTCTTATATAAATTGTCTTCTGTA
CTTTTGCTTCTTTATCCTACTTTTTCTTTCTATAGTTTGTGGCTGATTTAGAGGTAAATGCTAATTATAATATCTAGGCCTTTTTTTTGGTGTAGGAAAA
ATGCAAATCGTTGTAGACTTGCTACTTTCTGGATGCATCAATCAGTTCTTAAATCGCTCCATAAAACTCCCGGAGAGAGGTAGAGAGAAAACACCACAGT
GACGAATTGATTCGCCCCAATTTCTCCAAGGCCGTAAAGTCGAGAAGTTGAAAGCAGAGGTTATCAGTACTGCAGGCCTAGAGCTTGAACAGGTCTACAT
TTGACAGCAAAAAACATTTAAACTTTATCGTCTTTTGCCAACTGATAGCCTATGCGCTTATAGGTCTATAAGGTGATCTTCAGTTCGGCTATATGGTTTC
TCTCATTCATATTTTGAATGCAACACTTCAGCAGTCCTGTCTAAAATGTTGGCTAAATTATTTAACAAAATAACTAGGCAACATAAATAAACATATTAAT
GAAACACAGAGGCCTAAATGCATTCGCACGATGCTTAAAATCGTGTTGCTGAAGCCTAGTTGTACTATTTAGTCCGACGGAAAACACGCTGTTATTATTT
AGGCTAGGACCCTTAAATTAATTTCGTTTTAATTAGTATTTCGGTTGACGAAAAAATACCTTAAAAGTGAGCTTTTTTAAAAATTCAAATCAACATAGTT
TTAGTCAAATAAACAAGACTATTTGGTTTGATTCATGGTTGCATGATCACTTGTTTACAGCAATTTCGATAAAGTTTTAAATAGAGTAATACGATAGTAA
CTTTAGTACGATAGTAAATTATAGTTTTTTTAAGTAATATATCTTTTAAAACGTAGTTGAACTATTTTTTCGAATCACATTCAGAAAGATAGCTGATGCA
TCCATTGCATTTTAGAGCATTCAAATGTTTTTACTATTATGCATTTTGCAGATGTTTTATTGTTTTCTATACTGAGAAGTAAATATTAAATAAATAGAAA
TTGGCCTACTGAAATTTAACAGACTCTCTCGACTTCCGTACATGCAGTAGACTAGTAGGCCTACTAGAAGTCTGCAAAAGTAATATATACTTTTAATTTT
ATTTAATAGATAAAGATATTTTATTTTAACAAGCTTGGATATCGAAGAATATATTTGCTACATTTTACGTTGGCTTATTTTTAAACGGCCTATTTTTAAA
CGTGTATACCGTTGGAAAAGTCTATAACATTTTGAAACACTTCCATTAATTAGGTAAATTGGTAAACAAAACGCATATAGGGTAAACGAAGAAGAATGTG
TTTTCATTCTCACTGTCATTCAGTGTTGGAAATTGTGTCCCTACAGAATAGTGCATGTCAAGAAAATGACCCTCATATTGGTGCCTGTCATTGTCATGCA
TCTGCTAATGTGATAATCACATCTATTACTGCATGTGCTATCAACAATAGATAACAATCAATGTATATTAGCATGTACAAATTATTTGTAAATATATTTC
CTTTTTTAGGAGATCCTAAATTGCCTGCGTAATTTAAGAAGTGGCCACACTTTTCCATTTTTCCCCTGTGTGACTTTTGAAGTTTAAATCTTCCTTTGCT
ATGAAAAAATATATAATTTGGAAAACTCATTGACTATAATCCAGATTCTCCTTTTTTGTTTTTGTTTAAAGCTCGTATTTTGTTCTCCAGCACTTACTCT
CCTGCTGTGTAGTGTCGCTCCCGCTGGTGAGTCCCGGAGACTCCAGCATACTCCTGCCCGCCTCCCCTGCGCTTTCCTCCGCTTGAGCTCCCACCATCTC
CCGCGCCTCCTCCAGATCCAACAGGTGGCTCACGGAGAAGTTCTTTTTAGCCTGTAAATGATCCTGATTCGCGGTGATCGCAGTCTCCAGTCGGTTTGGT
AGTGTGGCTTGTCTGTCCATTACATGAGCATAGCTGGAAGTCATGACGCAAATTGACGAAACCCAAGATATCCTCTACCTACAACCAACATGGGTTACGT
TTCGACCATAAACAACCACTCTATACTCGACAAATTGATCGATTTCTGTTTTTACATCCGTAGATCGGCGTCTTGTGAATCCAAAAAAGGCGTATTTGCT
AGCTCTCTTTTATTTGACAATCGCGCGTCTTCATGGGAAACTGCATCACAGAACAAACATCTCCAGTTCAAGAGCTCCAAAGTTTCCAAAGTTTTGCCGA
TGACTCCACTCTTAACTCTGCAGCAGAGACTTTCCCCTCTCATTCCCAGAGTGAGGGAATGAGTAAACTGGCGAGGAAGGTTTGATGGCACAGATGAAGG
ATTTACGCCCAATGGTAGGAGGCCGGACGAGCTTCTCTCAGCTTCTTTCTCCGGTACTTCAAACGGCGCACACTGGCGCTGAGGGGGGAGCCTGAGAGCC
AAATACACGAAAATTTGAAGCCCTGGGGAAGGATGATACACATACAAGAAGTCCTAATTACGTGGCGATGTTTTTGTCTGTGTTTAAACGTTTAACGTCT
ATTTTTTCCACCTCCACTGAAATACGGAGTCTAATTAATACGTAATGACGTGATATAAATAATTACAGAATTTTAGAAATGCATTGAAAATGCGTTTCAT
GCATAAACTGTCTTAAATACAACTGCAAATATGATTAACGTATTTTAGAATTTGTTTAGCAGGCTATATAGTCTGTGTAGAATTATTGTTTTATAGGCAT
TATTGTTGTCATCTTTCACCCAACAAGTGAAATAAACATTTTAAGAAGGCAATTTTCGGAAAATGATCATAAAATATTTGAGAATTTTACTATAATTTTT
TCACAGCTGTTTTCTTGTTTAAAAATGAATTTTAAAAATATAGAATTTTTTTTATTATATGTAGGCCTAATTTTGTTGTTGTTTCTTTGAATATAATCAG
TTTTAGAGGTCGAATTAACATTCAAAATTCTTCCACTATACTCACGCTGTGTTTTGTTGTAACATGAGAATAAAACCAGTAGGCCTACCAAGTAATAAAT
GGGTGAATGACGTAATCCATTTTGAACGTGATAAACATTGGATCATTTTAAGTAGGCTAACTGATTTCCTATGCAATATAAGCTTATTTTATTTATCAAT
TTAATAGCTAGTGCATACTTAAATGGCTTAATCGGAATTCTTGAATTCATTCATGCTATTATTATTATTATTAGTATTATTATTATTATTATTTTGACTC
ATTTTAGTCTAGCAGTTAATCTGGAACATTATTTGTCTGTAAATCTAGTTACGTTCAGCAGTTGGCCAAGCATTTTTTGTTTTAAAAAGATGTCTAATAG
ATGTCTAAACAAAGTCGTCTTGGCTAAAACAAGACTAAATTTGGTTTGTCAGTGAAAATCTATCTAAACGTAATAATAGACCAAAAGACTAACAGAAATG
AATGACTACACATAAAAAGTCTGTTTAATCTGTCTAATTGACCACTATTTCGGCTATTCCAATTAGGTTTTCACTGACACAAGTTTAGCCTTGTTTTAGC
CAGGCTGTGTACGTTTAGATGTCCATTAGACGTCTATTAAACACAAAATTGTTTGCTGGGTGGGATTAACGGAACATGCATTTTGTGTTTCCAAGCAGCA
TTCTTCAACTTCACTAAATCATGGTGTGGTGGAGAAAAGTTTGAATCAAGAAATAACGGCCTTGTTAAGCTCAAGTGATCTTAACCCAGCCGGCACACGA
CCGACCTTCAACGTTGAAATATGGTTGAAATAAGGTCGGATGTGTTTTGAACGTTGAAATAATGTTGATAGAACGTTTAATGTTGAACGGTTGAAAAACA
GCCTGCAGATGACAGGGCTTCAACGTTGAAATGTGGTTGAAAATACGTCAGTAGTTGCTTTAATGTTGAAACAATGTTGAATGATAAATGGTTGAAAAAA
GTTACAGAAATCCTTGTACAAATCAACGTTGACATTTTTTATCATGATCTGTATGTTGAATTAACGTCATGGCTTCACCCAATATTCAACATGTTTGGTT
ACCATCGGAGGACTCGGCGCACGCGGCCACGCGCAGCGCTTCACTTCACTTAATTACGAACACCTGACCTGCAGTGCAGCTGAATTCAGAGGTAAGCGTG
TTTTTAGATGCTTTAATGTTTTATTTACCCCCGATTTTGTTTTACACACAGTGTATCTTGAATGTGACTTGGTGTAGTGTTTCGGTCCGTGTATCATCCG
TTCATTTGATTATTCCTTTTATTACGGAAAACGAAACATCAGAAAAACGATGAATATTTCGTTTTTCTGTTTATTCTGTAGGCAGAAAAAACGGAAACTC
ATCTGTTATTCTGCTTATTAACTTAAAACGAAATATTAAAACAAACACAAAACCAAAACGAAATAATGAGTCAAAAAATGGTCCGTGTACGGTCCGTGTA
CATTTTGTTCATTTGTTTTCCATTTTCAAACGGAATAAAGGAAATGGAGAAAACGGCCGTTTTTCCGTTTTTCTTTTGCTCACGAGAAAACGAAAAAAAC
GAGATTTTGGCTGGATTTTCGTTTTTTGGTTTAGGGTAGGAAAACGGATAAACGACTTTAAAATTCATTTTACACATGTAGGCGGTACTGAAACGCCCAC
TTTCCTCTAATTGGTCAACCAAATCTGCGCTCGTGACGTCACCCTCAAAATAAAAGCCTTACACGCAGGCTTTTAATTATTATATTTAATTATATATATA
AACAAAGTTTACATATTTCTATCATTTGAACCACACACAGTTAGCAGGCTTACATTCATTGCGTATGTATAAATTAAATAAAAACGTAAAGCAGTATTCT
TAGACATTTGTCATTAAAATAAGGTCATAGTTCACTGCCAAGCTTCTTGCTGCTGTGCACCTGATGCAGAGCAAAGATAGCCATTTCCGAGGAGCACCTG
GTTTGCATGATTGTTTCAGAACAAGGGCTTTATAATAATAATAATAATAATAATAATAATAATAATAATAATATAATAATAATAATAATAATAATAA
TACTTATTATTATTATTATCATTATTATTATCAGTTACTAGTGTTTTTATTGTTTGGTGTGTTTTTTTTTCTTTCTTTTTAATAGTTGTTAGCAGAACCA
GAATAAGAAACTTTGGGCCCTGCCATAAAAACAAAGGGTACCATTTGTTTTCATTTATTTCTTCCTATTTAATTATATATTGTAATTATTTATTTGATAA
TTAATACTATTTTATTGCAACGCAAAGTAGGCCTGTCTTATTATTTTTTTATTATTAAATTTGCTGTTATTGTAGGCTAGATGGCAGGCTTTAAATATTT
TAAATAACATTTACTCATAACCTTACTAATTCGAGAGCCGAACCTAAAAAAGTTTAAGTGCAATAAGCCTTTACAATAAAGGGAGCCGCAATGCATATTT
CTAGCGCATTCTTTCACTCAACAATTATTGCTGGAATAATGAAATAAATTTCAAATTAACTAATTAAACACATTATAGTTGATAAAAAAAAATTGTGTCT
AGTAGTTCCGACAAACAGATGAGCTCGCTACATGTGCGTTGCACTAATAATTTGACATTTGCTCTTCTTGAGAAAGTGCTACAGAGGGCCCTGCTATTTC
ACATATAAGGTTAGTGATAAGTATTTTAAATTAGTATTGCACATTATTCATTCATTTTCTTGTCGGCTTAGTCCCTTTATTAATCTGGGGTCGCCACAGC
GGAATGAACCGCCACTTATTTAGCAAGTTTTACGCAGCGGATGCCCTTCCAGCCACAACCCATCTCTGGAAACATCCACACATACACACACACTTATACA
GTATGGACAATTTAGCCTACCCAATTCACCTGTACCGCATGTCTTTGGACTGTGGGAAAACCGGAGCACCCGGAGGAAACCCACGCGAACGCACTACACT
ACTGAATTAGACACACAATCTCAAAAAAAAAACCTTGATGCTTATTTACATTATCTAAAGTCATTTCTTACGAGGTAGGATCTGTTGTTGTTTTATATAT
TTTCGTTTTAATATGCAAATAATTTTCGTTAATTTGTAATAGGCCTATTGTCCCTCATTGCATAAAATAACCAGCCTTTGTATTCAATATGTTTGTAAAT
TGTCCATCCAGTTTGATGCAAGCTAAACAACATTTGGCCTGTTTTGCTTATTAAATGAGTGATTTTCACGGAGAAACAAATCATCCATCCACACAGTTTT
GTATAGACCTGTGTGAACAAAATATACACTTCATCATAGCCTATAGGAAAATGTATCTGTTTAGAACCAATTTGTGTTGTATAATTAAAAATATTAAAAA
```

CAAAAAAAAAAAAAAAAAAGGAATAACATTTGAAGTGCAATGCTTCAAAGATCCAATACGCTGCATGCAGAGCCAAAACAAAACTGCTTTGTTTGTTTAAT
AAAATAATAATATGAAATTAAAGTGAAATAGTTCCATAACCTGTATTATAACTGTAGTCACTCTAGATGACAAATCATCCGTTGCGCGGTCAGTTTCACC
TTTAATCAAGAATGTGCCAGAACCAAAATGAGAATATTTACAGATTACCTCGGAATGAGACCTGGTCTGTGAAGGATGCAGACACAGAAACTCGTTGAGA
ATGGAGATGCATGTTAGATGTTTTTATCCCTTTAGATATTGTAAATAAACAACAAGTTTTTATAGAGATGTGTCATGTCTTTTTTTTAACGCACCGCTGA
ACAGCCCGAAAAGGCCGCCTTTATTTAGCCTGTAGCCTAAATAATAGTTTAGATTTTATAACATATATGTTCCATTACTTTAAATGTATAATGTTTTTTA
ATATTTATGGATTATATAAAATAGCTAGTTTATATAAATGCGCTCTGATCATTCAAGGATTATGCGTTGCATATTCTTCTTATTCTTCATTATGGCTTGA
TCCGTAAATTGTGTATTTTATAAACTGATCGTTTGTAAGGACTAAGCTATTAATAACTGTAATTCCTCGAATGGAAAAGTCTTGATTAAAGAAAAAGATA
ACTTAATTCGACATATGCGTATAGTGCGCTTAATTCTGCGCGCATCGCGCGCTTACCGAATTATCTTTTTATTTTTTATGCACTAAAATGCATCTGGCAG
CACCGAAAAGAGCAAATGCCAAGTTATTAGCGCAGCGCACAGACAGCTAGCTCGTCTAGTCTGAACCAGGGGTTTTCAAACTATTCATTGCACACCGATC
CCATTGCATTTTATTTAGGATAAACCACTTATTTAAAATTAATTAATTAAAAGGTGCGTACTCTCCAATACACCACTGAATTATAGTGTCATAATATTTA
ATAATTATAAATAAGATAGACACCTGACCTCCATTTAATAAACAATTTATTTGCCATACTCTGTTAATTTGAAGTTTAAAGTTTAAATCATTTATGTTAA
ACTTAAATTATTTAAAAACTTAAAAAGACTTAAATTTAGTCATTTAAGACATTTTAGGTCATAAGTTTACATACATTGACCGTTGCATTTAGTAGGCCTA
TTCTGGTCTAAATAAGCTTTTTGCATCTTAGGTGTTTACACACACAAATTAAGCAGTTTATACGTACATTTAATGTTACTTTGAATTACTTTTGATGCAA
TAAATAATCCTAAATAGTTGAAAGTGTTAACATATTCTTGAAATGTAGTTATTTTATAACTATGAAATTTATTACTAAACAAAGTGAATCGTTATGAATT
TGCCAAGAAATAATCCACTTTACCACTGTAAAGCAATGCCCCACACTCACTAGAACGGTAGAACCTGGACTGTATGATATGTTATTTATGTAAAATAGAG
AAAATAAGCAAGATGCGCAGTCTCATGCAGCCTTAGTGAAATATAGCGCGTGACAGCAATGAACTAAAAATAAAACCACCCTTGAATGCTGGCGATCTTA
ATTTTTTCAGCTTACACTGACTGAGAATAATGTTGCACTATTTTTATTAACAAATAATTAGAATGTTTTTTTTTCAACACATTCATCATTACTTTTGCC
TGTGCAAGTTTCTTGCGCGCGCTTGAGCGCTAAGGCATCTATTTTCAGGTGCAAGATGCCACTTTGGGTGGCACCAGCACACTCTGCCACACCTGTGGCT
ATGTCCCTGCTACAATAATAATAACAACAACAACGACAAAATATTAACAAAAATAATAATAATAAGACAGGCCTACTTTGACGTTGCAATAAAAAAGTAT
TAATTATCAAATAAATATTTACAATAGGCAAAAGGCATATATTAAATTAAAATAGGAGAAAATCCAATAGAAACAAATGGTACCATTTATTATAGCCCCA
TGCAGGGTCCAATGTGTTCTTACAACCATTGAATAACTAAGTAAATAATAATAACAATACTAATAATAATAGGCCTAATAATAATAATAATAATAATAAT
AATAATAATAATAAATTATTATTTTATACAGAGTAGGCTATTATTTAGGCCTACAATAGCTCTGAAACAATCATGTAAACCAGGTGCTCCTCGGAAATGG
CTTTCTTTGCTCTGCATCAGGTGCACAGCAGCAAGAAGCTTGGCAGTGAACTATGACCTTATTTTAATGACAAATGTCTAAGAATACTGCTTTACGTTTT
TATTTAATTTATACATACGCAATGAATGTAAGCCTGCTAACTGTGTGTGGTTTCAAATGATAGAATATGTAAACTTTGTTTATATATATATAATTAAATAAT
AATAATTAAAAGCCTGCGTGTAAGGCTTTTATTTGAGGGTGACGTCACGAGCGCAGATTTGGTTGACCAATTAGAGGAAAGTGGGCGTTTCAGTACCGCC
TACATGTGTAAAATGAATTTTAAAGTCGTTTATCCGTTTTCCTACCCTAAACCAAAAAACGAAAATCCAGCCAAAATCTCGTTTTTTCGTTTTTCTCGTG
AGCAAAAGAAAAACGGAAAAACGGCCGTTTCTCCATTTCCTTTATTCCGTTTGAAAATGGAAAACAAATGAACAAATGTACACGGACCGTACACGGACCA
TTTTTTGACTCATTATTTCGTTTTGGTTTTGTGTTTGTTTTAATATTTCGTTTTAAGTTATAAGCAGAATAACAGATGAGTTTCCGTTTTTTCTGCCTAC
AGAATAAACAGAAAAACGAAATATTCATCGTTTTTCTGATGTTTCGTTTTCCGTAATAAAAGGAATAATCAAATGAACGGATGATACACGGACCGCCAGC
ACAGGCTGTTTTTTCAGCTGGGTTTATGGTGTAAAAAGGATTTTTTACACTAAGAAAACTCTTTTCTCGAACTTTTCTCTGAAAGGTACATCGGGAAAGC
AAAAGTGGTTAATTTCTGTCTTACATTATTGCAAAAACTTCGATTTTGAAGCACTGACAAGATTATGAGAAGAATAGTTTATTTGAAAACAGATGCTGCC
ACCTGCTGGCGTGTAAACACATCACATGATTAAAAAAAGGTATAATTTTAAACAACCCATAACAAAATGGAAACTTTTTAAAGATGCAATTTTTTATGCG
TTTACAGTTTTGTGAGATTTCCTTCCTTTTTCTTCTTCTTTTTCAGATCTGATTATGTCTATCTGTCTATTTATATCTGATTATAGATACTACAAAAAGA
TCTTAGTCTTGAGTGCAAATTTAACTGATAAAAATCAGTTTATATATATATATATATATATATATATATATACGTATATATATATATATATATATATATA
TATATATATATATATATATTATATATATATATATATATATATATATATACACACACACACACACATATATATATACATATATATATTATATATATATATATATA
TATATATATATATATATATATATATATATTATATATATATATATATATATATATATATATATATATATATATAATAAAGTAAATTCAAGCAGCAGCTGTATA
ACTGCTATGGATGCTGTTTCTGTGGATTCAGCAAAGTGCCACCGCCGAGACACTTATCAGCCACTCTCACATGACTGGTTGTGCTGTCAGCATTGACCCA
GCAGTCATTTCAGCGTATGTTACTGCGCACGCCAATGAAACACAGCTAGTCACTGTCTAGGCCTAGTTCTAACACTTTTTTTGAAGAATTACAGTGATCT
TCAAAATCCTGTGCAAAAAGAAAAAAAAAAGTCAAGTTTTAGTGATGGTCGTGCAAAAGAAACAGGAGCGACATGGTTACATTCAGTCCTGAATGGAGTG
ATTGTAGAGAACACGTTTCCTTTGAGGGATGTCAGGAGGGGTGGGGCAGAATGTGGGCTGGTGTTAAGGAATGCCACAATGTGCTTAGCCCTTTGTTCTC
ATCACTTTGAGCATTCTTCTATGACTCAGCCACAGCAAACAAAGACTTCCCTCTCAGCTTTCGCAGGTACAACGAAGAAAACGCTTTAACAAAAGAGGAA
GCAGTCTACAGCTCTGTGGAAGGGCAGAAATTTTCCCGACGCCTCGGAGGGAGGAGTCTAGTAACACTCCTTATTTGGTTGTAAGGATCACGTGGGCAAC
CGCGTCTGACTCTGAACAACAGTTGTGAAAAAGTCAGAGAACTTCGCACTCTGGTGAGTTCAGGAGTTTTGGGGTGAGACTGAAAGAAAAAAAGACCCAA
GGAATAAAACACAGAGCACATGGTGTGAGGGGAAAGATGCCTACTAACTTCACAGTGGTGCCGGTGGAGGACACCCCATTATCAGACCTAGACTGTGATG
GAGATGGACTAAAAGAAGAGGACAAGGAGGAGCCAGAAACGGCAGCAGGGCACTATCCAGGTGAGAGCTTTCTGAAAAAAGTCATGTCTGGACTTGTATA
AACAGATGAGCTGGGCTTTTAAATGCTCGTGTTTTCTTTAAAGTAAAGCAAAGTAACTGTACTGTAGGTCCATCCTCATAAGCAAACGAGTGCCATGTTT
ACTGCCATGTGTTTGTGACATTATCCTCATGGAGTGTAAACAAGTGGCAGTTTCCTGACGCCCCTTTTCCATTTTCCTGTTTCTCTCCTACTCTGTGACG
TCTTTCGCTGTTTATTTTTAGATAAATTGCATACTGTTACACCACAAAGTAGAGCTTAAAGCCAAGGTTGCAGTTGCACTAACCTGAAGTGTACTTCTAT
TGTACTTGAATTGTAATCAATTCCTCAGTTCTGGATCTTGCTTCTTTTGTGTGTTCCTCACAGAGTTTAAGGGATAGTCGTCCATAAAACAAAACTGGTA
CAAAAAATTAAACGTGGACTCTGAATGCAATGCAAAATACTTTGGATTAATGCTTGTCAAATGTGCAATGTAATGTCAATGGAAAAAAGAGAAATGTCTG
CCATTATTTACTCACCCCAGTATTGTTCCAACTAAACATACAGGGTTTTATTCTGTAGAGCACAAAATAAACAAGTTGATAATTACATGTTTTAGTACAA
AGTTAGTATTAATTTAGATTTGATCAAAAGTGGACAATGCATAATACAGGTATAAACAACATTAGCAACAAGGAAATCATCATTAGAGCATTTTAATCAG
ACAGTGTAGTACAACAGGTCCTTTAAGGGCACCTATCTTTTTGTATGCTGTTTGGACAGAACTGTTTGTAGGTATAATGTGTCCACAGTCATGTTTGAGT
GATATAAAGACAATAAGCCTCTTTTATTTTTATTTTTTAATAGAATCCAAATCCCTCCCATTTTGAGGCCCACCGCAACGTGATGTAGGAGTGCGGTTTT
CCCCGCTCACCAATTTGATTGACAGCCGTGTATTAATGTGTCTCTGTATTAATGCGTATAATCATATCAACTAGACAGGATGTGCGCAAAGTATTTGGGA
TTTAAAGATCTGTTCAGCTCGCTGGGATCATCAATCATCATCAAATGTAATCAAGAATGACTTTTACATGTTTAAAACATTTTTAAAACAGGGCATGTTT
GTAATGAATTACAGCGATTTTACAGACTTTACTTCATCACCACAGCTACATGTCAGAACAAATTTAAAAGAAGATGCTTCAATCCCGGTTTGTGGACATT
AAATCAGGTTTATTTTGTACATTTAACATAACGGATACAGAAGTGTGTATTAACGCGTATCCTGTCACATTTGCTGTGCAAAAACAGTGCAAAGTTAAAT
GCGCATCATGTGTGTCTTTGTGTGCGTGTACTTTGTAATGACATTGTGTGTGACTCATCATTGCAGAAAGGCTTGAATTAACTCAGCAACAAATACATTA
AATAATCATTGCGAAAGTTCTTACTGTGGTATTTCTTACAAACGTCACGAGAGATCTGCTTCCTTCATGTCTGTCCCTGTGCTGTTTATCTGACACAGCC
GAGGCAGACTCTGACAGACACGTGGGGACTGTGGGCGGGGAGAACTAGCATTAAAGGCAAAAAAACAGCTACATTGTGTTCAGAGCAGAAAATCCAATAT
TCTGGAAGGTTTAATAAATAATCTAGTGGGTATTTTGAGCTGAATCTTTACAGACACATTCTGGAGACACAAAAGACTTATATAAAATCTTGAAAAAGGG
GTAAAATAGGTGCCCTTTAAAATACAGTATTATTTACAATAAAAAATGAGTTAAAAAGCAACTAATCTTTAAATGGTCTGTTTTGGGGATTTGCCTTTTA
GAATTTAGTTAGAAATGCTTTTGACCTGACATTTTCAAACCACAGAGTGTTTGCAAAAAATGACAAAAGACAAGCCACTGACTTAATTCTTGAATAATTT
GAGAATTATTATGTTGAAGTGTGGCACTGTGATGTTTAACTTGAAAAAGCACAATGATTTTTAACAGAAGCCCACATTGTTTGATGTAGTTTTGTAGTTG
TCAGAAATAAAAGCTGGTGTTTCCTGACATTTACCGCTAATTATTTGCATGTTCATATGCAAGTTGCACAAACCTACTTTACAATATTACATTAAATGAC
AAAAAAAGCCCATACACTTATCAGCCTATGGACCTTCCTCTCAAAGAAATCAGGAAAATGGACACAAGAGTGTCTCAACAATGTGTGTCTTTCATTTACT
TGTAATGTGATTGTCTAAAACAAATTGGTCACCACATGTAAATAGGGTCTTTAGGTGATTGAAAAAAAGCTAAGGTACTTTAGAAATGATATTTTTATGTT
CACAAACAAAACAAAACCAAAAAGTCTTTTTTTATGTAAAAGTATATTTTATGTAACTTCATGAAAACAGGAAATTGTGGTCTGGTACCAGCAGTTATTC
AATGTTCTTTCATTTTTTTACAGCACACATATTTTATTATCCTAAGGTTTGTAGTTATCATTCTCGTAAGTATTGTTTAATGCAGATGTTTCTTTTATGT
GTATTAATGTGTTCGTATGGGTGTGTGGCAGCGCTCATGGGTCATCATTATGGAATTTGCCCTCTTCCTTTAAAATGACCTATTTATAACTAACCGTGAC
AAAAATAAGTTTAATTGTGCTGTTTCCATGTGGAAATGAGAATGTGCTAGTAAAATACCTCGGTATGGCTGTAAAGACTGCATAATTGGTAGATAATTTT
ATCCTCATTGATCAAATAAAGACATTTACCTGTCATGTACATATTACTTTAGAAAGTAATGCTTTAAATATTGAGTTACTGCCCAAAAAAGTAAATAGCT
GCGTTACTTAGTTTCTTTTTATTGTAAGTAATGTTACGTTACTTTTGCATTACTTTTTCAGACCTGGCTGAGGCTTGATCTCTTTCAAAACTTGCAGCAC
TGTATTACCTTCATTTACCTTTAAAAAAATAATGTAGGATAATGTTATCTACTGAAAACTTCCTGACCCCAGAGCTGTACATGCCAAATGTACAGAATAC
AGATTAATGAAAGTTTAAAGCAATCTTTTTTGCTGCTTTTGTACTTGTTTGTCTTGTTAAAAAGTAAAATCACTTTCCATTCAGTTAATCCTCTTTTTCT
TTCCTTCCATGTTTTTAAAACAATGAGAATATTTCTATTGCAGAAACGCATTCTCTCATTGCATGGAGGATTCAACATAACTTTGTTAATTTTAAATCAG
CAATTTATTTTTATCTCGCTTTACATTTAAAAAAAGTAACTATAAATTTAAACTAAATAATATTACTTAATTTTTAAAAGTAATCTTACCTTTCTCGTTA
CTTAGAAAAGTAGTATTATTACATGTTTTACCCCAACACTGCATATTACAAATACATTATTTTTTCTACAAAGTTCAAATACTGCCGAAAAACCTGTTCT
GTAAAATTTACCACATGCTTTCTGACATCTGATTGACAGTTTATTTTATGTTTTAGTAAGTAAAAAGTCAACTGTACAATGTTCGCCTTCGGGTGTTTTT
TTGCAGTAAAATTGTAAGGATACACAAATAAAAAAAAACTGTTATTAAAATTTTAACAAACATTTTCTTACTATTATATAATCATATTAAAGTATAGGTA
ATACATGTGATGATGCTTTACAGATTTGTACTTCTGTCAATTTATATTGGTTTTATGAGATTTAAAAAATATTCAAAGCTTGGTGAAGTGATCGTAACTG
AAAAGGACATTCTAGTTTTCTAGCTGTGGCTATAAAGCCTCTGTAAACATCCATGTAAGGCACTTCGAAATGACTTTGGGTTTTCAAAACACCATGAAAT
GAAATGATTATGGTTACAGTACCTCAAAAAGACATGGCTAAACATAAAAAGTAAAATAATGTATTGCCATGGCAACGATTTCCACATTGAGAACATCTAC
TAAGATAAATGTATCTGTTAACTTTCTTTAGTTTTTAGTCATGATAAGTGTTAAATGTTATCTTGTGACATCATCAGTGACTGTACCTAATCTAACAGAA
ATTTCATACCAACAGTACCCTGCAGTTGGGGGTAAAACAGACCTCTTGGACAATATTTTTCTCATTATTTATTTTTAATTTTCTGATATGTTTTCAAACC
TTCTTGTTTTTTTTAAGCATAGCGGAGTCCTCTAATTGTGCAAATGTGCTAAATGATAAAAAAAAAAAAATATAAGTGAACAAACAAACAGACACAATCAA
ACAAGCCTTTGACTTTGCTGACCATAAGAAACCTCTTTCAAAAAATATTAGGAAATCTTATACTTATTTTTTATACAAACTTTTAGAACAATATGTATTA
TTGTCCAGACTGAGTATAAATCAGGTTTAGAGAAATATAAGCCAACTCTGAGCTTTTCCAAAACATTACAAATGAGTTCCTTTGGGATTTTTTTATTTTA
TACACATAATTAGCTGTGTGCAGGATTATAAACACAATTTACAAGCTACACGGAAAGAATTCTCAGGAAAGGGATGTATCTAAAGCCTTGAATGCCTTTT

TTTGTGAACCTCTTCTTGTATTGTCTTGCTATCACTGTGGGAAGAAACATGAAACTAAATCTAACAGCTGCACATGCTTAGAAAATAGATATTTCATATT
GTATTTAAAAAATAAACTAATTTAAATAACTTAAAAAAAATTTACCATGAATGAAATGAAATTTTATTAAAGATTGAAATTTTCAACCTATTATATCAAG
GATGGGCTTGGATCACAAACTTAAGATTGGTCCCTTTTTAAAGATGCTTGGATTTTTTTTTAAAGACCTCGAATTATAGCTGAACTAAAAGGTTTTAAGT
TAAACTATGGACAAAAAAAAATGTTAATTTATGGCTAAACTAAACTTTAATTAACTAACTAATGAACTAAAGTTTAATATAAGAAATGCACACACACAAC
TTTTTTCAAGTCTGATTTGAAATGTATTTTTCAAACACGCTTAACTTTTAATGTGGGAAAATAAATAATTTAAAACATTTAAGGTTGATATATCAAACAA
ATGAGCCTTCAGAATTATTTTGTTTTAGACGGTACAAAATGTTTCAGCTTCCAAGTTTTTAAAATAAATATGTATTGTTTTTGGCATATCATCATAGGTG
TTAGAGGCAGAACACTGCCATCTGCTAGTTGATGAAAAAAATGTAGAGGTCTCAGTTTTAACAAAATTTGAGCGTATTAATCCAGGAGAGGGCCATTGAA
GTCCGTTCATTTGTCAAGATGTAGGCAGTTGCTCCTATCACAATTTATATGCAGTTTAATTTAGAATTATTTTTATTTTCTTTTGTAAATATTTTTCAAA
TGATGTTTAACAGAGCAAGTAGGTTTTTACAGTATGTCTGATAATTGTTTTTCTTCTAGAGAAAGTATTATTTGTTTTATTTTCAACTAGAATAAAAGCA
GCTTTACATTTTTAAACACCATTTTTTTGGACAAAATTATTAGCCCCTTTAAGCAATTTTTTTCAATAGTCTACAGAACAAATCTCCATATACAATAGCT
TGCCTAATTTACCCTAACCTGCCTAGTTAACCTAATTAACCTAGTTAAGCCTTTTAAATGTCACTTTAAGCTGTGAAGTGTCTTGAAAAATATCTAGTCA
AATATTATTTAATGTCATCATGGCAAAGATAAAGAAATGAGTTATTAGAAATGAGTTATTAATACTATTATGTTTAGAAATGTGTTTAAAAATTCGTCTC
TCCGTTAAACAGAAATTGGGCAAAAAATAAACAGGGGGATGAATAATTCAGGGGGCTAATAATTCTGACTTTATCTGTATATATTTATTTATTTTGTGAA
AATCTAAAAATGGCCAAAAATCTATAGACAAAACTTGAAATATGCTGTTGTACTCAACTGTTGTTATCTGCACGAGTTCAACATAATTAAATAAATAAAC
TGAAATACCAATAAATTATAAGACCTAAAGTCAGATTGTTGTCACACAATTTTCAATTAAATCAATAATAATAATAATAATAATTGCAGTAAGTCATAAT
TAACAGGAGTTTGTTTGTGCATGTTTATATGTGGGTGTGTATGCATGTTTTGTGTTTGTTAGGTTACACCCTTCTATGTCTGTATTTAAGACTTTATATA
AATATAAAGATTTTATATGCCAAATTACTGCACAAAAGCAATTTTATTGCCGCTTGTACCGTAACAACACATTAAATTGTTTTGATCCCTTTCGTCTTGA
CTGCAAATTACACAAGCCTTTTAGGAACAATTAACTTTGTCTAAATGTAATCAACTTAAATTTATGTGTGTGTTTGTGTGTGTTCACATGGCAAGGGCCA
CACCCTATACTGTATCATATAATTCTGATTAAGTAAAAAATGTATATAGCTTTCAGTCGAAAAAGACAGAACCGCAAATGGTTTAATTAACTTAGTGGCT
TATAGATTGTCACTCCCTATTATACTACTAAGTGTAAACATTCATTTTAACCATTATAATCAAATCACATCTTAAAATGTTTACCATTCTCTACTGTGTG
TTTGTTCTGGTTAATAGTCGTCTTCTATAAAAAATCAGTAAATCCACAGGTATGTTTATAAGGCACTATTTAACACACTTATTAACCATAAAAATGTACT
TGTTTAAAAATGTTAATAGTTTGCACTGTTGATGTTAATTAATTATATATAGGTTTATATATTATATTTCCTAAAGGACACACTGCTTTTTCTTCTTGTT
CTCTCAGCAAACCAACTGGTTTGTAATGTTAATCCCCCCTCCTCTCGCTTCCTTTAGTAGTCATAAAAGTCATGCTCAGTCAACATTCAGGTTAAAGACA
CAGAGGACGGACAGTGTAGTGTTATTTAGCTTAAGAGAAGACACCACCTGAGGAACAGAACTTTACATACAGTACAATTAAAAAAAAAAAAAAACTCTTT
GCTGAAGAAATGCTGGAGACAATCGCAGCCACCAGAGGAATAAGACTCGCACAGCTGCTGAAACACGGGTACAAGCCAATTCCCTTCAGGCGGACAAACA
TGGATTTTGACTAGGACCTGTCAGTCTTAAATCAAATTAGTT

#### --PacBio contig2--

**-Prrx1a side (reverse strand) – 5’ breakpoint ~ 2:23,174,000 (Zv11)**
-Middle piece of DNA2-8_DR – 3’ breakpoint inside of this transposon
**-Kcnh2a side (forward strand) – (transposon underlined – <u>DNA-6-N2_DR)</u>**

TATATATATATACATACATACACACACAACACAATATAATATAATATATCCACACTGAATCAAAGCACCTTAAGTCAAGTCAAGTTTGCTTACACTTAAA
TGCCTCTGCTATTTCTCAGTTTCAAGTGCAGCTCTGATCTTTCTGAATGGGAACATCAATCAATCAATTCGCATTAATGGTAGGTCTGAAGCTTGAGTAG
CATCCCAACACACTGACTGCAGTTCACACAACTATACACACAGTTCTGGTTTTAAATAAATCAGGTAGTGTGTTCACATTACAAATTCCAAAATGTGTGA
AATGACAAACATCTTTAAAGGAAGTCTTGTATGGCGTAATAAAACTTAAATGTAATCTCTTACTGAAATATTTAGTAGAACATACATAAAATGTTTACAT
TATTTAAAACAAAATCCTCGTCCTTTTATACATTTTGGACTAAATGGCACACGATTGTTTAGTAGGCGTGGGTTACATTCTGCATCGTTGATGCCAAATG
ATTGCATTCGGTGATCTATTGCTGTTACATATTTGATATTTGTACTGCAACACAAATCAAAAATTAAAATACAGACAACATGATTGTATTATTACGTTTG
AGGGCAACAAGAGAAGTGATGCCAGTCTTCCCAAAGAAGAGAAAAAGAGAATGATATATCCCAGCTGGCAACTTAAGGTGAAAACAAATATTGGCGTAAA
AAAAAAAGTCAAAAATTGATTACAGGACAGTGAATTTAATATAGTTTTGACATGCACGTTTACATGCATTGTAAGGTTAAAAATCCAGTGTACTTTAGGA
ATCTGGAACTGAAACCTTTCAGATTGTTCTTACATTATAATACTTTTTGTGTTGTTTTTTTTGGTGTTAAAAATGGGATGATTGGATGTTTTTATGGATG
TCAATGTTTTTATTAAATCGATTTTGCATTTTAACATCATGCATTACATCATCTCATTTTGACATCAAGGCAACAGTATCTGGCAGCAGACTTGCACATG
CAATCTGAGACTCGCAAAATGAGACACATGCACTCATTAGAGCTAATGTCCCTGTAAGGGGGTACAGTTAGGGACAGGGTTATTTAAAAGCATTGCATGT
AGTGCCAGATTGCATGTGCACCCATCAAGGTGCCCACTGTGATAAAAATGATCAGAAATCTAATGTTTAGTGGGATATATCAATGCTTTACCAATAAATA
ATTCAATAAATCAATCTCTCACCTGCACTATTTTAATAAACACTGTTATTGATTGTGGCGACACGATGGCTTAGTGGTTAGCACTGTTGCCTCACAGCAA
GAAGGCCACTGGTTCGAGTGCTGCCTGGGCCAGTTGGCATTTGTGTTGACTTTGCACGTTCCCCTGTATTGGTGTGGGTTTCCTCCGGGTGCTTCGGGTT
CCCCACAGGCCAAAGACATTCGCTATAGGTGAATTGAATCAACTAAATGGCTGTAGTATATGTGTGTGAATGACTGTGTATGGATGTTCCCAGTACTGGG
TTGCAGCTGGAAGGGCATCTGCTGCGTAAAACATATGCTGGATAAGCTGTCGGTTCATTTCGCTGTGGTGACCCCTGTTGAATAAAGGGACTAAGTCAAA
GGAAAAATAATGAATGTTATTGTTTGTTATTACAGACAGCAAATTTGAACAGTTTATAGGTTTCTGTAAATAAACAAAATAGTAATTGACTTAATAGAAA
ACACTTAGTTATTAACTAAAAATGTACAAAATTAAAAAATGCCTCTGAAAATCACTTTCAGGGACATAAATTGCCAAGTAATTCTCCAACCATGATCCAC
AACAACTGTCTCGCATCTCTGTCTAAGTTACAGGCTCTGCTCCTGAAACCATAAAAATCCAAAGCCAAAAGAGGAAAGCTTTGAAACCAAGCCTGCAGAT
AAAATCTGAATTTAATAGACAAATGAGGGAAAATTGGATTAGCGATCCAATCTGCGCAGTCATCTTACATCTGTTTTTGGGAACTGCACTGCCCGTATCA
CCGCCTGTGATCTGAGTCCTCCACGCTGCCATTGGCCTTCATCACTCAATAACTGGGCTAATGGCCTAACACTTCCCATTAATAAATGACTGCTCCTCTT
GGCCTCTCGCTCCTTCAGAGGTGATATTCATCATCGCTCTTAATTTTGTTTATCTTTTCATGCTCTCTCCCCTCTGCGATGGCCATCAATTAGCTACCTT
GGCCTGCCTTTTCGTGGCCTCCCTCAGCCCGTCGTCCCGCAGGGTCATAGGGTTAATATAAGAATGAGCCTAATTTCCAAACAGCTTCAAGTCCCGTGAG
CTATCTGGAATGTGAAGAGGTGATGCTTAAATGTTGCTGTCCAGGCCAAGACGATCAATATATCATGGACAAAGACAGTCGAGAGGATAATATAAAGTAA
ATGAGCATTGTTGCTGTGTTTCAGCTCTGCAGTTAGAGTTGTTATTGGCCCATTTGCTATACATTGCAGCACTGAAGTCAGAGAATAACAAAACATTAAT
AAGATCACTCTCTGTATTTGGGATCTGTTGTTTGTTGTTTTAAATTTGATACTGTTTTCATAAGTTTATTATTTTGGGGCATGCTTTTCAGTCAGTAGGA
AGGAAGAGGTTCTCAACTGATCACTTTTCACCAATTTATTTTATTTTATATATTTTTATTATTATTAATTTCAATGAACACTTTTTTTTTTTGCATTTTA
CTTTTTTTTACACTTTTTTGACACTCAAACAAATATTAATTTAATATACAAATTTGTTTAAGGTAAGTGGTTGCAAATCATTTATTTGGGCTTAATTTAA
ACAAACAAATTAAGTTGAACTTTGCTTTAATTTATTTGTTTAAATGTAGCCCATATAAATTTATTGCTTACCACTTACCTTAAAAAATAAATAGATGAGG
TGCTTGGTGTGGTCTGTAAAATGATTGAAAAATATGTTCCTTTGGAACCAAAACTTTGTTTATTGCATAAATATCCTGAAAATGTGTCAGTTACTGCAAA
CAAACAAACAAACAAACAACAAAAAAACGTATAGATATCAGCTTTTTACAAACTAAACGTGTTATAACTTTGGTATGGAAAAATACCCTAAGACCTGTTC
TTGCACAATGAATAAAGAAAATGCCTAATAATCTTGCTTTTGAAAATGAACATACATAATCACAGGTAAAGCAGAGACTTTGAAAAAGGGAGGTGTCCAT
TCAATTCTTTTCTAAATGAATATAGATAAAAATTGGATGTCTTCGGAAAGATGACTATAATTTAAAATTCAATGTAATAATGTTTTTTTTGTTTGTTTTT
TATTTGTATTTTTTAAAAATATTTTATAGTATTCTGTTTTATAATTTATTTTAAATGTAATTTTTGTCTAACAAATGTTGGGGGTGGAAAATACATTCT
GAAATATCATAAAAGTTGATATTATGTACTGTTTGCTGGGTTCTGTAATATATTTAAAATCAATAAAAAAACAAAGTTAATGTTTTTTTAGTAAATACAA
ATGAATATTTTTTTCCAGTGTACATGTTAATTTATGCTATTTTAATTTATCATTTTATATCAATTAACATGTTACAAGAGGTCCTTTATAAAAATACATT
TAAATGTGGAATTTGGATTTAAAGTTTTAAGATTTAAACTTACAAAATCACCTATTTGCTCTGCAAATCATACTTGAAATATGATTATTCATAAAACTCA
AAAACTTTTGAGGCAACTTCAAGACAGGTGCGTTACTTTGCTCTTGTAGCCCAACTCTTTTTAAAAGGGTTGATCATAATTATTTAATTATTTTATTTTA
ATTAAAAAAATTAAGCGTGAGATATTTGCTTGCTACAGCAGTATTTCATCTTCATAGAGAAGCACTCTACAAGATCTGCTTTTAATTTAGCCGCTTTCTC
TCCAGACAGCTTGGAAAAAAAAGTGTAACATGAGACGTAAAAAGACTGTGAGATAAATACAGCGCAGTAACGCCGCTGGCCACGCATTAGTTATAGATTG
TAGCTTTTGACACAGTATGTAAACGAAATTAGTTAAAATAATTGGATTACAATAAATCATTAATAACTACTGGACAAGTTCCATCGCGCGCTGAAAGGGA
AAGTTACATCTTTATTACGTGCATTTGGAGGAGACTTAAAGTCAAGACAGCTGGACAATGTAAACTGCGAGACATTTGTCAAATGTAATTGCGGATTAAA
AGTGCTTTGAATTTAGTTTTAGACATGTTTTAAATTAGAAAACCTAAAAAGTAAAAGTATTTTTTACATTTCTAAGAATATAGGCCTAGGGCAAAAATGT
TAATTAAATAACCTTATATTATTTTTATTAGGCTCCTAAATTTATACACTTGTTTTGTTTAATGTTATGCATTTCTCCAAATATAATGTGCATCTAAAAG
CATGTTATGAGTAACGGGACATAAGACCACCTTTTTTGTAATGCACACTAGGCCTATTAGATATTACAATTATTATAAACAGTTCTGGAACGTTTTACAT
CAGGATCCTTGTAAGATCATAATTATTATCCTGATAAGCGCCTATTGATTTGGATGAACGTGGATTGATCAATTTGACAAATAATCTGCGCAGAACTGCT
ACAAAACGGGTTTCATTTTATGATAGCCTATCGTGCTCTGTGTTGCTGACAAACCACATCACTGAGCGAGAAAAAAACTACAGAAAAATATAAAAATGCA
ATTTAGCTAAATAACATGAATTAATAGCCTAGACTATAACCTGATATACCAAATTATTTAGGCCAACCTAAATGTAGGCTATGGGCCATGGCCTGTGATT
GATCGAAAAGTCTAAACTGATTAAAGAATATTCATATAGATATATATGATAATCTATTATAATAGATTAGAATACAATAATATTCATATGTTTATATTGC
AATGTTTATTTTCCTGACTGTAACGTTCGTGAAATGTAGTTCTAAAATCCTGAGGTAAAAAAATGTAACAATAAGCAAATCAGTAGGCTATTTACGACTT
AGGCTACATTGTTATATGTGTCTGTTGTCATTTTTAAAATATCGTTAAATGAGATATCAATGAGTTTCAAAGTTTTCACTTCATTCAGCTTTTCAGCTGT
TGGCCTAGCGCCCTAGCCTATATTTTACAATTATGCAAGGAAAAAGTATGACGGATTGCCACAAACCTAATGGCAAGATACAAAATATAGCCTATTGAAG
TGAACAAAAATAGTCTGTTTTATAGTTGGATCTTCACTGATATTAGCATTACCACACGACCAATATTTCTTTTGCCTCCTAATAATGCACTGCAAGTACG
GCTCATTAAAAACAACCGCTGTGATATTTTATCATACGCCAACAAATAGGCTTCTCTTTAACTTTTCCAATGCTGATTATTTTACTTTATTCAGTCTATA
TTACAATTTTATTTTATTGCCTACATTTCGGTAAAATATAGAGCTTGCTCGACGTTGACGTTTTATTAACCTACTGTAATTTTTGTTTTAGGGCAGAAAT
GTATTGCTTTATCTGTATTCACAGATATAGCCTTAAGTCCAGCTCATAAAGAAAATCATGTAATGGAGGCTGATTAAAAACGTTTCTTGTAGCCTATACA
GGCCTAGTTTTGTTTCCTTGCGTTCAAAAAAGGCAAAAACCCTCATTCTTCACAACGACTCGAGGTTTCGATATGTGGCAGATTTATGGGCAGAGTATAA
TAATAATAATAAAAGCAAAGTCTCGCGCTGGCACCAGGCAGCGAGCTGGGCTTCGAGGGAGAAATGGGATCTTGGGGGAAACTAATGATGAAGAAAGATC
TCCTCAATCCAAGCGCCTTAATTAAATGCACGGAAAGAAGGAGAGAGGCTCTGCACATCTGGATTCAATCTGCGCCCCTTTGATGGCATCAAGTGTTCTT
TTCCCAGACCTCGGAGGCTGACAGAGGCACACAGGCAGCCTGGAGCCTGAACTCATGTCATTGCGACACATGCGGAGCTCGATCCCCTCCTGTCGTGAGG
CTCTTTAAAAGCGCACGCACGCAATGCTCTGTAAACCCACTGAAAAAGAACAACATTTTAGTGTCACATCACGAGCTGACACCAGTGCTGAACGCATGAT
ACTCATGAGAGCTACGCCAGCGTTTATGACTTTGATCATTTTAAGGGTTCGTGTTACTTCCCAGCCGGCATTTCAACGGTGATTCAACGTTGAATCAACG
TCAGGTCTATGGTTGGATCAACGTTGAATTACCTTTCGATTTTGCAAATTGGATCAACGTTTATATAGTGACGTTGTTTCACCGTCGGACCACCACCGGT
GATTCACCGTTGAAATCACAACGTTGTTTCACTCTTACCTTCTTCATGGGTGAATTGTGGTCGTTTTGTAGACGTTGAATGAAGGTTGAATTACCTTTCG
ATTTTGCAAATTGGATCAACGTTGATACCGTGACGTTGTTTCACCGTCATACATTTCAAGTTAAATAAAAATTACATACAACGTAAAGTCCATAAATATA
GCAAACAAATTTTTACTAATAAAAAATACACGACACAAATACACAACTTAGCCTGTGTATCAATTTATTTAATACATACCTCATTATTTCAATTATTCTA
AATATACAGCAACATTTACAGCATACATAGTTCATTAAAACCATACAGGTTACTATACATGCTTTACACTCAAATACAGAAAAAGGCATACAATTAAATA
GTCACTTTTTAATTAGGTTTCATGAATTAAAGTTAGTTAATGTTTACTAACATGAACAAAGAACAAATACTTGTACAGCATTTATTAATCATTTAATTAA
TTGAAATATGTCCCGATGCTATTAAAATCAAAATGTATGCTTGTTAACATTAATGCACTGAGCTGACATGAACTGTACTTTCTTTATATGAAAGATGCTT
CATTGCTCTTTGTTCATGTTAGTAAATACATTAACACTTACTAATACAACCTTATTGTAAAGTGTTACCGTCTTCAGAAAGACAGATACTAACTAAATTA
AACTAAGTAACTAAACTACGCTTCAACCATCACAGTCTCCAAAATACAATTCTGTTTACAAAGTTTACAAAATACCCATTTTTAAAGTGGCTGCAGTAAA
AAATAGAATAAAAAAACATCACTCACAACTAAGAATCACAAGTGAAATACATTACATCTGTTTGAGGTTTTATAGGTTGAGAGAAACACTTGAGGTAAAA
GATGGCACTGGGACAAAGAATCAAGTGTAAAACACTTAAACATCTATATCAGTCTAGCAGTTAAATACATATTTATTTCTCAAACACCAATAGTGAAAAC
TCATGCTATAATCTAAACCTGACATTACTCCTTCACTTTGCAGGTCTGCATGCTAAATATTTAGTAGTTTTTTTTTTTTTTTTTGCATAAAGTACATCTA
TTGTGTTTAGCATTACTGCAGCAAAACGCACCTGTGACAGTGGAACATGTAGACAAATATTTCAGTAAATCATCAAGAAAATGTGTAGAATATATGGTTT
GATTCTCTCTCACAAACACAACAGTAAAAGCAATTTGTTATTCCATTATTATTAGTATCATAAAATTGCATGTTCACATGGGTCTTATAATAATGGCTCA
CATTTGTGAATGTAGGACCACTATTTCACTATTACATGTCACATTAAGTGTTCATATAGGTTCTGTTATTAATTACTTGTGTCATCCCAGAGACCTGTTA
ATCTTCAGAACACAAACTAAAATATTTTAGATTAAATGCTCCTCATCCTCTACAGACAACAATAGTCCCAAGATGATCATAGTCCAGATAAGGCACAAAA
CACATTGTCAAAACGTTCCATGTGACTTCAGTAATTTCAACTGTGAACATACCAAGCTCCAACAACACTTTAGTGAGGCAAAACAAAACAGTAAATACAA
ATGTGTTTGCGTATATCTCTCCCGCATCAGATCAGCATGTGTGTTTACTACGAGCAGTATGCACACACCATGACCTGTTGTGGCAAAGGCAAACAAGTTA
TTTTTACAGTACTTACTGTGATTTAACCTTTATGACAAATGTACTGGACCTGTTACACATATCTTTTATGAACAAATTTCACTGTAACGGATGCTGTTGT
TTTGCTCCAGTTATACCTTTAGAGAACACCGTACTAGTCAGACATCCATACAAAAGCAGAGAGTTTGGTCTTGTGTAGCAGAACTGCAATAGATGTCTGT
CTGTCTAGGCTGACACTGGTTTAGGCCATCAGTATGTCCTTGATGCCCTCAAATGAAGCTTCCAGAAAACCTGCAATAAATGAAATACGAGGTTACAACT

AAGTTGGTATCTGCTATTATACAAAGTGAAATTCAAATATTTATTTTTTATTGAATCCTAATTTGTCTTATATATCGCCACACTCAATAATTTTGACTCA
TCCATCTGAATATTCCTAACTAAGAGAAGTGCAGAATTAGGTGTGATATCGTCAGGAAGCCCTTCGCCACCACCGGTGCCAGCATTTCTAGAATTCAAAT
TTCAGATTCTCGAAAAAATTACTTTTTTTTTTTTTTAACAAAGGCTGCAGTAAAGAATCATGAAACATTTAACACATCTAATAATTACAAGTTGTTGTAA
ACCATTACATGTACTGTAAAAGTTTTTTGAAGCTTTTTAGACATATGGTTGAGAAAAACTCTAGAAAGAGTAGCACTTCTACAAAAGTCTGAATGTAGGA
CAGAGAATTAAAAGGATAGTTCAAAATTTGTATTCTGTCGTCATTTACTGACCACGTTAAACACAAAATAAGATATTTTGAAGAAAGCTGCAAATCTATA
ACCATTGACTTCTATAGTATTTTTTCTACTATGAAGGTCAATGTTTACAGGTTTTTAGTTTTTTTTTTTTTCAGAATAGCCTCTTTTGTGTTCAACAAAAT
AAACTCATAATGGTTTTAAACCATTTGGGGAGTAAAATGTGAGTAAATTTTCATTCTTGGGTGAACTATCCCTTTCAGTTGAAAAACACCAGAATTCTTA
AGTAGACCCTTTTCACAAGACAGATGATGCATTTCATTTCTCATCTTAATCCTCAAGTTTTTAATATTTAGATTTATTTAAATAGCATGTTTATTTATTT
GTTTTAATGACATTTAATAATAAAAACTTCTCAATGACATCACATGTACTGGACCTGCTACACATGTATCACTATTTTTATGAATACAAATTTCACTGTA
ACAGGTGTTGTTGTTTTGCTCCAGTAATACCTTTAGAGAACACAGTAGTAGTCAGACATCCATACAAAAGCCTTGGCGTGGAGTAGCAGAACTGCAGTCT
GTCCAGGCTGACACAGGTTTAGGCCATCTGCGTGTTCTTGATGCCCCTCAACTGGACCTTCCAGAAAACCAAGAAAGTTAAAACCTATTACACCTCTTAT
CTGCTACTATACAAGTGTAATTTAAGCATTAATTTCTCACTAAATACATATTTGTTCCCCAATACTGTCACATAGTATAACATCACAACATGATTTCACA
ATAATAATATCAATATAACAATATAATATCACAACATAGCCTAACAATCAATAAACAAACAGATTAGAAAGAGAATATCACCAACCAAAGCACCAAGAAG
ATTTAACAGATAGCTTAGTTAAAGGTAAAGTAAACTTACATGGACAAAGCCTTTTCCATTCAATGCTGTGCAAGTTAGTGCGTCTCTGGAGTAAGTTACA
TGTCTAAAAGCAAAAACAACAACAGTGAAACGAAAGCAGTTCACTTTGTAATAATAACACTAATGATACAGCGTATAAAAATGATATTGTTTTATAACGT
TGCTCAGTAACTTTAAACTGCGTTTGGAGTTTATCGCTTTGAATTTGCCATAAAAACGCCTCATTTAGACTAACGTTAAACATGTGACATTACCTCGAGA
TTAGCTGCGCGTCACCGGAGCGCAAACGACTAAAAACCAAACTGGAAAGTTTTAAATGCTTCAAAATTCGAAACA**GGTCCGTGTATCATCCGTTCATTTG**
**ATTATTCCTTTTATTACGGAAAACGAAACATCAGAAAAACGATGAATATTTCGTTTTTCTGTTTATTCTGTAGGCAGAAAAAACGGAAACTCATCTGTTA**
**TTCTGCTTATTAACTTAAAACGAAATATTAAAACAAACACAAAACCAAAACGAAATAATGAGTCAAAAAATGGTCCGTGTACGGTCCGTGTACATTTTGT**
**TCATTTGTTTTCCATTTTCAAACGGAATAAAGGAAATGGAGAAAACGGCCGTTTTTCCGTTTTTTCTTTTGCTCACGAGAAAACGAAAAAACGAGATTTT**
**GGCTGGATTTTCGTTTTTTGGTTTAGGGTAGGAAAACGGATAAACGACTTTAAAATTCATTTTACACATGTAGGCGGTACTGAAACGCCCACTTTCCTCT**
**AATTGGTCAACCAAATCTGCGCTCGTGACGTCACCTCAAAATAAAAGCCTTACACGCAGGCTTTTAATTATTATTATTTAATTATATATATAAACAAAGT**
**TTACATATTCTATCATTTGAACCACACACAGTTAGCAGGCTTACATTCATTGCGTATGTATAAATTAAATAAAAACGTAAAGCAGTATTCTTAGACATTT**
**GTCATTAAAATAAGGTCATAGTTCACTGCCAAGCTTCTTGCTGCTGTGCACCTGATGCAGAGCAAAGAAAGCCATTTCCGAGGAGCACCTGGTTTACATG**
**ATTGTTTCAGAGCTATTGTAGGCCTAAATAATAGCCTACTCTGTATAAAATAATAATTTATTATTATTATTATTATTATTATTATTATTATTATTAGGCC**
**TATTATTATTAGTATTGTTATTATTATTTACTTAGTTATTCAATGGTTGTAAGAACACACATTGGACCCTGCATGGGGCTATAATAAATGGTACCATTTG**
**TTTCTATTGGATTTTCTCCTATTTTAATTTAATATATGCCTTTTGCCTATTGTAAATATTTATTTGATAATTAATACTTTTTTATTGCAACGTCAAAGTA**
**GGCCTGTCTTATTATTATTATTTTTTGTTAATATTTTGTCGTTGTTGTTGTTATTATTATTGTAGCAGGGACATAGCCACAGGTGTGGCAGAGTGTGCTG**
**GTGCCACCCAAAGTGGCATCTTGCACCTGAAAATAGATGCCTTAGCGCTCAAGCGCGCGCAAGAAACTTGCACAGGCAAAAGTAATGATGAATGTGTTGA**
**AGGAAAAAAAAACATTCTAATTATTTGTTAATAAAATAGTGCAACATTATTCTCAGTCAGTGTAAGCTGAAAAAATTAAGATCGCCAGCATTTCAAGGGT**
**GGTTTTATTTTTAGTTCATTGCTGTCACGCGCTATATTTCACTAAGGCTGCATGAGACTGCGCATCTTGCTTATTTTCTCTATTTTACATAAATAACATA**
**TCATACAGTCCAGGTTCTACCGTTCTAGTGAGTGTGGGGCATTGCTTTACAGTGGTAAAGTGGATTATTTCTTGGCAAATTCATAACGATTCACTTTGTT**
**TAGTAATAAATTTCATAGTTATAAAAATAACTACATTTCAAGAATATGTTAACACTTTCAACTATTTAGGATTATTTATTGCATCAAAAGTAATTTCAAA**
**GTAACATTAAATGTACGTATAAACTGCTTAATTTGTGTGTGTAAACACCTAAGATGCAAAAAGCTTATTTAGACCAGAATAGGCCTACTAAATGCAACGG**
**TCAATGTATGTAAACTTATGACCTAAAATGTCTTAAATGACTAAATTTTAAGTCTTTTTTAAGTTTTTAAATAATTTAAGTTTAACATAAATGATTTAAA**
**CTTTAAACTTCAAATTAACAGAGTATGGCAAATAAATTGTTTATTAAATGGAGGTCAGGTGTCTATCTTATTTATAATTATTAAATATTATGACACTATA**
**ATTCAGTGGTGTATTGGAGAGTACGCACCTTTTAATTAATTAATTTTAAATAAGTGGTTTATCCTAAATAAAATGCAATGGGATCGGTGTGCAATGAATA**
**GTTTGAAAACCCCTGGTTCAGACTAGACGAGCTAGCTGTCTGTGCGCTGCGCTAATAACTGGCATTTGCTCTTTTCGGTGCTGCCAGATGCATTTATAGT**
**GCATAAAAAATAAAAAGATAATTCGGTAAAGCGCGCGATGCGCGCAGAATTAAGCGCACTATACGCATATGTCGAATTTAAGTTATCTTTTTCTTTAATC**
**AAGACTTTTCCATTCGAGGAATTACAGTTATTAATAGCTTAGTCCTTACAAACGATCAGTTTATAAAATACACAATTTACGGATCAAGCCATAATGAAGA**
**ATAAGAAGAATATGCAACGCATAATCCTTGAATGATCAGAGCGCATTTATATAAACTAGCTATTTTATATAATCCATAAATATTAAAAAACATTATACAT**
**TTAAAGTAATGGAACATTATATGTTATAAAATCTAAACTATTATTTAGGCTACAGGCTAAATAAAGGCGCCTTTTCGGGGCTGTTCAGCGCGTGCGTTAA**
**AGAAAAGACATGACACAATCTCTATAAAAACTTGTTGTTTATTTACAATATCTAAAGGGATATAAAACATCTAACATGCATCTCCATTCTCAACGAGTTT**
**CTGTGTTCTGCATCCTTCACAGACCGAGGTCTCATTCCGAGGTAATCTGTAAATATTCTCATTTGGTTCTGGCACATTCTTGATTAAAGGTGAAACTGAC**
**CGGCGCAACGGATGATTTGTCATCTAGAGTGACTACAGTTATAATACAGGTTATGGAACTATTTCACTTTAATTTCATATTATTATTTTATTAAACAAAC**
**AAAGCAGTTTTTGTTTTGGCTCTGCATGCAGCGTATTGGATCTTTGAAGCATTGCACTTCAAATGTTATTCCTTTTTTTTTTTTTTTTGTATTTTTAAT**
**ATTTTTAATTATACAACACAAAATTGGTTCTAAACAGATACATTTTCCTATAGGCTATGATGAAGTGTATATTTTTGTTCACACAGGTCCTATACAAAAC**
**TGTGTGGATGGATGATTTGTTTCTCCGTGAAAATCACTCATTTAATAAGCAAAACAGGCCAAAATGTTGTTTAGCTTGCATCAAACTGGATGGACAATTT**
**ACAAACATATTGAATACAAAGGCTGGTTATTTTATGCAATGAGGGACAATAGGCCTATTACAAATTAAACGAAAATTATTTGCATATTAAAACGAAAATA**
**TATAAAACAACAACAGATCCTACCTCGTAAGAAATGACTTTAGATAATGTAAATAAGCATCAAGGTTTTTTTTTGAGATTGTGTGTCTAATTCAGTAGTG**
**TAGTGCGTTCGCGTGGGTTTCCTCCGGGTGCTCCGGTTTCCCCACAGTCCAAAGACATGCGGTACAGGTGAATTGGGTAGGCTAAATTGTCCATACTGTA**
**TAAGTGTGTGTGTATGTGTGGATGTTTCCCAGAGATGGGTTGTGGCTGGAAGGGCATCCGCTGCGTAAAAACTTGCTAAATAAGTTGGCGGTTCATTCCG**
**CTGTGGCGACCCCAGATTAATAAAGGGACTAAGCCGACAAGAAAATGAATGAATAATGTGCAATACTAATTTAAAATACTTATCACTAACCTTATATGTG**
**AAATAGCAGGGCCCTCTGTAGCACTTTCTCAAGAAGAGCAAATGTCAAATTATTAGTGCAACGCACATGTAGCGAGCTCATCTGTTTGTCGGAACTACTA**
**GACACAATTTTTTTTTATCAACTATAATGTGTTTAATTAGTTAATTTGAAATTTATTTCATTATTCCAGCAATAATTGTTGAGTGAAAGAATGCGCTAGA**
**AATATGCATTGCGGCTCCCTTTATTGTAAAGGCTTATTGCACTTAAACTTTTTTAGGTTCGGCTCTCGAATTAGTAAGGTTATGAGTAAATGTTTATTTA**
**AAATATTTAAAGCCTGCCATCTAGCCTACAATAACAGCAAAATTAATAATAAAAAAAATAATAAGACAGGCCTACTTTGGCGTTGCAATAAAATAGTATT**
**AATTATCAAATAAATAATTACAATATATAATTAAAATAGGAAGAAATAAATGAAAACAAATGGTACCCTTTGTTTTTATGCAGGCCCAAAGTGTTCTTAT**
**TCTGGTTCTGCTAACAACTATTAAAAAGAAAGAAAAAAAAACACACCAAACAATAAAAACACTAGTAACTGATAATAATAATGATAATAATAATAATAAGT**
**ATTATTATTATTATTATTATTATTATTATTATTATTATTTATTATTATTATTATTATTATTATTATTATTATTATAAAGCCCTTGTTCTGAAACAATCATGC**
**AAACCAGGTGCTCCTCGGAAATGGCTATCTTTGCTCTGCATCAGGTGCACAGCAGCAAGAAGCTTGGCAGTGAACTATGACCTTATTTTAATGACAAATG**
**TCTAAGAATACTGCTTTACGTTTTTATTTAATTTATACATACGCAATGAATGTAAGCCTGCTAACTGTGTGTGGTTCAAATGATAGAATATGTAAACTTT**
**GTTTATATATATATAATTAAATAATAATAATTAAAAGCCTGCGTGTAAGGCTTTTATTTTGAGGGTGACGTCACGAGCGCAGATTTGGTTGACCAATTAGAG**
**GAAAGTGGGCGTTTCAGTACCGCCTACATGTGTAAAATGAATTTTAAAGTCGTTTATCCGTTTTCCTACCCTAAACCAAAAAACGAAAATCCAGCCAAAA**
**TCTCGTTTTTTCGTTTTCTCGTGAGCAAAAGAAAAACGGAAAAAACGGCCGTTTTCTCCATTTCCTTTATTCCGTTTGAAAATGGAAAACAAATGAACAA**
**AATGTACACGGACCGTACACGGACCATTTTTTGACTCATTATTTCGTTTTGGTTTTGTGTTTGTTTTAATATTTCGTTTTAAGTTAATAAGCAGAATAAC**
**AGATGAGTTTCCGTTTTTTCTGCCTACAGAATAAACAGAAAAACGAAATATTCATCGTTTTTCTGATGTTTCGTTTTCCGTAATAAAAGGAATAATCAAA**
**TGAACGGATGATACACGGACCGCTGGCAAGGCATGGTTTGATGCTGGTTTTTGCTGGCATTTTGTTGGTCAAGGACTAGCACAAGACCAGCAAAAACATG**
**AACTAGTAAGGAACCAACATTGTAACATTCCTTACCAGTAGGCTTATATGCAGTATTTTTCTGCAAGGAAGGACCATTTATGCAATCAAAAGATTCTGAG**
**TTAAAGGTTCTTATATGTAATTATTAATGCTAATAAATAAACAAATACAAACATTTGAAGCCAACAAAAACTATGTAAGGTCAGTGTAGGGGTCATCATT**
**TTTTTGGCAGTTTGTATAATATGCATTGATTTATCAAACATAACTTTTTAGGCATAATGAATGGATTCAGAAGCAGCGTGATTCAATTCAAGTTTATTTA**
**GTTTATTTTCTTCATTCATTAATTTTCCTTCAGCTTAATCTTTTTCTTCATCAGGTGTCACCACAGTGGAATGAACCGCCAACTTATCCAGCATATGTTT**
**TACACTGCGGATGTCCTTCCAGCCACAACACAACACTGGGAAACATCCATACACTCTTGCATTCACACACATATACTAGGCCATTTTAGCCTACCCAATT**
**CACCTACTGTATGCCGCATATTTGGTCGGTGGGGTAAACCGTAGCACCCGGAGGAAACCCACACAAACACGAGGAGAGCATGCAAACTACACAGAAATGC**
**TGCTCAGCTGGGACTTGCACCAGTGACCTTCTTGCTGTGAGGTGACAGCGCTAACCCCTGAGCCACTGTGCCACATTAGTTTATTTAGTAACAATATTAA**
**TTGTTTCAAAGCAGCTTTACAAAAGGTGCACGCATATTGCATTCAAATCAAAATCAGAAAAGATAAGAAGATGTGTATCGGGCAGACATCGTACATGAAC**
**ATATTTAACCTGCAGTTATATAGTGTACTTCCAAAAAATGTTCAGTGAAGGGCATTAAGTGTATGTGCTGTCCTCAAAGGCCTTGGATGCTTGACATTAT**
**CTCTTCACAGGTCTTGGATTTGATCAAGTGGATCTACACAATCTCAGGAGGCCTCAGAATGAGCATCCTAGATGGAAATAGAGAGCAAAGATAATCATTA**
**CCGTAGCTGTTCATAATGTATTTAAACAAGAGATAGTATTGTCAATGTGAGAATGCTTTTGAAGAGATGCATTTCACTGCATTTTCAAATTCAGATCTTG**
**TTTCTTTTATTTTCTCCCAAAGACATTTAATTATCAAAAATGAATTTGTATATAATATGACCAGTCTCACCAAACAGTGACAGACATGCATCCCTGCACT**
**GAAAAATAGATTATCCAAAAAGATTGTGCTTGCTGTTTTATTGTTTAACTTACATGTTTAAGCAGATACAGTATTTTTTTATATGCAAAAATTATTACAA**
**ATATTTTGCCTTACTTAAAGGCAGATCAACCCCTATCTCCAAAAACTAAAAAAGTACCATCTATTTTAAAAAAAGTAGAAATAAACTTTCATTTTCTTCT**
**TTTCTACACAAACACCAGTGAAACGTTCAGCATTTAACCACCCTAAATGGAATATTATAGACTGTATATGGCACATGCACACACCTTGACCTGTACTGTG**
**ATAAGATTCTGATAAGAGCTTTAGTTGTGTTAGAACTGAGGTCAGTCTTCTCATATCTCAAAATATTTGAAGTTCTTTTAAACTTTGAGCTCAATTTAAA**
**AAGTTTTATATGTAGTTAGAAAATTTTTACATGTATAAGAATAGTCAATTAGTTATTTATTTTGTGAGCGCGCATGAAAGAATTGGGTTTCTGGGAATTG**
**TAAAACTTTACATAGACTGTTCACTGATCTAACTGTTTTCCATCCGTTATATATGAAACCTTCCCAGTTTCCTGATTATTTATATTAAATACAAAAAGAT**
**AACAATTAGCAGATAATTTTACCATCCTACTTTGTCAGACTGCACATTTATTTATTTATTATTTGTTTGTTTGTTTGTTTGTTTAATTATTTTTTGTTTG**
**TTTGTTTGTTTGTTATGATTTGGCTCATCTACAAACGTTGTTTGCCCTTTAGAAGAGCTCAAGAACAGTGCATTAAGAGCTTAATTTTCATTGAGAGCTT**
**AATAATAATCTCAAAAATATTGTTCCAACATCTTTAAAAATTAATATTTTCTTCATTGACGGCTCCACAAACAAGTTATAATTGCATAATAATAAAGCCA**
**TTCAAACAATACCAATTTCGTTTTTTTATATTTTTCTTGAAGCGTTTTCAATGCATTGAGTGTCAGACATTGCCATCTTGAGGAATAAGGCTGCATTGCA**

ATTATTAAGTCTTGCACAGGACAAAATATAGTATATTCCTTAGAGATAGATACATTCTTTAATAGCAACATTTTATTTTTAGGCTGTTCATAAATAAGTT
CATAAATAATGGAAAACGCTGTTGATATTTGTAATCACCAAAACAAACATAGGCCCTATCAAAGGATCACACCCATATACTGTACCCACAGTCATGTACA
TGCTATGCGACACAGACAAAGAGTAACATCACAGTGCTGGAATCTAATCTGTTGTGCATGTCACCCGAAAACATAATTTTATCAGTAAGTGAACGAATTT
TATTTGTGTACTGTATGTTATCAACAGCTTACTATCAATGCTGGAGAAATTATTCTGAAACTGTAGTTTGATAAGCTACAAGCTAACCGGCAGCAGCAAA
GCTTCAGTCAAACTATACTTTTAAAATGTATCTGTAGCTACACTACAAGTAACTTCCTCCAAGTAGACATTGAAGCTATGTTTTTCACAAAGGTGTATTT
TACAAAATGTTAGTACAACTTTAAACTTAACGTTTTATTACACATAGTTTTCAAATTTCTCAGACTGTTTAAACTTTAAAGTCAAAGTACAAAGCTGCAA
CCAATATGGCATTGACTTAGCAAAACAAATTAGATTAGAATACTATATTGAATATTAACTGCTGTCACTTGTTATTATTCATTCATTAATTTCCTTTTCA
GCTTGGTACTTTTATTGATCAGGGGTCGCCACAGTGGAATGAACCGCCAACTTATCCAGCATATGTTTTACACGGCAGATGCCCTTCCAGCCGCAACCCA
ACACTGGGGAACACCCACACATAGACTACAGCCAATTTAGCATACCTAGTGCACCTATACCGTTTTTGGATTTGTGGGGGAAACCGGAGCACCTGGAGGA
AACCCACACAAACACAGAAAGAACATACAAATTGCACACAAAAATGCCAACTGACCCAGCCTGAACTTGACCCAGCAGCGACCTTCTTGCAGTGAGGCGA
ACGTGCTACTTGTTATATAATATTACTTAATGTTATTTAATTTAATAAGAGTAATACATGAATGCAAGTCTATGATATGTATGATTAACCAATATGTGGC
TGCAGTACACAATCATATGTGATCTTGTAACATTGTGCTGCCTCAGACCTTTGCAGGTTCACTACATAAAATGCAGCTGTTGTCAAGCTACTGCTACTTT
CTGGAAAAAAGTAGCTGCTGGAAAAGTTACACTGTATCTTAAGTAGCTGAGCTACTGTCAAAATGTAGCTAAACTTGTAGCTACCGAAGACAGCTCCCTT
TATTATATGAGGTTAAAAAGCTAATATTTTACAAATGTGACCTGATTTAGCTTTTTTATAGCATTCATTAAAGGAGAAACAACAGTTGTAATCATTTTTA
TGATTTTACTTGTCTTGCTCTTTGGACACACTTTGGGAGCCAATTCTTGATCCAGTGAAGGAGGTTTAAATTTCCCTCTAAATTAATGTCTTTTTACATT
AACTCTGAAGTGAAGGGAAATAATAAACTATTATCTTATGTTAAGAAACTAAACATTAGTTCATTGTGTGTTTAATGACCTTATCAAATACTACACTAAA
AATGATTTCTGCTGCTTGTTCAAACTATCTGTTTAAAAATGAGCTGAAACAACACAATTCTTGTCCTTTCTTTGGCCAATTTATTTGTTTAATGTTCAAT
CCACTTAAACTTGTAAACCATTAAGTTAACTTAAGTGTTTTGTGTTGGGACAACATGAAGGAATTGTAATGGTCCAATACCTGGTTAGTGGACTTGTAGT
TCCCAGTGTGCTTTGCATGGGACTAAATTAAGAGAGGCAAATGTTGACAATAAAAGTAATCTTAGGTGTTTAAAGTTAATAGAAAGACTTTTTAGAGTTT
AATCATCCCTTTATTTTTAAAGATTTAAAAGATTTTCATGTAATCCTTTTGAGTTTTGAGATAATCACCTTGCAGAAGCACATATGCTTTGGGTGTTGGC
ACCTTAATTAGCAAAACTGCAGTTTGTTGCTTTGCTCAGTAAGCAAAAACTGTCCTAAAATGAGTTCTGAGATGAGTCTCAATCAACTGTACATGTAACT
CGTGAAATAAGCTTAAAATTGCACTTTTAGTTAATTTAGGATAACTTTAAATAAATCAAAGAGACTTTGAAATATAAGACACATAATAGACACACATGAT
ATTATCAGACCTTTAAGTTTAAGTTAAATAAAAATACAAATGTATTTAAACTTATATTTGATCAAATTATGTTGAACCAACTCAAATGCATTCAATTGTG
TAAACTGTTTTTTTTTTTTTTTTAGTTGGTGCTACAGAAATTGATAGGATGGAAATCCTACCCTCAGTTACAGTAAATTGAGTTCATCTGATGAGTTATT
TTTAAGTGTACTTTCTTATTTTAAACATGCTTTAGTACATGCAATAATAACATTAAAATTAACACAATTATATAATATATAAAATATCATGGTCAGCGGT
GGTGCAGTCGGTAGTGTTGTCACCTCACAGCAAGAGGCTGGGTCAGTTGGTGTTTCTGTGTGGAGTTTGCATGTTCTCCTGCGTTCGCGCGGGTTTCCTC
TGGTACAGGTGAATTGGGCAGGCTAAATTGTCTGTAGTGTATAAGTGTGAATGAATTTGTATGGATGTTTCCCAGAGATGGGTTGCAGCTGGCAGGGCAT
CTGCTGCATAAAAAAAACATGCTGAATAAGTTAGCAGTTCTTTCTGCTGTGGCGACCCCGAATTAATAAAGGGACTAAGCTAAAAAAGAAAATGAATGAA
TGAATAAAAAAACAATTCTGTATTTAAATTAATAATTTTAAGTTAACAAATAAAACATCTTTATATTTTATATGTATTTATAACTTTGAATGTTGTGTGC
AAATGCATGGAAATCATCTGGAGGTCACTTCATTCATGTTATCCAAACACAGTACTAGACACAATTTCTTAGCTTTTGTGATATACCAGTTGTGAGAATT
ACAGCTTGAGCTCAACCTAGTCTAACTTACAATCGCGGTGTGACACTTTAAATCGGATTTGCGCCCTCCGAACACTAACACATGGAGAGGTAACATCAGG
ATGAGTGAATCATGCGCTCCTCCACACCCTTGACTCTCCCCCTCCCCAACCATCCTCTCACTGCCTCTCCATATTCACTATCAATCTATGCAATTTTAGT
CAATTTTATTTTTAAACGCTCCCAAACACAGAGAGGGGCTGTTTCGCAGCGCGTCGGAGAAAGTGATTGAGTGAACTGGGACAGAAAGTTGATTTGCATC
CATTTGCAAACAAAAAGGGAAGCAGGTGACGTGAGGTAATTGGAGAGACGGCTGATTTACTGAGCTCGAGGATGCGCGCACTGACAGATTTAACACGTAA
TCACTGGAGACTTTGAAGAGACGCGCTCTCGTGCTGCTTTTACATCGATGTGGTGTGAAGGTAAAGCTTGTGTACATGTGTAACACTTCGCGTAGAGTCG
ATATATCGCTCTTATTCAAGTAAATACAAGCTGTCGAGGAACTTGCAGAACTTTCTTGCCCTGCTGTGGTAGGACAACATGCCTGTACGACGGGGACACG
TTGCGCCTCAGAACACCTTTCTGGACACCATCATTCGCAAATTCGAGGGTCAAAGTAAGTGAATGACTTTTATTTAGTGAACGTGCATTGATTATGTTT
TTAAAACAGTTTCTAATGCTGACTATTGTTATTATTAAATATTTTGTTATTAATATTATTTTATTATCAATTGCCATTGAAAATTACTAATTGTCACTAA
TAATAATGTCGTTAATAACTTAATTTGAGTTAAACGTGTCAGAACAGCTAAGTAATCATTTACTTTTGCTATCTGAATATGTAAAAATAGTTAATTAAAC
ATAATTTTACTTTTATTATTATTTTTTAAAGTAGCCTATGTAAAACTAATTGATTTCAGATTTTCAGTAAGCTGCTGTTTCAGTAGATCTTTTAATGTAG
CTAAAGCAGACACATTTATTCTGAACTGCAGCTCAATTTTGATTCTGCATAGATGTGGTTAATAAAATTTTTGCATTTCTATTTTTCTTGTACCTCAACA
GATCGCAAATTCATCATCGCCAATGCTCGAGTTGAAAACTGTGCCATCATCTTCTGTAATGATGGCTTTTGTGGGATGTGTGGCTACACGCGTTCAGAGG
TCATGCAGAAACCGTGTACATGCAGTTTTCTGTATGGCCCACACACTGGACGACCAGCGGTGGCCCAGATGGCCAAAGCCTTACTGGGCTCTGAAGAAAG
GAAGGTGGAGATCTCCTTATACAGGAAAGATGGTAAGTTTTTTTGGAGAGTTTATTTAATTTGTGTGTTTCTTGGGTGCCAAAAATGGCAAACCATATTG
CTAAAACACATAAACAGTCAGAAACAAGAACTGTGACTAAACATCCCACTTTTACCAGGACAGCTTAGGGGCCCTGACTTAAAAAAGATCATGTTTTGTC
CTGTATTCACCAGTCTTAAAATGCCTTTATTGCTGAGGGAATGTAGAAGTCTTTTACCACAAATACAATTTGTCAAATCAAAGGTAGCCAGTCACATGAT
CATAGTAAATATATGTGACTACGGACCACAAAAACAATCATAAGGGTCAATGTTTAGAAAATGAGATGTATAGATCATCTGAAAATTGAATAAATTAGAT
TTCTGTTGGTGTACACTACCTGACAAAAGTTTTGTCGCCTATCCAAGTTAAGAACAAATACTAATGTAACTTCTAGTTGATCATTTGGTATAAGAAGTGG
CTTATATAAAAGGCAAAGGCCTCTAGATTACAGTTATTTGACAAAAAAATTTATCTATTTAATTAGGAGAGTAAGGTTTGACTTGGCTTAGACAAAAGTA
CAGTATTAAATATAAAGTCATTTGAATATAAAGCCATGAAGCAGTGGAAAAAAAATAATATTGTGTTTGACTCCCATGAGCTTGGATGACTGCATCCATA
CATCTCTGCAATGACTCTAATAACTTATTAATAAAGTCATTTGGAATGGCAAAGAAAGCGTTCTTGCAAGACTCCCAGAGTTCATCAAGATTCTTTGAAA
TCATCTTCAATGCCTCCTCCTTCATCTTACCCCAGACATGCTTAATAATGTTCATGCCTGGTGACTGGGCTGGCCAATACTGGAGCACCTAGACCTTGTT
TGCTTTAAGGAACTTTGATGTGGAGGCTGAAGTATGAGAAGGAGCGCTATCCTGTTGAAGAATTTGCCCTCTCCTGCGGTTTTTAATGCAATGTAATGTA
ATGTAAATGTAAACCTCAGGCTGTTGATGTTGTCATCCACTTTGTAGATCTCTCACACGCCCCCATACTGAATGTAACCCCAAACTATGATTTTTTCCTC
ACCAAACTTGACGGATTTCTGTTATAATCTTGGGTCCATGCTGGTTCCAATAGGTCTTCTGCAGTATTTGTGATGCTTGAAATGCAGTTCAACAGATGAT
TCATCAGAAAAATCTACCTTCTGCCACTTTTCCAAATGATCAACTAGAAGTCAAATTATTATTTGTTGCTCTTACAACTGATATTTATGACAAAACTTTT
GTCAGGCAGTGTATTGTTTGTTAGGAAATGACATTTGTTTGATTTACTACTGTATTTGGAAATCTGAAATCTGAGGATTAAAAAAAAAAGCAAAAAAAAA
AATGAGAAAAGCAACGTATAAAGTTGTCTAAATCAAGCTCATGCTCTCTGAAATTAAGTTTTTGATACATCTATGGTAGGAAATTTAACATTCAATGGAA
TGGAAGTTAAATGGAACAGCATTCTAATGATTTTGGCATAAAATTTAAATCAGTAATTTAAACTATTAAAAATTAATTGTTGGCTTTTCCTACAAGTAAA
CCTTTGCAACCTAAGACTGGAAGAATTACCATCCAATCACGATTTTTCATCAATTAATTTGCTTTAATGAAGGCAGGTCAAGCCTGAGTGACAGCAATAG
AGAGCACAGAGAAGATAGAGTAGTACACAATACTAGTTACAGTGCAGAAAATACAATCATAAAACCTATTTAAAAACATTTTTTTTAAATCACAGACTAC
ATCAAATTCTAGAACAGCTTACATTAATACCATGTGTAGATATATGCAGTGCAGTGTATCTCTGCCTAAATCTGGCAACCACAATAGCAAGTGACTTTGA
CCACCAGTTTGTCAGTAATTACACTTTTGGTGGTCTTATTCTCATTCCTTGTTATTTCTTTTCTGCTGAATTCCTCTTTGTGAAATTTGATGCGTCCCAT
GGTGACTGATCTAGGAGTTCAGCCTTGCTCCTGGAAGTCTCTCTCGCCCAAACATCCTTTATTGTGGATGCTCAACTGAAAGATGAGGATTACAGTAGAA
GTGCTATTCTAATTGGGTCTATTTCTGGCCTCTCCTATTTAAAGGTAGCATTTCTGGGTGATGTATTGATTTCCAGGACAGCTGAGAGTTTGAGAACAGG
AATAGTCATAGTAGACATAATTGCAAAGGACAGTTTCTCATCATCCAGATCTATTCCCACTGCTATATGTTAGTAATCCAAATGGTTGCGTAAGGGCATC
ATAACATTGTGTCCTCTAGAGGTTCCTGTATAGCCCGTATTATTGTTTAGAGGAAGAATATACCATATCATGTTAATAAAGATGATAAACATGTAATTCG
TCCTAGGGCTGCACGTCCCAGCGGAGAGTCTTTTTCCACTGTTCTGGATGTTAGGAATACTGCTCATTGTGACCCTTTGTAACATAACCAAGCTAAATTT
CTCAGGTGTGCTCAGGCTACATTTAGACTTTCCGAAATATGAAACCTCTTGATTTTCTTCTTTGGATTGGTCTCTAAGTCGCATTACAATACAGTATGTT
GTTGGTTAATTTAAGTTTTAGAATATGTATGTGTACAATCATGCATATATTTAGACGTTTATAGTAGTTTGTGTCAGTATGTAAGCCTTTACTTTCTTAA
TTGTCCACAAGTGGAAATTTATTTTTGGCTTCACCACACAACCTCATCAGCATTCACATGACACATATCACAGAAAAACAAGAAACTCAAATACATTCAA
TGTCATCACATAAATCCCATTAAAAACTCAGTAAAAACCTTGTACTTACGTATAAGCCTATATGTCAGACTCATCAACCTGCGCCTAGCAAGATTGTCAT
TGTATGGTATGTTCACTATGTGTATTGTATATAATAGCCATATATATAAACTGTTGTTGTGCAGATGCCATGTTTGAAAGTTTAAGCATGTTTTGGGAAG
GATGTGTAAAATGCAATATGCAAATAGTTTCCAGTACAGACAGAGGTTGGGGAAGGTTATAACATCTCCATTGCAATAGGCAGTTTGCCAGCCTGTGCAC
TTTAGTTTATAGCATATATATATTTCTTGGATGTATGTTGAAATATGGAGGTATAGACAGCTGGCAACATTCATTCATTAATTTTCCTTCGGCTTAATAC
CTTATTTATCAGGGGTCGCCACAGCGGAATAAACCAGTGGATGCCCTTTCAGCCGCAACCCAGTGCTGGGAAACACCCACATACACTCATACAGCAGGCA
CAGGACATCAACATGACAACAGATTGATGTTGTATCCCAATGTTGTGGGCATTTTACTTGGAAATGACAATCTGATTGATGTTAGAACCAACGTCAATGT
CCAACGTAGGACCGACGTTCATTTTTGGTTATTTTCCAGTGCAACCTAAAAACAACAAAATCTTAACGTCTAATTGTGTTACAGCTTGACATTGTGGGAC
ATTACCAATATGATGTCTATCAGATGTTGGATTTTGGTTGCCATACCTGACGAATAAATATCAGTATTTGACGTCAATATAACACTGGTTTAAGATGTTG
GCTTGACGGTGGATTTTCGTCACTTTCCAACACAACCTAAAATCAGCGTAATTTCACGTCGTTATTGGACAATAAAATAACATTGTCTTTAGACACTGGC
AACTTTATTCTAACCTTATATTAACAATGTGTGTCTGCTGGGACATGTCATTGGACTGTCAGGAAAACCAGAGCACACGAAGAGAGCATGCAAACTCCAC
ACAGAAATACCAACTGACCCAGCGGGGACTCGAACCAGCGACCTTCTTGCTTTGAGGCAATAGTGCTAACCACTAAGCCACTGTGCCGCCCCAGCTGGCA
ACATAATCTAACTATGCATCATCTGATTGAGGTGTACACAGTTTAGCATGTTTATTTCCTTATTTAAAGAGTGTGTACAAAAAAAACATATTGCTCACGC
AATCCCACTGGTCCTATACCACCCAACCCGCTCTGAGCTGGGAATGAACCGGCGACCTTCCGCATGGGAGTCGGTTGCTCTAACAAGGAGGCTAAAGACC
ATGGCCTCTGGTCTGTCGCTAGAGCACCTTTAGAGTCAGAGGAGTGAGGTTTACCTGAGCAGCACTTACTACTGGCCTCTGTTACACTCACCCCCTAAAC
CTACTTCCATCCGGGACACGGCACAATGTAACCCAACTGGTCCTACACAACCCAACCCGCTCTGAGTTGGGAATGAACCGGCGACCTTCCGCATGGGAGT
CGGTTGCTCTAACAAGGAGGCTAAAGACCATGGCCTCTGGGTCTGTCGCTAGAGCAGCTTTAAAGTTTAGAAGAGTGAGGTTTACCTGCACAGCACTTAC
TAGCTGGCCTCTGTTACACTCACAAAAA

#### S1 Table. Beloniform pectoral fin measurements

| species | group | MCZ Catalog No | SL (cm) | pectoral (cm) | pectoral/ SL |
|---|---|---|---|---|---|
| *Hirundichthys rondeleti* | flying fish | 722,6223,27301,27302 | 23.0 | 17.4 | 0.76 |
| *Hirundichthys rondeleti* | flying fish | 722,6223,27301,27302 | 19.5 | 12.8 | 0.66 |
| *Hirundichthys rondeleti* | flying fish | 722,6223,27301,27302 | 23.0 | 15.8 | 0.69 |
| *Hirundichthys rondeleti* | flying fish | 722,6223,27301,27302 | 18.2 | 13.0 | 0.71 |
| *Hirundichthys rondeleti* | flying fish | 156319 | 12.7 | 9.0 | 0.71 |
| *Hirundichthys rondeleti* | flying fish | 156319 | 11.2 | 8.2 | 0.73 |
| *Hirundichthys rondeleti* | flying fish | 156301 | 4.9 | 4.0 | 0.82 |
| *Hirundichthys rondeleti* | flying fish | 156301 | 2.7 | 1.7 | 0.65 |
| *Hirundichthys rondeleti* | flying fish | 156301 | 3.1 | 2.3 | 0.75 |
| *Hirundichthys rondeleti* | flying fish | 156301 | 2.4 | 1.8 | 0.76 |
| *Hirundichthys rondeleti* | flying fish | 156301 | 2.9 | 2.1 | 0.72 |
| *Hirundichthys rondeleti* | flying fish | 156301 | 1.7 | 0.9 | 0.54 |
| *Hirundichthys rondeleti* | flying fish | 156301 | 2.3 | 1.4 | 0.61 |
| *Hirundichthys rondeleti* | flying fish | 156301 | 2.6 | 1.8 | 0.70 |
| *Hirundichthys rondeleti* | flying fish | 156307 | 3.0 | 2.0 | 0.66 |
| *Hirundichthys rondeleti* | flying fish | 156307 | 2.9 | 1.8 | 0.63 |
| *Hirundichthys rondeleti* | flying fish | 156313 | 6.0 | 4.8 | 0.80 |
| *Hirundichthys rondeleti* | flying fish | 156313 | 3.2 | 2.1 | 0.64 |
| *Exocoetus volitans* | flying fish | 714,5208,6207,2415,4482 | 17.5 | 12.5 | 0.71 |
| *Exocoetus volitans* | flying fish | 714,5208,6207,2415,4483 | 17.5 | 13.0 | 0.74 |
| *Exocoetus volitans* | flying fish | 714,5208,6207,2415,4484 | 17.0 | 12.3 | 0.72 |
| *Exocoetus volitans* | flying fish | 714,5208,6207,2415,4485 | 16.2 | 12.2 | 0.75 |
| *Exocoetus volitans* | flying fish | 714,5208,6207,2415,4486 | 16.9 | 12.2 | 0.72 |
| *Exocoetus volitans* | flying fish | 714,5208,6207,2415,4487 | 16.5 | 11.0 | 0.67 |
| *Exocoetus volitans* | flying fish | 714,5208,6207,2415,4488 | 15.0 | 11.0 | 0.73 |
| *Exocoetus volitans* | flying fish | 714,5208,6207,2415,4489 | 14.0 | 9.4 | 0.67 |
| *Exocoetus volitans* | flying fish | 714,5208,6207,2415,4490 | 17.5 | 13.0 | 0.74 |
| *Exocoetus volitans* | flying fish | 714,5208,6207,2415,4491 | 13.5 | 9.5 | 0.70 |
| *Exocoetus volitans* | flying fish | 156369 | 10.1 | 6.8 | 0.67 |
| *Exocoetus volitans* | flying fish | 156369 | 11.8 | 8.0 | 0.68 |
| *Exocoetus volitans* | flying fish | 156550 | 4.6 | 2.9 | 0.63 |
| *Exocoetus volitans* | flying fish | 156550 | 3.4 | 1.8 | 0.53 |
| *Exocoetus volitans* | flying fish | 156550 | 2.9 | 2.0 | 0.67 |
| *Exocoetus volitans* | flying fish | 156550 | 3.2 | 1.3 | 0.41 |
| *Exocoetus volitans* | flying fish | 156550 | 3.5 | 1.9 | 0.54 |
| *Exocoetus volitans* | flying fish | 156550 | 4.3 | 2.5 | 0.58 |

| | | | | | |
|---|---|---|---|---|---|
| *Exocoetus volitans* | flying fish | 156550 | 3.8 | 2.3 | 0.59 |
| *Cheilopogon furcatus* | flying fish | 35001 | 24.0 | 18.0 | 0.75 |
| *Cheilopogon furcatus* | flying fish | 49165 | 23.0 | 16.0 | 0.70 |
| *Cheilopogon furcatus* | flying fish | 156640 | 9.6 | 6.0 | 0.63 |
| *Cheilopogon furcatus* | flying fish | 156640 | 10.0 | 6.0 | 0.60 |
| *Cheilopogon furcatus* | flying fish | 156640 | 13.5 | 8.5 | 0.63 |
| *Cheilopogon furcatus* | flying fish | 42535 | 8.7 | 5.5 | 0.63 |
| *Cheilopogon furcatus* | flying fish | 42535 | 4.8 | 3.2 | 0.67 |
| *Cheilopogon furcatus* | flying fish | 42535 | 8.0 | 5.5 | 0.69 |
| *Cheilopogon furcatus* | flying fish | 42535 | 7.3 | 4.5 | 0.62 |
| *Cheilopogon furcatus* | flying fish | 42535 | 6.5 | 4.5 | 0.69 |
| *Cheilopogon furcatus* | flying fish | 156660 | 3.8 | 2.7 | 0.71 |
| *Cheilopogon furcatus* | flying fish | 156660 | 2.0 | 1.4 | 0.70 |
| *Euleptorhamphus velox* | flying halfbeak | 55426 | 24.0 | 5.5 | 0.23 |
| *Euleptorhamphus velox* | flying halfbeak | 55426 | 23.0 | 5.5 | 0.24 |
| *Euleptorhamphus velox* | flying halfbeak | 56810 | 25.0 | 6.5 | 0.26 |
| *Euleptorhamphus velox* | flying halfbeak | 56810 | 24.0 | 6.4 | 0.27 |
| *Euleptorhamphus velox* | flying halfbeak | 56648 | 24.5 | 7.0 | 0.29 |
| *Euleptorhamphus velox* | flying halfbeak | 56648 | 20.5 | 4.3 | 0.21 |
| *Euleptorhamphus velox* | flying halfbeak | 55435 | 6.5 | 1.1 | 0.18 |
| *Euleptorhamphus velox* | flying halfbeak | 55435 | 6.8 | 1.5 | 0.22 |
| *Euleptorhamphus velox* | flying halfbeak | 156025 | 5.8 | 1.2 | 0.21 |
| *Hemiramphus brasiliensis* | halfbeak | 5203 | 19.8 | 3.5 | 0.18 |
| *Hemiramphus brasiliensis* | halfbeak | 5203 | 23.6 | 3.5 | 0.15 |
| *Hemiramphus brasiliensis* | halfbeak | 5203 | 21.5 | 3.0 | 0.14 |
| *Hemiramphus brasiliensis* | halfbeak | 5203 | 20.8 | 3.0 | 0.14 |
| *Hemiramphus brasiliensis* | halfbeak | 5203 | 17.2 | 2.5 | 0.15 |
| *Hemiramphus brasiliensis* | halfbeak | 5203 | 20.5 | 3.2 | 0.16 |
| *Hemiramphus brasiliensis* | halfbeak | 5203 | 19.8 | 3.0 | 0.15 |
| *Hemiramphus brasiliensis* | halfbeak | 55424 | 5.7 | 0.8 | 0.14 |
| *Hemiramphus brasiliensis* | halfbeak | 56635,56636,56637,56638 | 3.1 | 0.4 | 0.13 |
| *Hemiramphus brasiliensis* | halfbeak | 56635,56636,56637,56638 | 2.6 | 0.3 | 0.12 |
| *Oxyporhamphus micropterus* | flying halfbeak | 156057 | 13.3 | 4.7 | 0.35 |
| *Oxyporhamphus micropterus* | flying halfbeak | 156057 | 13.2 | 5.0 | 0.38 |
| *Oxyporhamphus micropterus* | flying halfbeak | 156057 | 14.0 | 5.0 | 0.36 |
| *Oxyporhamphus micropterus* | flying halfbeak | 156057 | 13.4 | 4.5 | 0.34 |
| *Oxyporhamphus micropterus* | flying halfbeak | 41500 | 12.1 | 4.7 | 0.39 |
| *Oxyporhamphus micropterus* | flying halfbeak | 41500 | 12.0 | 4.0 | 0.33 |
| *Oxyporhamphus micropterus* | flying halfbeak | 41500 | 13.0 | 4.4 | 0.34 |
| *Oxyporhamphus micropterus* | flying halfbeak | 149668 | 11.5 | 3.5 | 0.30 |

| | | | | | |
|---|---|---|---|---|---|
| *Oxyporhamphus micropterus* | flying halfbeak | 149668 | 6.2 | 1.7 | 0.27 |
| *Oxyporhamphus micropterus* | flying halfbeak | 149668 | 3.2 | 0.8 | 0.25 |
| *Hyporhamphus unifasciatus* | halfbeak | 28869 | 14.5 | 2.1 | 0.14 |
| *Hyporhamphus unifasciatus* | halfbeak | 28869 | 14.7 | 2.0 | 0.14 |
| *Hyporhamphus unifasciatus* | halfbeak | 28869 | 15.8 | 2.0 | 0.13 |
| *Hyporhamphus unifasciatus* | halfbeak | 28869 | 14.5 | 2.0 | 0.14 |
| *Hyporhamphus unifasciatus* | halfbeak | 28869 | 15.0 | 2.1 | 0.14 |
| *Hyporhamphus unifasciatus* | halfbeak | 44163 | 20.0 | 2.5 | 0.13 |
| *Hyporhamphus unifasciatus* | halfbeak | 57865 | 13.5 | 1.9 | 0.14 |
| *Hyporhamphus unifasciatus* | halfbeak | 57865 | 9.5 | 1.3 | 0.14 |
| *Hyporhamphus unifasciatus* | halfbeak | 57865 | 10.5 | 1.5 | 0.14 |
| *Hyporhamphus unifasciatus* | halfbeak | 57865 | 9.5 | 1.3 | 0.14 |
| *Hyporhamphus unifasciatus* | halfbeak | 57865 | 9.2 | 1.2 | 0.13 |
| *Hyporhamphus unifasciatus* | halfbeak | 57865 | 9.7 | 1.4 | 0.14 |
| *Belone belone* | needlefish | 23443 | 35.0 | 3.0 | 0.09 |
| *Belone belone* | needlefish | 23443 | 53.0 | 4.0 | 0.08 |
| *Belone belone* | needlefish | 655 | 42.0 | 3.5 | 0.08 |
| *Belone belone* | needlefish | 655 | 20.5 | 1.5 | 0.07 |
| *Strongylura marina* | needlefish | 642 | 42.0 | 3.5 | 0.08 |
| *Strongylura marina* | needlefish | 642 | 30.5 | 3.1 | 0.10 |
| *Strongylura marina* | needlefish | 642 | 31.0 | 2.2 | 0.07 |
| *Strongylura marina* | needlefish | 642 | 37.5 | 3.6 | 0.10 |
| *Strongylura marina* | needlefish | 642 | 28.5 | 2.5 | 0.09 |
| *Strongylura marina* | needlefish | 642 | 25.5 | 2.0 | 0.08 |
| *Strongylura marina* | needlefish | 642 | 23.5 | 2.2 | 0.09 |
| *Strongylura marina* | needlefish | 642 | 25.0 | 2.0 | 0.08 |
| *Strongylura marina* | needlefish | 40915 | 13.0 | 1.3 | 0.10 |
| *Strongylura marina* | needlefish | 40915 | 22.5 | 2.0 | 0.09 |
| *Strongylura marina* | needlefish | 40915 | 15.0 | 1.5 | 0.10 |
| *Strongylura marina* | needlefish | 40915 | 18.0 | 1.8 | 0.10 |
| *Strongylura marina* | needlefish | 40915 | 17.0 | 1.6 | 0.09 |
| *Strongylura marina* | needlefish | 40915 | 14.1 | 1.5 | 0.11 |
| *Strongylura marina* | needlefish | 60185 | 17.5 | 1.5 | 0.09 |

#### S2 Table. Pooled populations and targeted sequence capture

| Species | # Individuals Sequenced | CDS capture | CNE capture | Avg nucleotide diversity (π) |
|---|---|---|---|---|
| *Ablennis hians* | 5 | ✓ | ✓ | 0.0025 |
| *Arrhamphus sclerolepis* | 2 | ✓ | ✓ | 0.0030 |
| *Belone belone* | 5 | ✓ | ✓ | 0.0044 |
| *Belonion dibranchodon* | 5 | ✓ | ✓ | 0.0020 |
| *Cheilopogon furcatus* | 7 | ✓ | ✓ | 0.0059 |
| *Cheilopogon papilio* | 7 | ✓ | ✓ | 0.0057 |
| *Cheilopogon xenopterus* | 8 | ✓ | ✓ | 0.0054 |
| *Chriodorus atherinoides* | 8 | ✓ | ✓ | 0.0027 |
| *Cololabis saira* | 1 | ✓ | ✓ | 0.0101 |
| *Cypselurus callopterus* | 8 | ✓ | ✓ | 0.0048 |
| *Euleptorhamphus viridis* | 5 | ✓ | ✓ | 0.0050 |
| *Exocoetus volitans* | 8 | ✓ | ✓ | 0.0030 |
| *Fodiator rostratus* | 5 | ✓ | ✓ | 0.0063 |
| *Hemiramphus brasiliensis* | 5 | ✓ | ✓ | 0.0042 |
| *Hemiramphus far* | 6 | ✓ | ✓ | 0.0053 |
| *Hemiramphus unifasciatus* | 8 | ✓ | ✓ | 0.0027 |
| *Hemirhamphodon pogonognathus* | 5 | ✓ | ✓ | 0.0049 |
| *Hemirhamphodon tengah* | 1 | ✓ | ✓ | 0.0043 |
| *Hirundichthys rondeleti* | 5 | ✓ | | 0.0055 |
| *Hyporhamphus brederi* | 6 | ✓ | ✓ | 0.0036 |
| *Hyporhamphus quoyi* | 5 | ✓ | ✓ | 0.0031 |
| *Melapedalion breve* | 2 | ✓ | ✓ | 0.0024 |
| *Oxyporhamphus micropterus* | 8 | ✓ | ✓ | 0.0044 |
| *Parexocoetus brachypterus* | 6 | ✓ | ✓ | 0.0057 |
| *Potamorrhamphis guianensis* | 8 | ✓ | ✓ | 0.0032 |
| *Prognichthys tringa* | 7 | ✓ | ✓ | 0.0053 |
| *Pseudotylosurus angusticeps* | 6 | ✓ | ✓ | 0.0014 |
| *Rhynchorhamphus georgii* | 2 | ✓ | ✓ | 0.0030 |
| *Strongylura fluviatilis* | 5 | ✓ | ✓ | 0.0017 |
| *Strongylura hubbsi* | 3 | ✓ | | 0.0075 |
| *Strongylura marina* | 5 | ✓ | ✓ | 0.0034 |
| *Strongylura notata* | 4 | ✓ | ✓ | 0.0024 |
| *Tylosurus crocodilus* | 5 | ✓ | ✓ | 0.0041 |
| *Xenentodon cancila* | 5 | ✓ | ✓ | 0.0059 |
| *Zenarchopterus spp* | 2 | ✓ | ✓ | 0.0099 |

### S3 Table. Beloniform sample collection

| Species | Lovejoy Fish # | Museum catalog number | Waterbody |
|---|---|---|---|
| Ablennes hians | 1024 | | |
| Ablennes hians | 1025 | | |
| Ablennes hians | 1026 | | |
| Ablennes hians | 915 | | |
| Ablennes hians | 916 | | |
| Arrhamphus sclerolepis | 2691 | | |
| Arrhamphus sclerolepis | 2692 | | |
| Belone belone | 922 | | Trieste, Adriatic, Italy |
| Belone belone | 1002 | | Courtmacsherry Bay, Co. Cork, Ireland |
| Belone belone | 1003 | | Courtmacsherry Bay, Co. Cork, Ireland |
| Belone belone | 1004 | | Courtmacsherry Bay, Co. Cork, Ireland |
| Belone belone | 1417 | | Haifo, Mediterranean, Israel |
| Belonion dibranchodon | 1504 | | Rio Atabapo, Orinoco, Venezuela |
| Belonion dibranchodon | 1505 | | Rio Atabapo, Orinoco, Venezuela |
| Belonion dibranchodon | 1507 | | Rio Atabapo, Orinoco, Venezuela |
| Belonion dibranchodon | 1508 | | Rio Atabapo, Orinoco, Venezuela |
| Belonion dibranchodon | 1509 | | Rio Atabapo, Orinoco, Venezuela |
| Cheilopogon furcatus | 3696 | ROM-90530 | South Pacific Ocean |
| Cheilopogon furcatus | 3317 | ROM-79317 | Gulf of Mexico |
| Cheilopogon furcatus | 3363 | ROM-79259 | Gulf of Mexico |
| Cheilopogon furcatus | 3176 | ROM-90668 | Gulf of Mexico |
| Cheilopogon furcatus | 3177 | ROM-90668 | Gulf of Mexico |
| Cheilopogon furcatus | 3178 | ROM-90668 | Gulf of Mexico |
| Cheilopogon furcatus | 3313 | ROM-91967 | Gulf of Mexico |
| Cheilopogon papilio | 5076 | ROM-92617 | North Pacific Ocean |
| Cheilopogon papilio | 6246 | ROM-92911 | North Pacific Ocean |
| Cheilopogon papilio | 4752 | ROM-92599 | North Pacific Ocean |
| Cheilopogon papilio | 5351 | ROM-92641 | North Pacific Ocean |
| Cheilopogon papilio | 4382 | ROM-92570 | Eastern Tropical Pacific |
| Cheilopogon papilio | 4408 | ROM-92570 | Eastern Tropical Pacific |
| Cheilopogon papilio | 5354 | ROM-92642 | North Pacific Ocean |
| Cheilopogon papilio | 5355 | ROM-92642 | North Pacific Ocean |
| Cheilopogon xenopterus | 4073 | ROM-92546 | North Pacific Ocean |
| Cheilopogon xenopterus | 4076 | ROM-92546 | North Pacific Ocean |
| Cheilopogon xenopterus | 4078 | ROM-92546 | North Pacific Ocean |

| | | | |
|---|---|---|---|
| Cheilopogon xenopterus | 4126 | ROM-92551 | North Pacific Ocean |
| Cheilopogon xenopterus | 4128 | ROM-92551 | North Pacific Ocean |
| Cheilopogon xenopterus | 5057 | ROM-92616 | North Pacific Ocean |
| Cheilopogon xenopterus | 5113 | ROM-92620 | North Pacific Ocean |
| Cheilopogon xenopterus | 5114 | ROM-92620 | North Pacific Ocean |
| Chriodorus atherinoides | 1135 | | Florida Bay |
| Chriodorus atherinoides | 1136 | | Florida Bay |
| Chriodorus atherinoides | 1137 | | Florida Bay |
| Chriodorus atherinoides | 1138 | | Florida Bay |
| Chriodorus atherinoides | 1139 | | Florida Bay |
| Chriodorus atherinoides | 1140 | | Florida Bay |
| Chriodorus atherinoides | 1141 | | Florida Bay |
| Chriodorus atherinoides | 1142 | | Florida Bay |
| Cololabis saira | 1646 | | Eastern Pacific |
| Cypselurus callopterus | 3984 | ROM-92541 | North Pacific Ocean |
| Cypselurus callopterus | 4149 | ROM-92552 | North Pacific Ocean |
| Cypselurus callopterus | 4151 | ROM-92552 | North Pacific Ocean |
| Cypselurus callopterus | 6221 | ROM-92902 | North Pacific Ocean |
| Cypselurus callopterus | 6361 | ROM-92932 | North Pacific Ocean |
| Cypselurus callopterus | 5396 | ROM-92648 | North Pacific Ocean |
| Cypselurus callopterus | 5397 | ROM-92648 | North Pacific Ocean |
| Cypselurus callopterus | 5016 | ROM-92610 | North Pacific Ocean |
| Euleptorhamphus viridis | 3912 | | |
| Euleptorhamphus viridis | 3913 | | |
| Euleptorhamphus viridis | 4375 | | |
| Euleptorhamphus viridis | 4745 | | |
| Euleptorhamphus viridis | 4746 | | |
| Euleptorhamphus viridis | 5298 | | |
| Exocoetus volitans | 5065 | ROM-79310 | North Pacific Ocean |
| Exocoetus volitans | 5185 | ROM-79271 | North Pacific Ocean |
| Exocoetus volitans | 5195 | ROM-79219 | North Pacific Ocean |
| Exocoetus volitans | 5686 | ROM-79293 | North Pacific Ocean |
| Exocoetus volitans | 5693 | ROM-79220 | North Pacific Ocean |
| Exocoetus volitans | 5706 | ROM-79222 | North Pacific Ocean |
| Exocoetus volitans | 5721 | ROM-79284 | North Pacific Ocean |
| Exocoetus volitans | 5842 | ROM-79309 | North Pacific Ocean |
| Fodiator rostratus | 3228 | ROM-91960 | Eastern Tropical Pacific |
| Fodiator rostratus | 5383 | ROM-92647 | North Pacific Ocean |
| Fodiator rostratus | 5384 | ROM-92647 | North Pacific Ocean |
| Fodiator rostratus | 5447 | ROM-92654 | North Pacific Ocean |

| | | | |
|---|---|---|---|
| Fodiator rostratus | 5762 | ROM-92694 | North Pacific Ocean |
| Fodiator rostratus | 5884 | ROM-92705 | North Pacific Ocean |
| Hemiramphus brasiliensis | 1054 | | |
| Hemiramphus brasiliensis | 1055 | | |
| Hemiramphus brasiliensis | 1056 | | |
| Hemiramphus brasiliensis | 1057 | | |
| Hemiramphus brasiliensis | 1058 | | |
| Hemiramphus far | 1145 | | |
| Hemiramphus far | 1146 | | |
| Hemiramphus far | 1418 | | |
| Hemiramphus far | 1420 | | |
| Hemiramphus far | 1421 | | |
| Hemiramphus far | 7124 | | |
| Hemiramphus unifasciatus | 1275 | | |
| Hemiramphus unifasciatus | 1276 | | |
| Hemiramphus unifasciatus | 1277 | | |
| Hemiramphus unifasciatus | 1278 | | |
| Hemiramphus unifasciatus | 1279 | | |
| Hemiramphus unifasciatus | 1280 | | |
| Hemiramphus unifasciatus | 1281 | | |
| Hemiramphus unifasciatus | 1282 | | |
| Hemirhamphodon pogonognathus | 1235 | | Singapore |
| Hemirhamphodon pogonognathus | 1236 | | Singapore |
| Hemirhamphodon pogonognathus | N7125 | | |
| Hemirhamphodon pogonognathus | N7126 | | |
| Hemirhamphodon pogonognathus | N7127 | | |
| Hemirhamphodon tengah | 7129 | | Kalimantan Tengah |
| Hirundichthys rondeleti | 6923 | ROM-90537 | North Pacific Ocean |
| Hirundichthys rondeleti | 3324 | ROM-79273 | Gulf of Mexico |
| Hirundichthys rondeleti | 3360 | ROM-79265 | Gulf of Mexico |
| Hirundichthys rondeleti | 3361 | ROM-79265 | Gulf of Mexico |
| Hirundichthys rondeleti | 3510 | ROM-79290 | Gulf of Mexico |
| Hirundichthys rondeleti | 3454 | ROM-79324 | Gulf of Mexico |
| Hirundichthys rondeleti | 3512 | ROM-92493 | Gulf of Mexico |
| Hirundichthys rondeleti | 3394 | ROM-91988 | Gulf of Mexico |
| Hyporhamphus brederi | 10872 | | |
| Hyporhamphus brederi | 10873 | | |
| Hyporhamphus brederi | 10874 | | |
| Hyporhamphus brederi | 10875 | | |
| Hyporhamphus brederi | 10876 | | |

| | | | |
|---|---|---|---|
| Hyporhamphus brederi | 10877 | | |
| Hyporhamphus quoyi | 7167 | | Senoko Fishery Port, Singapore |
| Hyporhamphus quoyi | 7168 | | Senoko Fishery Port, Singapore |
| Hyporhamphus quoyi | 7169 | | Senoko Fishery Port, Singapore |
| Hyporhamphus quoyi | 7170 | | Senoko Fishery Port, Singapore |
| Hyporhamphus quoyi | 7171 | | Senoko Fishery Port, Singapore |
| Melapedalion breve | 7179 | | Senoko Fishery Port, Singapore |
| Melapedalion breve | 7180 | | Senoko Fishery Port, Singapore |
| Oxyporhamphus micropterus | 2796 | | |
| Oxyporhamphus micropterus | 2797 | | |
| Oxyporhamphus micropterus | 2798 | | |
| Oxyporhamphus micropterus | 3917 | | |
| Oxyporhamphus micropterus | 3918 | | |
| Oxyporhamphus micropterus | 3919 | | |
| Oxyporhamphus micropterus | 1589 | | |
| Oxyporhamphus micropterus | 1590 | | |
| Parexocoetus brachypterus | 3306 | ROM-93019 | |
| Parexocoetus brachypterus | 3307 | ROM-93019 | |
| Parexocoetus brachypterus | 3308 | ROM-93019 | |
| Parexocoetus brachypterus | 3309 | ROM-93019 | |
| Parexocoetus brachypterus | 3310 | ROM-93019 | |
| Potamorrhaphis guianensis | 10339 | | |
| Potamorrhaphis guianensis | 10340 | | |
| Potamorrhaphis guianensis | 10341 | | |
| Potamorrhaphis guianensis | 10342 | | |
| Potamorrhaphis guianensis | 934 | | |
| Potamorrhaphis guianensis | 935 | | |
| Potamorrhaphis guianensis | 937 | | |
| Prognichthys tringa | 4154 | ROM-92553 | North Pacific Ocean |
| Prognichthys tringa | 4743 | ROM-92596 | North Pacific Ocean |
| Prognichthys tringa | 4764 | ROM-92596 | North Pacific Ocean |
| Prognichthys tringa | 4765 | ROM-92596 | North Pacific Ocean |
| Prognichthys tringa | 5205 | ROM-92628 | North Pacific Ocean |
| Prognichthys tringa | 5466 | ROM-92657 | North Pacific Ocean |
| Prognichthys tringa | 5477 | ROM-92657 | North Pacific Ocean |
| Prognichthys tringa | 5493 | ROM-92658 | North Pacific Ocean |
| Pseudotylosurus angusticeps | 1007 | | Rio Napo, Ecuador |
| Pseudotylosurus angusticeps | 1008 | | Rio Napo, Ecuador |
| Pseudotylosurus angusticeps | 1010 | | Rio Napo, Ecuador |
| Pseudotylosurus angusticeps | 1011 | | Rio Napo, Ecuador |

| | | |
|---|---|---|
| Pseudotylosurus angusticeps | 1013 | Rio Napo, Ecuador |
| Pseudotylosurus angusticeps | 1014 | Rio Napo, Ecuador |
| Rhyncorhamphus georgii | 9889 | |
| Rhyncorhamphus georgii | 9896 | |
| Strongylura fluviatilis | 1015 | Rio Cayapas, Ecuador |
| Strongylura fluviatilis | 1016 | Rio Cayapas, Ecuador |
| Strongylura fluviatilis | 1017 | Rio Cayapas, Ecuador |
| Strongylura fluviatilis | 1018 | Rio Cayapas, Ecuador |
| Strongylura fluviatilis | 1019 | Rio Cayapas, Ecuador |
| Strongylura hubbsi | 1068 | Rio Usumacinta, Guatemala |
| Strongylura hubbsi | 1069 | Rio Usumacinta, Guatemala |
| Strongylura hubbsi | 1070 | Rio Usumacinta, Guatemala |
| Strongylura hubbsi | 1071 | Rio Usumacinta, Guatemala |
| Strongylura marina | 1319 | |
| Strongylura marina | 1320 | |
| Strongylura marina | 1321 | |
| Strongylura marina | 1322 | |
| Strongylura marina | 1323 | |
| Strongylura notata | 1081 | |
| Strongylura notata | 1082 | |
| Strongylura notata | 1083 | |
| Strongylura notata | 1084 | |
| Tylosurus crocodilus | 914 | |
| Tylosurus crocodilus | 1051 | |
| Tylosurus crocodilus | 1092 | |
| Tylosurus crocodilus | 1093 | |
| Tylosurus crocodilus | 1326 | |
| Xenentodon cancila | 1253 | |
| Xenentodon cancila | 1254 | |
| Xenentodon cancila | 1255 | |
| Xenentodon cancila | 1256 | |
| Xenentodon cancila | 1257 | |
| Xenentodon cancila | 1258 | |
| Zenarchopterus sp. | 1207 | Bunaken, Sulawesi |
| Zenarchopterus sp. | 1208 | Bunaken, Sulawesi |

#### S4 Table. Coverage breakdown

| | Reference genome of target | | | | | | | | | | | | | |
|---|---|---|---|---|---|---|---|---|---|---|---|---|---|---|
| | Medaka | | Platyfish | | Amazon Molly | | CDS* | | CNE*† | | miRNA* | | UCNE* | |
| Species | Cov. | Depth | Cov. | Depth | Cov. | Depth | Cov. | Depth | Cov. | Depth | Cov. | Depth | Cov. | Depth |
| *Xenentodon cancila* | 80.5% | 40.2 | 63.8% | 24.5 | 41.8% | 49.2 | 81.3% | 35.3 | 78.1% | 48.9 | 91.1% | 51.1 | 97.5% | 78.6 |
| *Ablennis hians* | 80.4% | 34.2 | 64.6% | 22.2 | 44.0% | 26.8 | 81.1% | 32.3 | 78.5% | 37.7 | 89.3% | 36.5 | 97.3% | 51.3 |
| *Tylosurus crocodilus* | 75.3% | 43.8 | 57.9% | 23.7 | 40.8% | 44.8 | 73.2% | 32.8 | 78.8% | 65.2 | 89.0% | 52.3 | 96.5% | 78.3 |
| *Strongylura notata* | 81.9% | 46.7 | 66.8% | 32.9 | 45.9% | 47.0 | 83.5% | 48.1 | 78.1% | 43.1 | 89.2% | 53.1 | 97.7% | 70.3 |
| *Belone belone* | 81.4% | 43.7 | 66.3% | 34.3 | 46.6% | 53.1 | 83.2% | 48.2 | 77.2% | 34.4 | 89.6% | 41.9 | 96.4% | 49.9 |
| *Cololabis saira* | 80.2% | 41.3 | 63.3% | 29.1 | 44.5% | 54.0 | 81.3% | 43.4 | 77.4% | 36.4 | 89.8% | 39.9 | 96.5% | 58.1 |
| *Pseudotylosurus angusticeps* | 81.8% | 38.5 | 63.1% | 24.9 | 32.9% | 22.1 | 81.8% | 38.5 | - | - | - | - | - | - |
| *Strongylura marina* | 76.4% | 37.4 | 59.0% | 24.8 | 39.2% | 31.0 | 75.7% | 35.8 | 77.1% | 40.0 | 88.2% | 37.4 | 96.6% | 53.5 |
| *Strongylura hubbsi* | 80.4% | 28.7 | 62.4% | 18.3 | 35.2% | 21.6 | 80.4% | 28.7 | - | - | - | - | - | - |
| *Strongylura fluviatilis* | 81.1% | 40.2 | 65.0% | 24.4 | 44.2% | 26.7 | 82.2% | 33.8 | 78.2% | 51.5 | 89.3% | 57.4 | 97.7% | 88.8 |
| *Potamorrhamphis guianensis* | 78.7% | 33.0 | 61.2% | 20.0 | 40.4% | 36.6 | 78.8% | 27.2 | 77.7% | 43.8 | 90.0% | 46.1 | 97.5% | 71.1 |
| *Belonion dibranchodon* | 76.6% | 23.6 | 59.4% | 17.7 | 36.0% | 17.1 | 78.6% | 27.1 | 71.8% | 16.1 | 85.0% | 21.7 | 95.4% | 29.2 |
| *Zenarchopterus spp* | 82.6% | 59.2 | 67.9% | 38.6 | 46.9% | 46.8 | 84.5% | 55.7 | 78.1% | 64.6 | 90.5% | 63.6 | 97.4% | 108.0 |
| *Hemirhamphodon pogonognathus* | 80.0% | 49.5 | 63.9% | 35.9 | 39.7% | 31.3 | 82.7% | 56.1 | 73.7% | 35.4 | 87.3% | 41.4 | 95.9% | 66.2 |
| *Hemirhamphodon tengah* | 79.4% | 43.2 | 62.8% | 29.6 | 37.6% | 26.9 | 81.7% | 45.4 | 73.9% | 37.6 | 88.3% | 41.6 | 95.6% | 71.9 |
| *Melapedalion breve* | 81.6% | 39.4 | 67.0% | 32.2 | 46.5% | 38.8 | 83.7% | 46.2 | 76.7% | 25.5 | 88.2% | 30.2 | 96.2% | 39.3 |
| *Hyporhamphus quoyi* | 82.0% | 43.8 | 67.7% | 36.1 | 47.9% | 53.1 | 84.2% | 51.5 | 77.0% | 27.9 | 88.3% | 36.5 | 96.6% | 43.2 |
| *Arrhamphus sclerolepis* | 78.1% | 29.1 | 63.0% | 23.6 | 43.5% | 33.1 | 80.3% | 36.3 | 73.1% | 14.4 | 83.2% | 21.3 | 94.2% | 22.9 |
| *Chriodorus atherinoides* | 81.6% | 51.6 | 65.8% | 32.6 | 46.1% | 44.9 | 82.9% | 45.3 | 78.2% | 63.0 | 89.2% | 67.2 | 97.0% | 94.6 |
| *Hemiramphus unifasciatus* | 71.6% | 25.5 | 52.7% | 16.6 | 36.8% | 19.0 | 69.1% | 23.5 | 75.7% | 28.7 | 85.8% | 32.1 | 95.8% | 46.3 |
| *Hyporhamphus brederi* | 72.4% | 31.3 | 54.0% | 16.7 | 39.3% | 28.9 | 69.4% | 20.7 | 77.8% | 51.1 | 88.7% | 53.6 | 96.8% | 89.3 |
| *Euleptorhamphus viridis* | 75.1% | 35.4 | 57.6% | 21.3 | 40.7% | 35.6 | 73.6% | 31.2 | 77.5% | 43.0 | 88.9% | 39.6 | 97.0% | 61.2 |
| *Rhynchorhamphus georgii* | 81.6% | 47.3 | 66.2% | 26.5 | 45.3% | 31.6 | 82.6% | 37.4 | 79.1% | 66.7 | 89.5% | 53.0 | 97.6% | 71.4 |
| *Hemiramphus far* | 80.6% | 35.9 | 64.1% | 20.8 | 45.6% | 34.2 | 80.4% | 29.0 | 80.2% | 49.0 | 89.3% | 40.1 | 97.6% | 63.5 |
| *Hemiramphus brasiliensis* | 72.6% | 25.4 | 56.1% | 19.5 | 39.3% | 40.5 | 72.7% | 30.5 | 71.5% | 14.8 | 82.4% | 18.3 | 95.3% | 25.2 |
| *Oxyporhamphus micropterus* | 77.0% | 32.6 | 59.1% | 18.2 | 41.0% | 32.5 | 76.1% | 25.3 | 78.3% | 47.0 | 89.1% | 40.7 | 95.9% | 56.1 |
| *Parexocoetus brachypterus* | 78.1% | 36.4 | 61.3% | 20.8 | 43.6% | 32.4 | 78.4% | 29.4 | 76.8% | 50.0 | 89.4% | 42.5 | 96.3% | 59.5 |
| *Fodiator rostratus* | 72.1% | 37.0 | 53.6% | 20.7 | 37.0% | 26.2 | 68.7% | 29.8 | 78.1% | 50.6 | 89.0% | 49.2 | 96.7% | 72.6 |
| *Exocoetus volitans* | 67.6% | 25.3 | 48.8% | 15.1 | 34.5% | 33.8 | 63.5% | 22.0 | 74.9% | 31.3 | 86.3% | 28.6 | 95.5% | 40.9 |
| *Cheilopogon papilio* | 80.3% | 38.5 | 63.6% | 24.2 | 43.9% | 29.1 | 80.7% | 33.8 | 78.8% | 46.9 | 89.9% | 48.7 | 97.2% | 72.1 |
| *Cypselurus callopterus* | 78.9% | 44.3 | 61.4% | 25.4 | 42.5% | 32.8 | 78.5% | 35.5 | 79.0% | 60.6 | 89.2% | 58.8 | 97.3% | 92.1 |
| *Prognichthys tringa* | 72.5% | 36.3 | 54.2% | 20.9 | 37.9% | 28.7 | 69.8% | 30.2 | 77.2% | 47.5 | 88.5% | 44.0 | 97.0% | 75.2 |
| *Cheilopogon furcatus* | 79.5% | 35.0 | 62.7% | 22.4 | 44.0% | 31.6 | 79.8% | 31.4 | 78.2% | 41.3 | 89.0% | 39.3 | 97.2% | 60.2 |
| *Hirundichthys rondeleti* | 78.4% | 45.2 | 59.9% | 28.7 | 37.0% | 62.9 | 78.4% | 45.2 | - | - | - | - | - | - |
| *Cheilopogon xenopterus* | 80.6% | 41.0 | 64.4% | 26.0 | 43.8% | 32.7 | 81.2% | 36.6 | 78.8% | 48.9 | 89.0% | 46.3 | 97.3% | 71.7 |

* - medaka (*Oryzias latipes*) reference genome targets

† - not inclusive of miRNAs and ultraconservative elements (UCNE)

#### S5 Table. Sequencing read coverage by depth

| | coverage† | | | |
|---|---|---|---|---|
| | 1X | 2X | 4X | 10X |
| *Xenentodon cancila* | 80.5% | 80.1% | 78.0% | 67.9% |
| *Ablennis hians* | 80.4% | 79.9% | 77.4% | 65.8% |
| *Tylosurus crocodilus* | 75.3% | 74.3% | 70.5% | 59.5% |
| *Strongylura notata* | 81.9% | 81.5% | 79.9% | 72.3% |
| *Belone belone* | 81.4% | 81.1% | 79.4% | 72.1% |
| *Cololabis saira* | 80.2% | 79.8% | 78.0% | 69.7% |
| *Pseudotylosurus angusticeps* | 81.8% | 81.4% | 79.3% | 69.4% |
| *Strongylura marina* | 76.4% | 75.6% | 72.1% | 61.4% |
| *Strongylura hubbsi* | 80.4% | 79.8% | 76.7% | 64.5% |
| *Strongylura fluviatilis* | 81.1% | 80.6% | 78.4% | 69.0% |
| *Potamorrhamphis guianensis* | 78.7% | 78.1% | 75.2% | 62.9% |
| *Belonion dibranchodon* | 76.6% | 75.9% | 72.4% | 58.5% |
| *Zenarchopterus spp* | 82.6% | 82.4% | 81.4% | 75.8% |
| *Hemirhamphodon pogonognathus* | 80.0% | 79.7% | 78.5% | 72.3% |
| *Hemirhamphodon tengah* | 79.4% | 79.0% | 77.4% | 69.6% |
| *Melapedalion breve* | 81.6% | 81.2% | 79.5% | 70.9% |
| *Hyporhamphus quoyi* | 82.0% | 81.7% | 80.1% | 72.5% |
| *Arrhamphus sclerolepis* | 78.1% | 77.5% | 74.4% | 60.8% |
| *Chriodorus atherinoides* | 81.6% | 81.2% | 79.4% | 71.8% |
| *Hemiramphus unifasciatus* | 71.6% | 70.4% | 65.7% | 53.0% |
| *Hyporhamphus brederi* | 72.4% | 71.3% | 66.8% | 54.5% |
| *Euleptorhamphus viridis* | 75.1% | 74.1% | 70.1% | 58.3% |
| *Rhynchorhamphus georgii* | 81.6% | 81.2% | 79.3% | 69.5% |
| *Hemiramphus far* | 80.6% | 80.0% | 77.4% | 65.9% |
| *Hemiramphus brasiliensis* | 72.6% | 71.4% | 66.5% | 51.9% |
| *Oxyporhamphus micropterus* | 77.0% | 76.4% | 73.7% | 60.5% |
| *Parexocoetus brachypterus* | 78.1% | 77.6% | 75.0% | 63.0% |
| *Fodiator rostratus* | 72.1% | 71.0% | 66.7% | 54.6% |
| *Exocoetus volitans* | 67.6% | 66.2% | 60.8% | 47.6% |
| *Cheilopogon papilio* | 80.3% | 79.8% | 77.6% | 67.1% |
| *Cypselurus callopterus* | 78.9% | 78.3% | 75.7% | 64.5% |
| *Prognichthys tringa* | 72.5% | 71.4% | 67.0% | 55.7% |
| *Cheilopogon furcatus* | 79.5% | 79.0% | 76.4% | 65.1% |
| *Hirundichthys rondeleti* | 78.4% | 77.6% | 74.4% | 63.9% |
| *Cheilopogon xenopterus* | 80.6% | 80.2% | 78.1% | 68.1% |

#### S7 Table. Recovery of heterozygosity and species-specific SNPs

| Species | # Het SNPs | Het SNP/ base | Percent of targets with het SNP | # Fixed species-specific SNPs | Fixed SNPs/ base | Percent of targets with fixed, species-specific SNP |
|---|---|---|---|---|---|---|
| *Xenentodon cancila* | 31,905 | 0.0020 | 36.9% | 281,278 | 0.0075 | 49.2% |
| *Ablennis hians* | 27,735 | 0.0011 | 14.0% | 87,054 | 0.0024 | 19.9% |
| *Tylosurus crocodilus* | 32,914 | 0.0016 | 17.2% | 46,321 | 0.0014 | 11.7% |
| *Belone belone* | 38,257 | 0.0016 | 20.4% | 274,616 | 0.0070 | 44.7% |
| *Cololabis saira* | 44,872 | 0.0025 | 34.5% | 471,359 | 0.0123 | 58.4% |
| *Strongylura notata* | 27,012 | 0.0009 | 12.4% | 204,406 | 0.0052 | 39.7% |
| *Pseudotylosurus angusticeps* | 13,415 | 0.0006 | 4.0% | 261,936 | 0.0089 | 54.0% |
| *Strongylura fluviatilis* | 25,742 | 0.0008 | 3.1% | 123,477 | 0.0032 | 28.2% |
| *Strongylura hubbsi* | 28,115 | 0.0017 | 11.2% | 74,543 | 0.0026 | 21.9% |
| *Strongylura marina* | 27,161 | 0.0011 | 13.9% | 74,334 | 0.0021 | 19.1% |
| *Potamorrhamphis guianensis* | 28,427 | 0.0013 | 18.5% | 188,759 | 0.0053 | 39.5% |
| *Belonion dibranchodon* | 14,170 | 0.0006 | 10.4% | 280,410 | 0.0082 | 53.2% |
| *Zenarchopterus spp* | 41,630 | 0.0032 | 50.4% | 436,017 | 0.0108 | 58.4% |
| *Hemirhamphodon tengah* | 17,385 | 0.0006 | 13.9% | 288,967 | 0.0077 | 47.3% |
| *Hemirhamphodon pogonognathus* | 21,463 | 0.0011 | 33.8% | 321,368 | 0.0084 | 48.9% |
| *Hyporhamphus quoyi* | 31,867 | 0.0011 | 12.4% | 82,190 | 0.0021 | 16.3% |
| *Melapedalion breve* | 27,593 | 0.0009 | 5.5% | 87,424 | 0.0023 | 18.4% |
| *Arrhamphus sclerolepis* | 24,017 | 0.0009 | 10.4% | 202,073 | 0.0058 | 41.2% |
| *Hemiramphus unifasciatus* | 22,973 | 0.0010 | 12.6% | 199,931 | 0.0064 | 41.5% |
| *Hyporhamphus brederi* | 29,840 | 0.0015 | 16.3% | 178,365 | 0.0056 | 38.3% |
| *Chriodorus atherinoides* | 29,571 | 0.0010 | 10.2% | 239,728 | 0.0062 | 42.2% |
| *Euleptorhamphus viridis* | 23,271 | 0.0013 | 25.9% | 92,226 | 0.0028 | 24.2% |
| *Rhynchorhamphus georgii* | 25,121 | 0.0010 | 15.1% | 165,714 | 0.0044 | 34.2% |
| *Hemiramphus far* | 36,001 | 0.0017 | 24.9% | 58,759 | 0.0016 | 11.8% |
| *Hemiramphus brasiliensis* | 32,914 | 0.0016 | 17.2% | 46,321 | 0.0014 | 11.7% |
| *Oxyporhamphus micropterus* | 37,391 | 0.0018 | 19.8% | 138,340 | 0.0042 | 27.1% |
| *Parexocoetus brachypterus* | 38,084 | 0.0022 | 31.3% | 250,948 | 0.0072 | 44.1% |
| *Fodiator rostratus* | 31,132 | 0.0022 | 33.8% | 109,653 | 0.0036 | 28.7% |
| *Exocoetus volitans* | 25,278 | 0.0013 | 14.1% | 91,222 | 0.0032 | 26.5% |
| *Cheilopogon papilio* | 38,315 | 0.0020 | 29.3% | 70,793 | 0.0019 | 16.1% |
| *Hirundichthys rondeleti* | 23,574 | 0.0019 | 27.4% | 47,339 | 0.0018 | 16.1% |
| *Prognichthys tringa* | 28,037 | 0.0015 | 23.0% | 44,700 | 0.0014 | 12.5% |
| *Cypselurus callopterus* | 31,566 | 0.0016 | 26.4% | 40,919 | 0.0012 | 9.5% |
| *Cheilopogon xenopterus* | 32,893 | 0.0017 | 29.0% | 41,310 | 0.0011 | 9.3% |
| *Cheilopogon furcatus* | 32,595 | 0.0018 | 30.6% | 37,882 | 0.0010 | 8.7% |

#### S8 Table. Prediction of gene duplication

| species | avg. exon copy # | # predicted duplicate genes† |
|---|---|---|
| *Xenentodon cancila* | 1.026 | 2,989 |
| *Ablennis hians* | 1.024 | 2,822 |
| *Tylosurus crocodilus* | 1.063 | 7,427 |
| *Strongylura notata* | 1.023 | 2,770 |
| *Belone belone* | 1.026 | 2,961 |
| *Cololabis saira* | 1.033 | 3,643 |
| *Pseudotylosurus angusticeps* | 1.023 | 1,955 |
| *Strongylura marina* | 1.039 | 4,512 |
| *Strongylura hubbsi* | 1.066 | 4,227 |
| *Strongylura fluviatilis* | 1.022 | 2,581 |
| *Potamorrhamphis guianensis* | 1.022 | 2,568 |
| *Belonion dibranchodon* | 1.019 | 2,049 |
| *Zenarchopterus spp* | 1.030 | 3,470 |
| *Hemirhamphodon pogonognathus* | 1.021 | 2,308 |
| *Hemirhamphodon tengah* | 1.020 | 2,226 |
| *Melapedalion breve* | 1.026 | 3,068 |
| *Hyporhamphus quoyi* | 1.027 | 3,110 |
| *Arrhamphus sclerolepis* | 1.025 | 2,793 |
| *Chriodorus atherinoides* | 1.026 | 3,030 |
| *Hemiramphus unifasciatus* | 1.042 | 4,738 |
| *Hyporhamphus brederi* | 1.043 | 4,804 |
| *Euleptorhamphus viridis* | 1.024 | 2,753 |
| *Rhynchorhamphus georgii* | 1.023 | 2,736 |
| *Hemiramphus far* | 1.027 | 3,202 |
| *Hemiramphus brasiliensis* | 1.060 | 6,633 |
| *Oxyporhamphus micropterus* | 1.029 | 3,330 |
| *Parexocoetus brachypterus* | 1.030 | 3,342 |
| *Fodiator rostratus* | 1.025 | 2,853 |
| *Exocoetus volitans* | 1.021 | 2,325 |
| *Cheilopogon papilio* | 1.027 | 3,191 |
| *Cypselurus callopterus* | 1.024 | 2,835 |
| *Prognichthys tringa* | 1.023 | 2,592 |
| *Cheilopogon furcatus* | 1.025 | 2,953 |
| *Hirundichthys rondeleti* | 1.033 | 1,815 |
| *Cheilopogon xenopterus* | 1.026 | 3,044 |

† defined as genes where ≥ half of all exons for the gene are represented by ≥2 contigs

### S10 Table. Convergent amino acid substitutions in gliding beloniforms

| Ensembl ID | gene name | SNP*† | *E.viridis* | *O. micropterus* | Exocoetidae** | fin screens‡ | fin/limb GO‡ |
|---|---|---|---|---|---|---|---|
| ENSORLG00000009715 | lat4a | F326L | ✓ | ✓ | ✓ | ✓ | |
| ENSORLG00000017925 | gja5a | G329V | ✓ | ✓ | ✓ | | ✓ |
| ENSORLG00000011281 | fat2 | H2745M | ✓ | ✓ | ✓ | | |
| ENSORLG00000006187 | vit-6 | L321H | ✓ | ✓ | ✓ | | |
| ENSORLG00000014112 | dph5 | R24K | ✓ | ✓ | ✓ | | |
| ENSORLG00000009770 | ptprt | S671G | ✓ | ✓ | ✓ | | |
| ENSORLG00000004903 | ltn1 | K1534R | ✓ | ✓ | ✓ | | |
| ENSORLG00000020892 | capn2b | M571V | ✓ | ✓ | ✓ | | |
| ENSXMAG00000017216 | NA | I31V | ✓ | ✓ | ✓ | | |
| ENSORLG00000017466 | tmem74b | K141R | ✓ | ✓ | ✓ | | |
| ENSORLG00000001003 | smc1b | L2017G | ✓ | ✓ | ✓ | | |
| ENSORLG00000008243 | lipca | L225M | ✓ | ✓ | ✓ | | |
| ENSORLG00000020791 | abcg1 | E496P | ✓ | ✓ | ✓ | | |
| ENSORLG00000012658 | exoc2 | D109P | ✓ | ✓ | ✓ | | |
| ENSXMAG00000013054 | pcdh8l | L222M | ✓ | ✓ | ✓ | | |
| ENSORLG00000009277 | sec31b | L24T | ✓ | ✓ | ✓ | | |
| ENSPFOG00000024240 | ralb | L283G | ✓ | ✓ | ✓ | | |
| ENSORLG00000017081 | pak6a | P397C | ✓ | ✓ | ✓ | | |
| ENSORLG00000001355 | ralb | T138S | ✓ | ✓ | ✓ | | |
| ENSORLG00000009749 | iqgap2 | K103R | ✓ | ✓ | ✓ | | |
| ENSORLG00000002875 | abce1 | E542M | ✓ | ✓ | ✓ | | |
| ENSXMAG00000013372 | si:ch211-113g11.6 | H435R | ✓ | ✓ | ✓ | | |
| ENSORLG00000017258 | arhgef10 | M645I | ✓ | ✓ | ✓ | | |
| ENSORLG00000003234 | fcho2 | H383V | ✓ | ✓ | ✓ | | |
| ENSORLG00000017825 | qrsl1 | K390R | ✓ | ✓ | ✓ | | |
| ENSORLG00000015171 | slc16a10 | E104R | ✓ | ✓ | ✓ | | |
| ENSORLG00000007714 | slc9a1b | E451G | ✓ | ✓ | ✓ | | |
| ENSORLG00000015105 | ocstamp | V109I | ✓ | ✓ | ✓ | | |
| ENSORLG00000013633 | cdh6 | I357V | ✓ | ✓ | ✓ | | |
| ENSXMAG00000009950 | doc2d | P44P | ✓ | ✓ | ✓ | | |
| ENSORLG00000013692 | p3h4 | L260I | ✓ | ✓ | ✓ | | |
| ENSXMAG00000018830 | pkd1l1 | T1282S | ✓ | ✓ | ✓ | | |
| ENSORLG00000000363 | gpr171 | A285E | ✓ | ✓ | ✓ | | |
| ENSORLG00000006625 | slc39a6 | M615T | ✓ | ✓ | ✓ | | |
| ENSORLG00000013531 | mc4r | V99A | ✓ | ✓ | ✓ | | |
| ENSORLG00000012478 | trpm6 | R489C | ✓ | ✓ | ✓ | | |
| ENSORLG00000000363 | gpr171 | A285E | ✓ | ✓ | ✓ | | |
| ENSORLG00000010311 | slc27a4 | S414T | ✓ | ✓ | ✓ | | |
| ENSORLG00000020785 | pde5a | K726G | ✓ | ✓ | ✓ | | |
| ENSORLG00000005152 | enc3 | T182K | ✓ | ✓ | ✓ | | |
| ENSORLG00000015350 | clip2 | N895S | ✓ | ✓ | ✓ | | |
| ENSORLG00000008035 | chrne | I236T | ✓ | ✓ | ✓ | | |
| ENSORLG00000002046 | coro1a | A384G | ✓ | ✓ | ✓ | | |
| ENSORLG00000011056 | wasf2 | L126I | ✓ | ✓ | ✓ | | |

** present in >70% of sequenced flying fishes (Exocoetidae)

#### S11 Table. Fin and limb genes

| Ensembl ID | gene name | fin screens | fin/limb GO† | avg. coverage |
|---|---|---|---|---|
| ENSORLG00000010229 | kcnj13 | ✓ | | 93.2% |
| ENSORLG00000009715 | lat4a | ✓ | | 92.1% |
| ENSORLG00000006810 | kcnk5b | ✓ | ✓ | 79.0% |
| ENSORLG00000012908 | kcc4a | ✓ | | 80.5% |
| ENSORLG00000004137 | kcnh2b | ✓ | | 84.1% |
| ENSORLG00000010176 | aqp3a | ✓ | | 76.7% |
| ENSORLG00000010431 | col9a1 | ✓ | | 63.5% |
| ENSORLG00000028143 | col9a1 | ✓ | | 63.5% |
| ENSORLG00000015485 | cx43 | ✓ | ✓ | 95.9% |
| ENSORLG00000017788 | kcnk5a | ✓ | | 93.5% |
| ENSORLG00000004461 | slc12a7b | ✓ | | 86.5% |
| ENSORLG00000005019 | lat4b | ✓ | | 95.3% |
| ENSORLG00000015086 | kcnj1b | ✓ | | 96.6% |
| ENSORLG00000024246 | kcnj10a | ✓ | | 96.6% |
| ENSORLG00000013245 | kcnk9 | ✓ | | 99.9% |
| ENSORLG00000000299 | kremen1 | | ✓ | 93.5% |
| ENSORLG00000010075 | grip2a | | ✓ | 88.8% |
| ENSORLG00000006966 | NA | | ✓ | 17.6% |
| ENSORLG00000005789 | pcsk5b | | ✓ | 95.5% |
| ENSORLG00000019737 | met | | ✓ | 79.1% |
| ENSORLG00000007687 | plxna2 | | ✓ | 93.4% |
| ENSORLG00000022179 | psen1 | | ✓ | 93.4% |
| ENSORLG00000007960 | sox9 | | ✓ | 94.1% |
| ENSORLG00000006013 | fgf1a | | ✓ | 87.7% |
| ENSORLG00000022701 | tbx5a | | ✓ | 87.7% |
| ENSORLG00000018545 | lrp5 | | ✓ | 91.3% |
| ENSORLG00000029196 | lmo2 | | ✓ | 91.3% |
| ENSORLG00000010734 | osr1 | | ✓ | 99.3% |
| ENSORLG00000010531 | lmbr1 | | ✓ | 90.3% |
| ENSORLG00000010536 | scfd1 | | ✓ | 92.0% |
| ENSORLG00000015015 | id2a | | ✓ | 100.0% |
| ENSORLG00000006545 | ext2 | | ✓ | 94.2% |
| ENSORLG00000001350 | krt8l | | ✓ | 44.0% |
| ENSORLG00000029413 | fgf10b | | ✓ | 44.0% |
| ENSORLG00000016082 | sema3c | | ✓ | 93.5% |
| ENSORLG00000007361 | shox2 | | ✓ | 99.7% |
| ENSORLG00000015897 | edar | | ✓ | 81.8% |

| | | | |
|---|---|---|---|
| ENSORLG00000014420 | hmx4 | ✓ | 8.1% |
| ENSORLG00000002662 | alx1 | ✓ | 98.3% |
| ENSORLG00000001210 | znrf3 | ✓ | 99.7% |
| ENSORLG00000005925 | mbnL1 | ✓ | 96.4% |
| ENSORLG00000010960 | galnt2 | ✓ | 95.8% |
| ENSORLG00000002708 | yap1 | ✓ | 95.4% |
| ENSORLG00000010102 | fkbp1b | ✓ | 86.5% |
| ENSORLG00000007009 | NA | ✓ | 28.6% |
| ENSORLG00000010725 | dync2h1 | ✓ | 65.6% |
| ENSORLG00000000906 | fbxw4 | ✓ | 49.6% |
| ENSORLG00000018971 | hnf1a | ✓ | 96.7% |
| ENSORLG00000016099 | ptn | ✓ | 92.6% |
| ENSORLG00000022704 | si:dkey-182i3.8 | ✓ | 92.6% |
| ENSORLG00000022160 | asph | ✓ | 92.6% |
| ENSORLG00000020705 | rspo2 | ✓ | 99.6% |
| ENSORLG00000024896 | XlCGF57.1 | ✓ | 99.6% |
| ENSORLG00000020663 | frem1a | ✓ | 67.0% |
| ENSORLG00000005157 | irx1a | ✓ | 99.9% |
| ENSORLG00000009600 | hmcn1 | ✓ | 84.6% |
| ENSORLG00000008813 | wnt5a | ✓ | 99.4% |
| ENSORLG00000001366 | lnpk | ✓ | 58.8% |
| ENSORLG00000005702 | erbb2 | ✓ | 75.4% |
| ENSORLG00000012238 | kat2b | ✓ | 86.7% |
| ENSORLG00000029206 | hapln1a | ✓ | 86.7% |
| ENSORLG00000000073 | cdh2 | ✓ | 9.0% |
| ENSORLG00000001615 | si:dkey-182i3.9 | ✓ | 76.0% |
| ENSORLG00000013530 | megf8 | ✓ | 90.0% |
| ENSORLG00000010637 | hspa9 | ✓ | 57.4% |
| ENSORLG00000005063 | ARL6 | ✓ | 89.6% |
| ENSORLG00000013827 | sall1a | ✓ | 94.9% |
| ENSORLG00000007425 | ift140 | ✓ | 76.7% |
| ENSORLG00000008429 | nr2f2 | ✓ | 100.0% |
| ENSORLG00000029633 | sox4a | ✓ | 100.0% |
| ENSORLG00000013436 | col10a1a | ✓ | 94.4% |
| ENSORLG00000017878 | grem1a | ✓ | 89.0% |
| ENSORLG00000011080 | ctsba | ✓ | 81.0% |
| ENSORLG00000005880 | wnt2ba | ✓ | 84.8% |
| ENSORLG00000006191 | alx3 | ✓ | 99.7% |
| ENSORLG00000022243 | c2cd3 | ✓ | 99.7% |
| ENSORLG00000002596 | bmpr1aa | ✓ | 93.4% |

| | | | |
|---|---|---|---|
| ENSORLG00000030227 | fras1 | ✓ | 93.4% |
| ENSORLG00000000135 | ahr2 | ✓ | 59.9% |
| ENSORLG00000012027 | wnt3 | ✓ | 98.9% |
| ENSORLG00000027472 | acd | ✓ | 98.9% |
| ENSORLG00000004625 | cul4a | ✓ | 96.1% |
| ENSORLG00000002307 | NA | ✓ | 24.4% |
| ENSORLG00000018020 | dll4 | ✓ | 74.7% |
| ENSORLG00000000246 | camk2b1 | ✓ | 46.8% |
| ENSORLG00000009978 | smoc1 | ✓ | 69.0% |
| ENSORLG00000003744 | rnf165b | ✓ | 97.9% |
| ENSORLG00000005532 | wdr19 | ✓ | 81.8% |
| ENSORLG00000015665 | tp63 | ✓ | 91.5% |
| ENSORLG00000029983 | gdf5 | ✓ | 91.5% |
| ENSORLG00000003465 | cyp26b1 | ✓ | 99.4% |
| ENSORLG00000025793 | inhbaa | ✓ | 99.4% |
| ENSORLG00000014081 | rpgrip1l | ✓ | 44.1% |
| ENSORLG00000024353 | has2 | ✓ | 44.1% |
| ENSORLG00000000976 | tcf7 | ✓ | 83.9% |
| ENSORLG00000017084 | cdh2 | ✓ | 89.9% |
| ENSORLG00000012390 | znf709 | ✓ | 77.9% |
| ENSORLG00000004542 | dlx6a | ✓ | 97.2% |
| ENSORLG00000002036 | cyp26c1 | ✓ | 93.6% |
| ENSORLG00000015772 | plxna2 | ✓ | 95.0% |
| ENSORLG00000020874 | hspd1 | ✓ | 97.7% |
| ENSORLG00000013834 | acvr1l | ✓ | 81.6% |
| ENSORLG00000007341 | hhip | ✓ | 94.6% |
| ENSORLG00000000900 | camk2b | ✓ | 67.3% |
| ENSORLG00000014155 | hdac1 | ✓ | 93.3% |
| ENSORLG00000008185 | tbx5 | ✓ | 87.2% |
| ENSORLG00000026489 | chsy1 | ✓ | 87.2% |
| ENSORLG00000020505 | flvcr1 | ✓ | 83.4% |
| ENSORLG00000015684 | prdm1 | ✓ | 88.9% |
| ENSORLG00000013440 | prrx1a | ✓ | 94.2% |
| ENSORLG00000008319 | aldh1a2 | ✓ | 98.2% |
| ENSORLG00000014640 | zbtb16 | ✓ | 100.0% |
| ENSORLG00000012738 | col2a1a | ✓ | 93.2% |
| ENSORLG00000014312 | serpinh1b | ✓ | 95.4% |
| ENSORLG00000011720 | bmp7b | ✓ | 92.7% |
| ENSORLG00000027985 | fgf10a | ✓ | 92.7% |
| ENSORLG00000017013 | col1 | ✓ | 89.8% |

| | | | |
|---|---|---|---|
| ENSORLG00000007606 | nipbl | ✓ | 76.6% |
| ENSORLG00000017092 | rdh1 | ✓ | 86.0% |
| ENSORLG00000007913 | hoxc10a | ✓ | 99.6% |
| ENSORLG00000024737 | lrp6 | ✓ | 99.6% |
| ENSORLG00000029931 | si:ch73-281k2.5 | ✓ | 99.6% |
| ENSORLG00000017491 | hdac1 | ✓ | 93.4% |
| ENSORLG00000014806 | tbx4 | ✓ | 79.9% |
| ENSORLG00000017492 | sp9 | ✓ | 99.7% |
| ENSORLG00000023821 | znf516 | ✓ | 99.7% |
| ENSORLG00000010277 | crabp2a | ✓ | 99.5% |
| ENSORLG00000012857 | fgf8b | ✓ | 77.4% |
| ENSORLG00000007858 | comp | ✓ | 90.2% |
| ENSORLG00000004209 | itga3b | ✓ | 48.2% |
| ENSORLG00000023950 | pik4aa | ✓ | 48.2% |
| ENSORLG00000013492 | chd7 | ✓ | 91.7% |
| ENSORLG00000005031 | rnf165a | ✓ | 99.2% |
| ENSORLG00000011261 | wnt7ab | ✓ | 99.7% |
| ENSORLG00000015787 | atp6v1e1b | ✓ | 88.1% |
| ENSORLG00000012324 | fam92a1 | ✓ | 62.5% |
| ENSORLG00000024876 | gja5b | ✓ | 62.5% |
| ENSORLG00000007685 | erbb3b | ✓ | 80.6% |
| ENSORLG00000005722 | gnas | ✓ | 89.7% |
| ENSORLG00000012987 | tubb5 | ✓ | 96.1% |
| ENSORLG00000013512 | med31 | ✓ | 90.2% |
| ENSORLG00000010018 | rc3h2 | ✓ | 83.7% |
| ENSORLG00000007861 | rarga | ✓ | 99.9% |
| ENSORLG00000029523 | bmpr2 | ✓ | 99.9% |
| ENSORLG00000025597 | pi4ka | ✓ | 99.9% |
| ENSORLG00000011156 | bmp1a | ✓ | 74.4% |
| ENSORLG00000009187 | atrx | ✓ | 55.5% |
| ENSORLG00000008300 | med1 | ✓ | 89.6% |
| ENSORLG00000003661 | gnas | ✓ | 91.6% |
| ENSORLG00000013277 | fgfr2 | ✓ | 95.5% |
| ENSORLG00000022089 | aff3 | ✓ | 95.5% |
| ENSORLG00000011187 | prdm9 | ✓ | 74.8% |
| ENSORLG00000026863 | igf2b | ✓ | 74.8% |
| ENSORLG00000006633 | tfap2a | ✓ | 99.5% |
| ENSORLG00000000929 | nog1 | ✓ | 89.8% |
| ENSORLG00000006924 | lef1 | ✓ | 85.6% |
| ENSORLG00000019677 | lama5 | ✓ | 54.8% |

| | | | |
|---|---|---|---|
| ENSORLG00000018904 | plod1a | ✓ | 69.3% |
| ENSORLG00000008008 | tll1 | ✓ | 94.1% |
| ENSORLG00000001349 | hoxd9 | ✓ | 92.3% |
| ENSORLG00000017695 | tfap2b | ✓ | 94.0% |
| ENSORLG00000002333 | slc39a3 | ✓ | 99.6% |
| ENSORLG00000005102 | col2a1a | ✓ | 85.8% |
| ENSORLG00000007833 | bmpr1bb | ✓ | 84.9% |
| ENSORLG00000014423 | map3k20 | ✓ | 96.6% |
| ENSORLG00000004596 | lfng | ✓ | 81.8% |
| ENSORLG00000017140 | faf1 | ✓ | 92.0% |
| ENSORLG00000003890 | nid2a | ✓ | 64.1% |
| ENSORLG00000010866 | mog-12 | ✓ | 100.0% |
| ENSORLG00000023692 | dkk1 | ✓ | 100.0% |
| ENSORLG00000011685 | sox2 | ✓ | 100.0% |
| ENSORLG00000020123 | sec23a | ✓ | 93.7% |
| ENSORLG00000016394 | RARB | ✓ | 97.4% |
| ENSORLG00000010470 | tnf | ✓ | 51.9% |
| ENSORLG00000013172 | and1 | ✓ | 85.5% |
| ENSORLG00000006340 | mmp9 | ✓ | 73.9% |
| ENSORLG00000001319 | cldnc | ✓ | 95.0% |
| ENSORLG00000007608 | hoxa11b | ✓ | 97.7% |
| ENSORLG00000030564 | kctd10 | ✓ | 97.7% |
| ENSORLG00000011562 | extl3 | ✓ | 99.6% |
| ENSORLG00000010166 | slc7a2 | ✓ | 90.7% |
| ENSORLG00000008648 | ift172 | ✓ | 44.0% |
| ENSORLG00000005453 | hes4 | ✓ | 99.4% |
| ENSORLG00000028091 | XlCGF57.1-like | ✓ | 99.4% |
| ENSORLG00000001499 | fgf1b | ✓ | 97.6% |
| ENSORLG00000003955 | smarca4 | ✓ | 88.5% |
| ENSORLG00000013379 | hmcn2 | ✓ | 63.3% |
| ENSORLG00000005037 | hoxa11a | ✓ | 100.0% |
| ENSORLG00000006075 | atrx | ✓ | 72.5% |
| ENSORLG00000020718 | sall3a | ✓ | 99.4% |
| ENSORLG00000020852 | etv5a | ✓ | 78.7% |
| ENSORLG00000026886 | prdx5 | ✓ | 78.7% |
| ENSORLG00000027241 | a-sdf1a | ✓ | 78.7% |
| ENSORLG00000005359 | rnf2 | ✓ | 94.5% |
| ENSORLG00000013366 | col22a1 | ✓ | 58.6% |
| ENSORLG00000016129 | zmp:0000000711 | ✓ | 67.5% |
| ENSORLG00000025622 | esco2 | ✓ | 67.5% |

| | | | |
|---|---|---|---|
| ENSORLG00000025148 | kdf1b | ✓ | 67.5% |
| ENSORLG00000020806 | fgf20b | ✓ | 88.8% |
| ENSORLG00000004105 | frem2a | ✓ | 83.6% |
| ENSORLG00000023719 | rspo3 | ✓ | 83.6% |
| ENSORLG00000005443 | mtrex | ✓ | 83.2% |
| ENSORLG00000013805 | hs6st1b | ✓ | 98.2% |
| ENSORLG00000017600 | wnt9a | ✓ | 99.3% |
| ENSORLG00000020587 | pitx2 | ✓ | 93.9% |
| ENSORLG00000005446 | nipbla | ✓ | 94.1% |
| ENSORLG00000013972 | ep300b | ✓ | 89.3% |
| ENSORLG00000008502 | rarb | ✓ | 99.4% |
| ENSORLG00000005565 | bmpr1ba | ✓ | 94.5% |
| ENSORLG00000005755 | rspo3 | ✓ | 88.7% |
| ENSORLG00000010463 | shha | ✓ | 100.0% |
| ENSORLG00000017531 | lnpa | ✓ | 73.9% |
| ENSORLG00000017540 | hoxd9 | ✓ | 88.8% |
| ENSORLG00000009669 | fam53b | ✓ | 89.9% |
| ENSORLG00000005845 | ctnnb1 | ✓ | 98.8% |
| ENSORLG00000005182 | notch1a | ✓ | 96.7% |
| ENSORLG00000006087 | fgf16 | ✓ | 98.6% |
| ENSORLG00000017936 | and2 | ✓ | 88.9% |
| ENSORLG00000009380 | plcg1 | ✓ | 90.5% |
| ENSORLG00000010264 | kdm6bb | ✓ | 72.4% |
| ENSORLG00000027052 | fgf21 | ✓ | 72.4% |
| ENSORLG00000006310 | chd4 | ✓ | 96.2% |
| ENSORLG00000001551 | acanb | ✓ | 71.9% |
| ENSORLG00000007097 | NA | ✓ | 22.8% |
| ENSORLG00000030115 | alx4a | ✓ | 22.8% |
| ENSORLG00000026349 | nrg1 | ✓ | 22.8% |
| ENSORLG00000030507 | mbnl1 | ✓ | 22.8% |
| ENSORLG00000002188 | hs6st2 | ✓ | 87.1% |
| ENSORLG00000012077 | mecom | ✓ | 94.4% |
| ENSORLG00000014296 | cacna1c | ✓ | 97.1% |
| ENSORLG00000024917 | fgf8a | ✓ | 97.1% |
| ENSORLG00000015382 | rargb | ✓ | 99.3% |
| ENSORLG00000002493 | vdac3 | ✓ | 80.0% |
| ENSORLG00000029777 | etv5b | ✓ | 80.0% |
| ENSORLG00000017122 | gata3 | ✓ | 88.8% |
| ENSORLG00000024149 | hoxc8 | ✓ | 88.8% |
| ENSORLG00000002496 | ep300a | ✓ | 84.4% |

| | | | |
|---|---|---|---|
| ENSORLG00000012381 | anxa1 | ✓ | 92.1% |
| ENSORLG00000026325 | pam | ✓ | 92.1% |
| ENSORLG00000015588 | arl6 | ✓ | 82.3% |
| ENSORLG00000007884 | intu | ✓ | 86.0% |
| ENSORLG00000016130 | sall4 | ✓ | 81.0% |
| ENSORLG00000000321 | fgfr1a | ✓ | 68.1% |
| ENSORLG00000013881 | foxl1 | ✓ | 70.3% |
| ENSORLG00000005001 | sox4b | ✓ | 95.7% |
| ENSORLG00000025013 | wnt7aa | ✓ | 95.7% |
| ENSORLG00000013494 | bmp7a | ✓ | 78.2% |
| ENSORLG00000006068 | hoxa9 | ✓ | 78.4% |
| ENSORLG00000007983 | ap1g1 | ✓ | 96.8% |
| ENSORLG00000005764 | aldh7a1 | ✓ | 99.8% |
| ENSORLG00000007667 | wnt8a | ✓ | 87.0% |
| ENSORLG00000022857 | mustn1a | ✓ | 87.0% |
| ENSORLG00000025251 | kdf1a | ✓ | 87.0% |
| ENSORLG00000010756 | ccn1 | ✓ | 92.5% |
| ENSORLG00000005573 | YAP1 | ✓ | 74.8% |
| ENSORLG00000024546 | meox2b | ✓ | 74.8% |
| ENSORLG00000004345 | ptch1 | ✓ | 97.7% |
| ENSORLG00000012260 | tnfrsf1a | ✓ | 30.2% |
| ENSORLG00000017534 | evx2 | ✓ | 100.0% |
| ENSORLG00000003868 | kat2a | ✓ | 95.9% |
| ENSORLG00000005050 | evx1 | ✓ | 91.6% |
| ENSORLG00000014206 | fgfr1 | ✓ | 91.8% |
| ENSORLG00000007986 | rdh10a | ✓ | 99.9% |
| ENSORLG00000017925 | gja5a | ✓ | 98.7% |
| ENSORLG00000009822 | gpc3 | ✓ | 93.5% |
| ENSORLG00000008547 | sufu | ✓ | 96.1% |
| ENSORLG00000003034 | bmpr2a | ✓ | 91.6% |
| ENSORLG00000000636 | notch1b | ✓ | 93.9% |
| ENSORLG00000014357 | smo | ✓ | 96.9% |
| ENSORLG00000015452 | smarca4a | ✓ | 95.4% |
| ENSORLG00000014996 | sox11a | ✓ | 94.2% |
| ENSORLG00000011953 | chst11 | ✓ | 83.1% |
| ENSORLG00000013304 | bmp4 | ✓ | 99.9% |
| ENSORLG00000000217 | NA | ✓ | 74.3% |
| ENSORLG00000007763 | mustn1b | ✓ | 84.4% |
| ENSORLG00000004731 | alx4b | ✓ | 99.6% |
| ENSORLG00000025920 | dhrs9 | ✓ | 99.6% |

| | | | |
|---|---|---|---|
| ENSORLG00000004568 | baxa | ✓ | 36.4% |
| ENSORLG00000011039 | IFT122 | ✓ | 88.1% |
| ENSORLG00000020070 | bbs7 | ✓ | 86.2% |
| ENSORLG00000017597 | wnt3a | ✓ | 87.4% |
| ENSORLG00000004561 | dlx5a | ✓ | 99.9% |
| ENSORLG00000022655 | apc | ✓ | 99.9% |
| ENSORLG00000025714 | sec23a | ✓ | 99.9% |
| ENSORLG00000028686 | agr1 | ✓ | 99.9% |
| ENSORLG00000009133 | furina | ✓ | 89.5% |
| ENSORLG00000029979 | nr2f2 | ✓ | 89.5% |
| ENSORLG00000009629 | iqce | ✓ | 53.8% |
| ENSORLG00000008532 | igf1ra | ✓ | 85.2% |
| ENSORLG00000016981 | NA | ✓ | 66.3% |
| ENSORLG00000007566 | acvr1b | ✓ | 90.9% |
| ENSORLG00000020379 | bbs1 | ✓ | 97.9% |
| ENSORLG00000006552 | vdac2 | ✓ | 92.8% |
| ENSORLG00000002797 | osr2 | ✓ | 99.9% |
| ENSORLG00000013709 | wnt5b | ✓ | 95.7% |
| ENSORLG00000016855 | skia | ✓ | 86.9% |
| ENSORLG00000000295 | fgf4 | ✓ | 94.1% |
| ENSORLG00000007242 | sp8b | ✓ | 96.0% |
| ENSORLG00000016908 | grem1 | ✓ | 95.2% |
| ENSORLG00000007648 | wnt8a | ✓ | 90.6% |
| ENSORLG00000010117 | NA | ✓ | 97.9% |
| ENSORLG00000021995 | junbb | ✓ | 97.9% |
| ENSORLG00000025844 | crabp2b | ✓ | 97.9% |
| ENSORLG00000011799 | mta2 | ✓ | 84.0% |
| ENSORLG00000004154 | notch2 | ✓ | 63.7% |
| ENSORLG00000007316 | tbx3a | ✓ | 99.7% |
| ENSORLG00000012490 | gli3 | ✓ | 87.5% |
| ENSORLG00000014732 | igf1rb | ✓ | 94.2% |
| ENSORLG00000000356 | lamb1a | ✓ | 80.2% |
| ENSORLG00000016905 | fmn1 | ✓ | 84.4% |

† included fin/limb related GO terms:
GO:0035108,GO:0035143,GO:0035141,GO:0060173,GO:0060174,GO:0035122,GO:0035125,GO:0035124,GO:0035128,GO:0030326,GO:0033334,GO:0033333,GO:0033336,GO:0033337,GO:0031101,GO:0033335,GO:0033338,GO:0033339,GO:0035115,GO:0035116,GO:0035136,GO:0035137,GO:0060887,GO:0035138,GO:0035118,GO:0035119,GO:0033340

#### S12 Table. Convergent relative evolutionary rate across key fin and limb genes

| Ensembl ID | gene name | rho | p-value | q-value |
|---|---|---|---|---|
| ENSORLG00000005019 | slc43a2b | 0.32 | 6.54E-03 | 2.53E-01 |
| ENSORLG00000010229 | kcnj13 | 0.26 | 3.42E-02 | 4.23E-01 |
| ENSORLG00000007687 | plxna2 | 0.57 | 3.52E-07 | 2.10E-03 |
| ENSORLG00000004105 | frem2a | 0.47 | 5.68E-05 | 3.33E-02 |
| ENSORLG00000013492 | chd7 | -0.40 | 6.16E-04 | 9.12E-02 |
| ENSORLG00000015684 | prdm1 | 0.36 | 2.21E-03 | 1.66E-01 |
| ENSORLG00000011685 | sox2 | 0.36 | 2.26E-03 | 1.69E-01 |
| ENSORLG00000010866 | mog-12 | 0.36 | 2.37E-03 | 1.71E-01 |
| ENSORLG00000010018 | rc3h2 | -0.34 | 4.41E-03 | 2.18E-01 |
| ENSORLG00000016394 | rarb | 0.34 | 4.87E-03 | 2.28E-01 |
| ENSORLG00000005182 | notch1a | 0.33 | 5.51E-03 | 2.41E-01 |
| ENSORLG00000011039 | ift122 | -0.33 | 5.84E-03 | 2.45E-01 |
| ENSORLG00000013827 | sall1a | 0.33 | 5.95E-03 | 2.48E-01 |
| ENSORLG00000004542 | dlx6a | 0.33 | 6.24E-03 | 2.48E-01 |
| ENSORLG00000005050 | evx1 | 0.32 | 8.34E-03 | 2.71E-01 |
| ENSORLG00000020587 | pitx2 | 0.31 | 8.49E-03 | 2.71E-01 |
| ENSORLG00000020123 | sec23a | 0.31 | 8.91E-03 | 2.73E-01 |
| ENSORLG00000008547 | sufu | -0.31 | 9.59E-03 | 2.79E-01 |
| ENSORLG00000013440 | prrx1a | 0.31 | 1.04E-02 | 2.81E-01 |
| ENSORLG00000013436 | col10a1a | 0.28 | 1.99E-02 | 3.55E-01 |
| ENSORLG00000007361 | shox2 | 0.28 | 2.08E-02 | 3.61E-01 |
| ENSORLG00000003868 | kat2a | 0.27 | 2.44E-02 | 3.82E-01 |
| ENSORLG00000000929 | nog1 | 0.27 | 2.51E-02 | 3.84E-01 |
| ENSORLG00000016905 | fmn1 | -0.27 | 2.66E-02 | 3.92E-01 |
| ENSORLG00000007316 | tbx3a | 0.27 | 2.73E-02 | 3.95E-01 |
| ENSORLG00000014732 | igf1rb | 0.25 | 4.08E-02 | 4.47E-01 |
| ENSORLG00000014640 | zbtb16 | 0.25 | 4.14E-02 | 4.47E-01 |
| ENSORLG00000015452 | smarca4a | 0.24 | 4.35E-02 | 4.53E-01 |
| ENSORLG00000009133 | furina | -0.24 | 4.52E-02 | 4.60E-01 |
| ENSORLG00000008319 | aldh1a2 | -0.24 | 4.58E-02 | 4.62E-01 |
| ENSORLG00000012027 | wnt3 | 0.24 | 4.98E-02 | 4.71E-01 |
| ENSORLG00000017013 | col1 | 0.24 | 4.99E-02 | 4.71E-01 |

#### S13 Table. Accelerated or constrained sequence evolution across key fin and limb genes at the ancestral node of flying fishes (Exocoetidae)

| Ensembl ID | gene name | cons/acc | phyloP pvalue |
|---|---|---|---|
| ENSORLG00000015485 | cx43 | con | 0.000000 |
| ENSORLG00000005019 | slc43a2b | acc | 0.000010 |
| ENSORLG00000013245 | kcnk9 | acc | 0.000170 |
| ENSORLG00000017788 | kcnk5a | acc | 0.002350 |
| ENSORLG00000009715 | slc43a2a | acc | 0.008370 |
| ENSORLG00000003465 | cyp26b1 | acc | 0.000000 |
| ENSORLG00000004105 | frem2a | acc | 0.000000 |
| ENSORLG00000007687 | plxna2 (1 of 3) | acc | 0.000000 |
| ENSORLG00000008429 | nr2f2 | acc | 0.000000 |
| ENSORLG00000013827 | sall1a | acc | 0.000000 |
| ENSORLG00000014155 | hdac1 (1 of 2) | acc | 0.000000 |
| ENSORLG00000014640 | zbtb16a (2 of 2) | acc | 0.000000 |
| ENSORLG00000018904 | plod1a | acc | 0.000000 |
| ENSORLG00000003868 | kat2a | acc | 0.000020 |
| ENSORLG00000016981 | NA | con | 0.000030 |
| ENSORLG00000002307 | si:dkey-261j4.4 | con | 0.000060 |
| ENSORLG00000004154 | notch2 | con | 0.000070 |
| ENSORLG00000000929 | nog1 | acc | 0.000080 |
| ENSORLG00000006633 | tfap2a | acc | 0.000100 |
| ENSORLG00000001551 | acanb | con | 0.000150 |
| ENSORLG00000020379 | bbs1 | con | 0.000590 |
| ENSORLG00000002662 | alx1 | con | 0.000670 |
| ENSORLG00000001349 | hoxd9b | acc | 0.001270 |
| ENSORLG00000004542 | dlx6a | acc | 0.001270 |
| ENSORLG00000016394 | NA | acc | 0.001490 |
| ENSORLG00000011685 | sox2 | acc | 0.001760 |
| ENSORLG00000017013 | col1 | acc | 0.001860 |
| ENSORLG00000015382 | rargb | con | 0.002060 |
| ENSORLG00000018020 | dll4 | con | 0.002120 |
| ENSORLG00000009600 | hmcn1 | acc | 0.002570 |
| ENSORLG00000000295 | fgf4 | acc | 0.003200 |
| ENSORLG00000002708 | yap1 | con | 0.004180 |
| ENSORLG00000001615 | zgc:171570 | acc | 0.004560 |
| ENSORLG00000012390 | si:dkey-14k9.3 | acc | 0.004640 |
| ENSORLG00000017878 | grem1a | con | 0.004950 |
| ENSORLG00000005001 | sox4 | acc | 0.006430 |

| | | | |
|---|---|---|---|
| ENSORLG00000009380 | plcg1 | con | 0.006430 |
| ENSORLG00000013277 | fgfr2 | con | 0.007720 |
| ENSORLG00000005050 | evx1 | acc | 0.008510 |
| ENSORLG00000001350 | krt5 | con | 0.008960 |
| ENSORLG00000002036 | cyp26c1 | con | 0.009130 |
| ENSORLG00000010756 | cyr61 | acc | 0.009350 |
| ENSORLG00000018545 | lrp5 | con | 0.009420 |
| ENSORLG00000003034 | bmpr2a | acc | 0.009710 |
| ENSORLG00000014732 | igf1ra | acc | 0.009740 |
| ENSORLG00000007242 | sp8b | con | 0.010220 |
| ENSORLG00000001210 | znrf3 | con | 0.010470 |
| ENSORLG00000006310 | chd4a | con | 0.011210 |
| ENSORLG00000014357 | smo | con | 0.011650 |
| ENSORLG00000016908 | grem1 (1 of 2) | con | 0.012340 |
| ENSORLG00000002333 | slc39a3 | con | 0.014960 |
| ENSORLG00000013494 | bmp7a | con | 0.015610 |
| ENSORLG00000016855 | skia | con | 0.016170 |
| ENSORLG00000004625 | cul4a | acc | 0.021280 |
| ENSORLG00000017092 | rdh1 | acc | 0.022440 |
| ENSORLG00000005755 | rspo3 | con | 0.024160 |
| ENSORLG00000001366 | lnpb | acc | 0.024580 |
| ENSORLG00000000135 | ahr2 (1 of 2) | acc | 0.025280 |
| ENSORLG00000010725 | dync2h1 | con | 0.026010 |
| ENSORLG00000003955 | smarca4 (1 of 2) | acc | 0.027050 |
| ENSORLG00000020123 | sec23a | con | 0.029330 |
| ENSORLG00000013492 | chd7 | con | 0.029500 |
| ENSORLG00000003890 | nid2a | con | 0.030610 |
| ENSORLG00000007648 | wnt8a | acc | 0.034900 |
| ENSORLG00000005182 | notch1a | acc | 0.036250 |
| ENSORLG00000003744 | rnf165b | con | 0.036330 |
| ENSORLG00000014081 | rpgrip1l | con | 0.040010 |
| ENSORLG00000014996 | sox11 | con | 0.040670 |
| ENSORLG00000020587 | pitx2 | con | 0.041670 |
| ENSORLG00000002188 | hs6st2 | con | 0.043640 |
| ENSORLG00000002797 | osr2 | con | 0.045290 |
| ENSORLG00000005157 | irx1a | con | 0.045660 |
| ENSORLG00000012987 | tubb5 | con | 0.048130 |
| ENSORLG00000012260 | tnfrsf1a | acc | 0.048620 |
| ENSORLG00000011080 | ctsba | acc | 0.049640 |
| ENSORLG00000002596 | bmpr1aa | con | 0.049920 |
